# Sex Disparities in Gastric Cancer Tumor Mutation Burden and Gene Mutation Patterns

**DOI:** 10.1101/2025.05.24.655976

**Authors:** Noa Klein, Dorit Shweiki

## Abstract

Significant sex-related differences exist in gastric cancer, yet the molecular basis remains unclear. Tumor mutational burden (TMB) correlates with successful immune checkpoint inhibitor therapy, but TMB varies across different cancers, and evidence regarding sex-specific differences remains inconsistent. We analyzed sex-specific single-nucleotide mutations in gastric cancer using The Cancer Genome Atlas Stomach Adenocarcinoma (TCGA-STAD) dataset and found that females had significantly higher autosomal TMB. We identified 1,052 sexually dimorphic genes, with enrichment analyses highlighting distinct biological pathways in each sex. These genes overlap key cancer pathways, emphasizing their role in tumorigenesis. Differing co-mutation patterns—with females exhibiting more stable patterns—enabled a machine-learning model to predict tumor sex from mutation profiles. Our findings suggest that male and female gastric cancers could represent distinct diseases, underscoring the need for sex-specific diagnostic markers and personalized therapies.

## Background

Gastrointestinal cancers account for a quarter of cancer incidence and a third of cancer-related deaths worldwide. Significant geographical, socioeconomic, age, and sex-related differences are evident across all gastrointestinal cancers [1, 2].

Among these, gastric cancer (GC) is one of the most common and fatal malignancies. It ranks as the fifth most common cancer and the fourth leading cause of cancer-associated mortality, accounting for 7.7% of all cancer deaths [3, 4]. Globally, the incidence and mortality rates are approximately two to three times higher in men than in women. In 2020, the global incidence rate in men was 15.8 per 100,000 compared to 7.0 per 100,000 in women. Notable variations are observed worldwide: in the United States in 2020, the incident rate was 8.5 per 100,00 in men and 5.8 per 100,000 in women, while in Japan, the rates were as high as 48.1 per 100,000 in men and 17.3 per 100,000 in women [3, 5].

High tumor mutational burden (TMB) is considered a predictive biomarker for the success of immune checkpoint inhibitors (ICIs) therapy and has been approved by the FDA as an indicator for pembrolizumab treatment [6]. Consequently, the behavior of TMB as a valid biomarker has attracted considerable attention. Sex-related differences in immunotherapeutic outcomes and their correlation with TMB values have been widely studied; however, the reports vary between cancer types and studies. In some pan-cancer studies [7, 8] and study focusing on bladder cancer [9], males exhibited better immunotherapy outcomes correlating with higher TMB. Conversely, better immunotherapy outcomes and higher TMB values were reported for female lung cancer patients [10]. Furthermore, a meta-analysis of twenty-three clinical trials found no significant correlation between patient sex and the effectiveness of immunotherapy in advanced cancer treatment [11].

Reports on sex disparities in TMB values are inconsistent. While numerous studies have reported higher TMB values in male cancers [7, 12, 13], others have found higher TMB values in females [14], and some have observed no significant differences between the sexes [15, 16]. Additionally, differences in TMB values have been shown to correlate with age [14] and ethnicity [16]. These inconsistencies are primarily attributed to the use of dissimilar gene panels and whether TMB values are calculated for all chromosomes or solely for autosomes. This underscores the need for accurate comparisons that consider all relevant parameters.

In this study, we conducted a comprehensive analysis of sex disparities in single-nucleotide mutations in GC. By demonstrating sex-specific molecular patterns, a deeper understanding is achieved, which can significantly advance diagnostics and personalized medical treatments. Here, we report distinct genetic profiles of male and female patients as identified in The Cancer Genome Atlas Stomach Adenocarcinoma (TCGA-STAD) dataset, potentially guiding the development of patient-specific therapeutic approaches.

## Results

### Sexually Dimorphic Autosomal Tumor Mutation Burden in TCGA Gastrointestinal Samples

Autosomal TMB values were calculated for the five gastric system projects based on data generated by the TCGA Research Network. TMB values were analyzed for significant differences between males and females in each project. The stomach adenocarcinoma project (STAD) showed significant differences in TMB values between males and females (corrected p-value 0.039036, Fig. 1A), while the colon adenocarcinoma (COAD), rectum adenocarcinoma (READ), hepatocellular carcinoma (LIHC), and pancreatic adenocarcinoma (PAAD) projects showed no significant differences between the sexes (the esophageal carcinoma, (ESCA) project was not included due to the small number of women’s samples included).

**Fig. 1:**
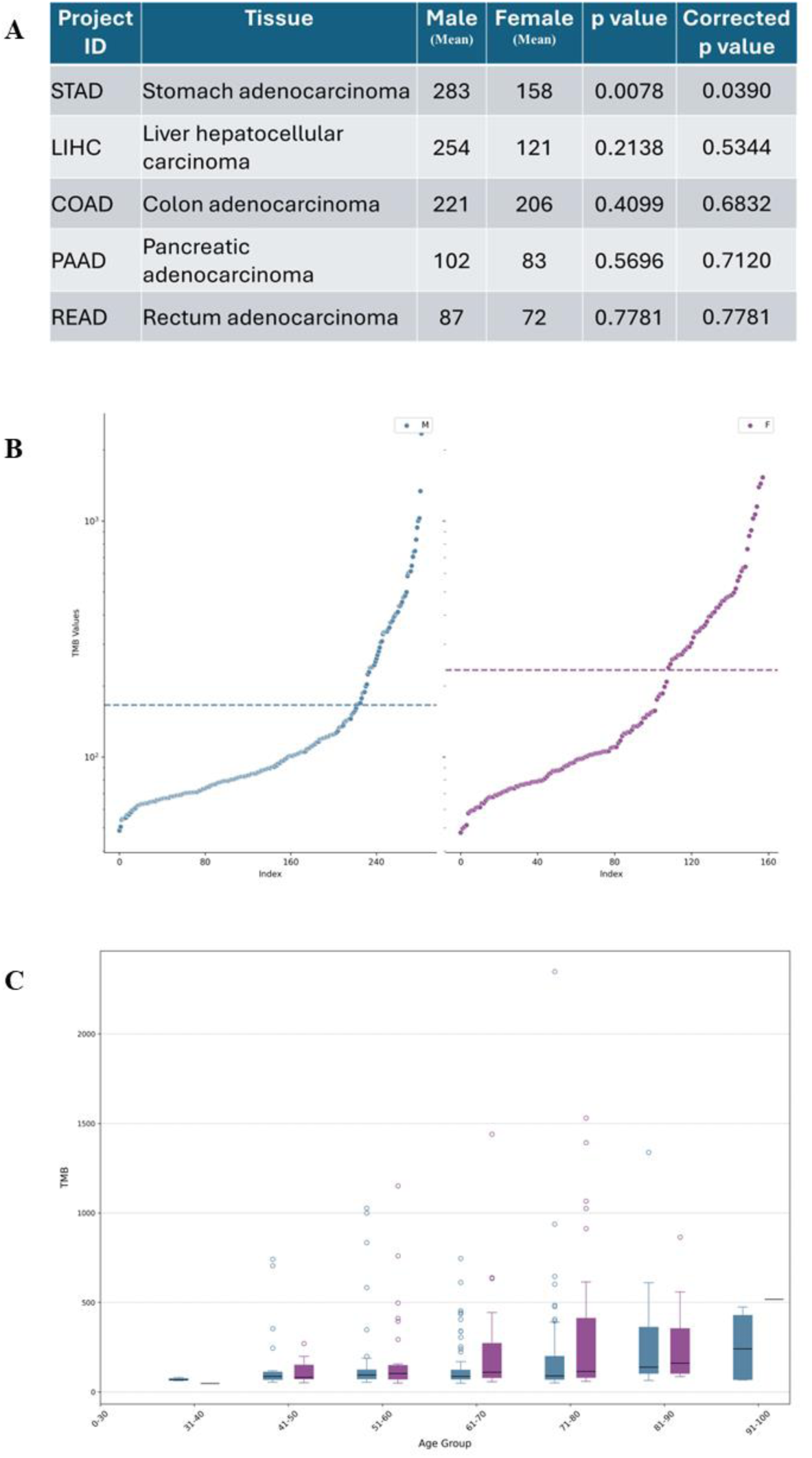
Autosomal tumor mutation burden in STAD-TCGA samples by sex, by age. (A) T-test analysis of male and female TMB values of Gastrointestinal TCGA projects; (B) Female and male TMB values scatter; (C) TMB comparison by age group and sex. (Males in black, Females in purple).

The mean TMB value of females was higher than that of males, particularly evident in the age groups of 61 to 70 and 71 to 80 (233.33 vs. 166.02, respectively. Fig. 1B and 1C).

### Sexual Dimorphism in Mutation Occurrence Rate is Evident in 1,052 Genes

Many genes contribute to the sex-related differences in TMB values; we specifically searched for genes that show a significant difference in mutation rate between the two sexes. A chi-square analysis was performed for each gene in the TCGA cancer panel (as elaborated in Methods), followed by Benjamini-Hochberg p-value correction. A total of 1,052 genes showed sexual dimorphism in mutation rate between males and females (Table S1). Of these, 460 exhibited higher mutation percentages in males, while 592 showed higher mutation percentages in females. These sexually dimorphic genes are distributed across all autosomes, yet some chromosomes contain fewer sex-biased genes than others (Fig. 2A, Fig. S1 and S2).

**Fig. 2:**
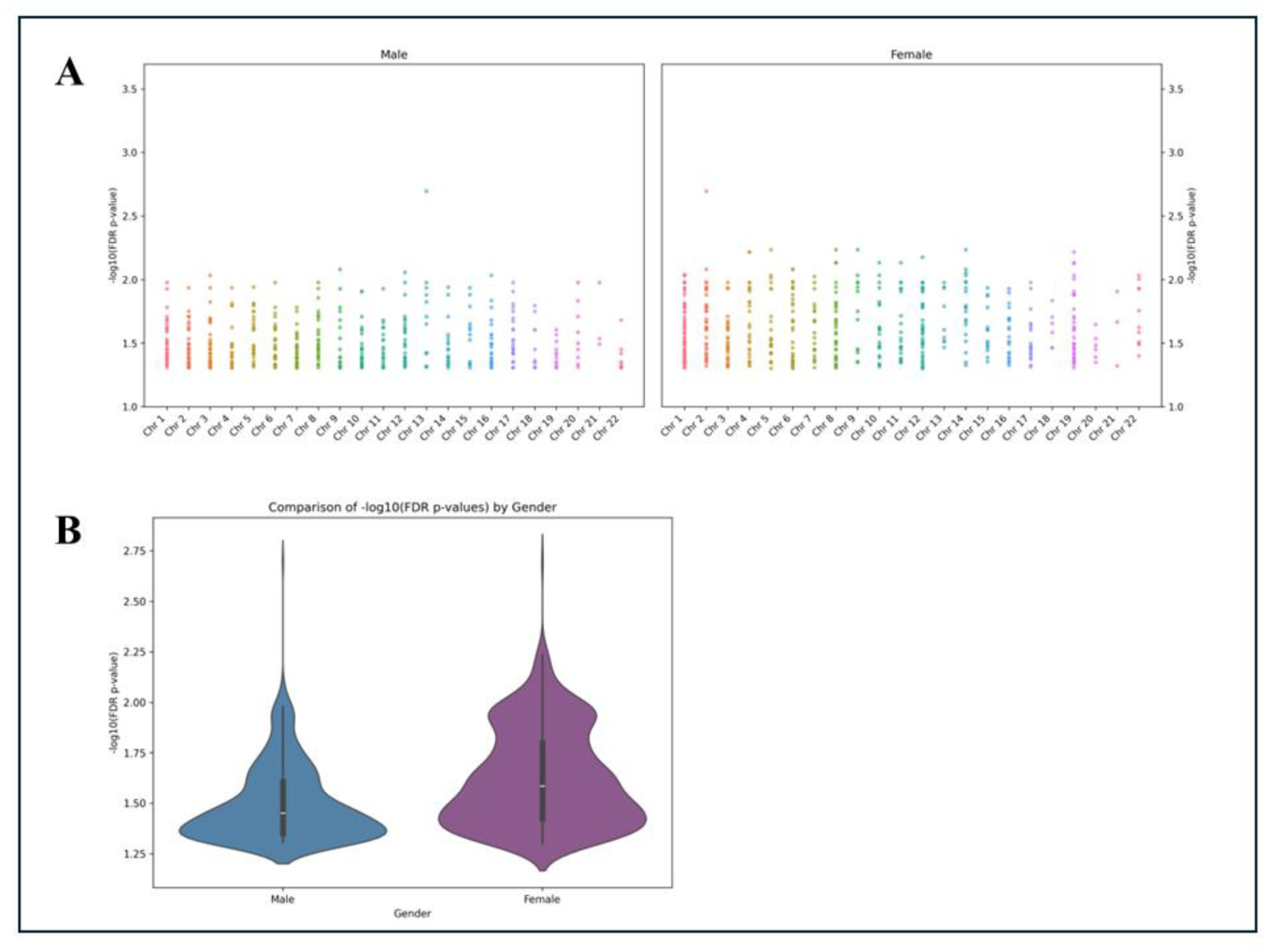
Sexual dimorphism in mutation occurrence rate is evident in 1052 genes. (A) Sexually dimorphic gene scatter by chromosomes according to higher mutation percentages in males or females. Genes assigned to the male plot (460) and, genes assigned to the female plot (592). (B) Mann-Whitney U Test comparison of differences in the distributions of statistical significance between the two groups.

We investigated whether the distributions of statistical significance differ between genes with higher mutation percentages in males and those with higher percentages in females. To compare the distributions of −log10(FDR p-values) between these two groups, we performed a Mann-Whitney U test for two independent samples, with normality assessed using the Shapiro-Wilk test (data not shown). The test yielded a p-value of −3.1916e-21, indicating a statistically significant difference between the distributions (Fig. 2B). This substantial difference suggests that the level of statistical significance varies between the groups, potentially reflecting underlying biological differences in mutation patterns between the sexes.

We further analyzed the two groups of sexually dimorphic genes with higher mutation percentages in males and females to assess whether they are functionally enriched in underlying biological mechanisms. The ClusterProfiler tool was utilized to conduct enrichment analyses of Gene Ontology (GO) terms and biological pathways significantly associated with our input gene lists [17]. The male gene group with a higher mutation rate is enriched with fourteen GO categories related to heterochromatin formation, epigenetic gene expression regulation, cell cycle phase transition, and hematopoietic progenitor cell differentiation. In the female gene group, enrichment is evident for the positive regulation of DNA repair and positive regulation of double-strand break repair GO categories (Fig. 3, Table S2).

**Fig. 3:**
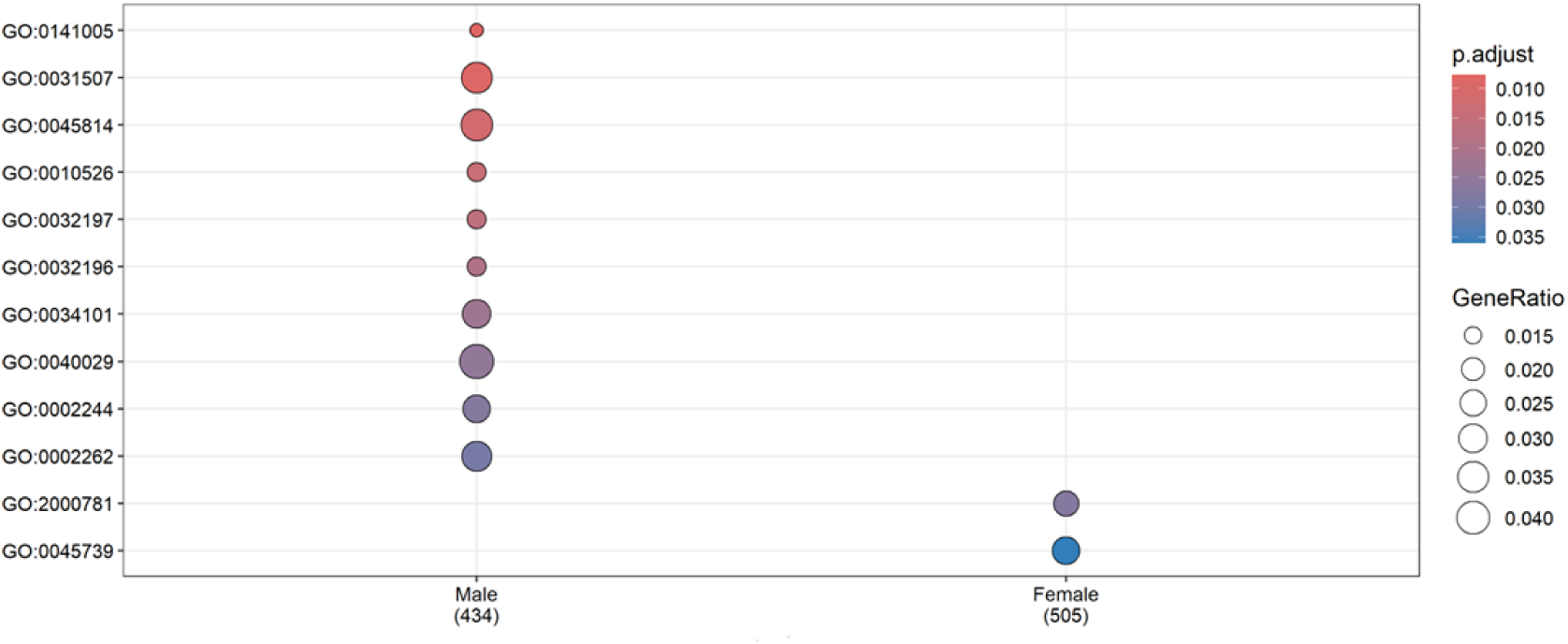
Enrichment analyses of Gene Ontology terms in male and female higher mutation rate gene groups. GO terms are as follows: GO:0141005 - retrotransposon silencing by heterochromatin formation; GO:0031507 - heterochromatin formation; GO:0045814 - negative regulation of gene expression, epigenetic; GO:0010526 - retrotransposon silencing; GO:0032197 – retrotransposition; GO:0032196 – transposition; GO:0034101 - erythrocyte homeostasis; GO:0040029 - epigenetic regulation of gene expression; GO:0002244 - hematopoietic progenitor cell differentiation; GO:0002262 - myeloid cell homeostasis; GO:0030218 - erythrocyte differentiation; GO:1901987 - regulation of cell cycle phase transition; GO:0006346 - DNA methylation-dependent heterochromatin formation; GO:0046885 - regulation of hormone biosynthetic process; GO:2000781 - positive regulation of double-strand break repair; GO:0045739 - positive regulation of DNA repair

### GC Sexually Dimorphic Mutated Genes are Represented in KEGG Cancer Pathways, COSMIC Gene Census Dataset, and the Tumor Suppressor Genes Database

Investigating sexually dimorphic GC mutated genes within authoritative cancer gene databases may enhance our understanding of their contributions to tumorigenesis and the differences between the sexes in cancer occurrence, progression, and mortality. We cross-referenced the 1,052 sexually dimorphic mutated genes with four databases: the KEGG Cancer Pathway, the KEGG Gastric Cancer Pathway [18], the COSMIC Gene Census [19], and the Tumor Suppressor Genes database [20]. Our analysis of overlapping genes among these five datasets highlights significant connections between sexually dimorphic GC mutated genes and established cancer-related gene sets (illustrated in Fig. 4; a complete list of overlapping genes is provided in Table S3). Specifically, thirty genes overlap between sexually dimorphic GC and the KEGG Cancer Pathway (**C1**), and ten genes overlap between sexually dimorphic GC and the KEGG Gastric Cancer Pathway (**C2**). Notably, eight genes overlap among all three sets (**C5**).

**Fig. 4:**
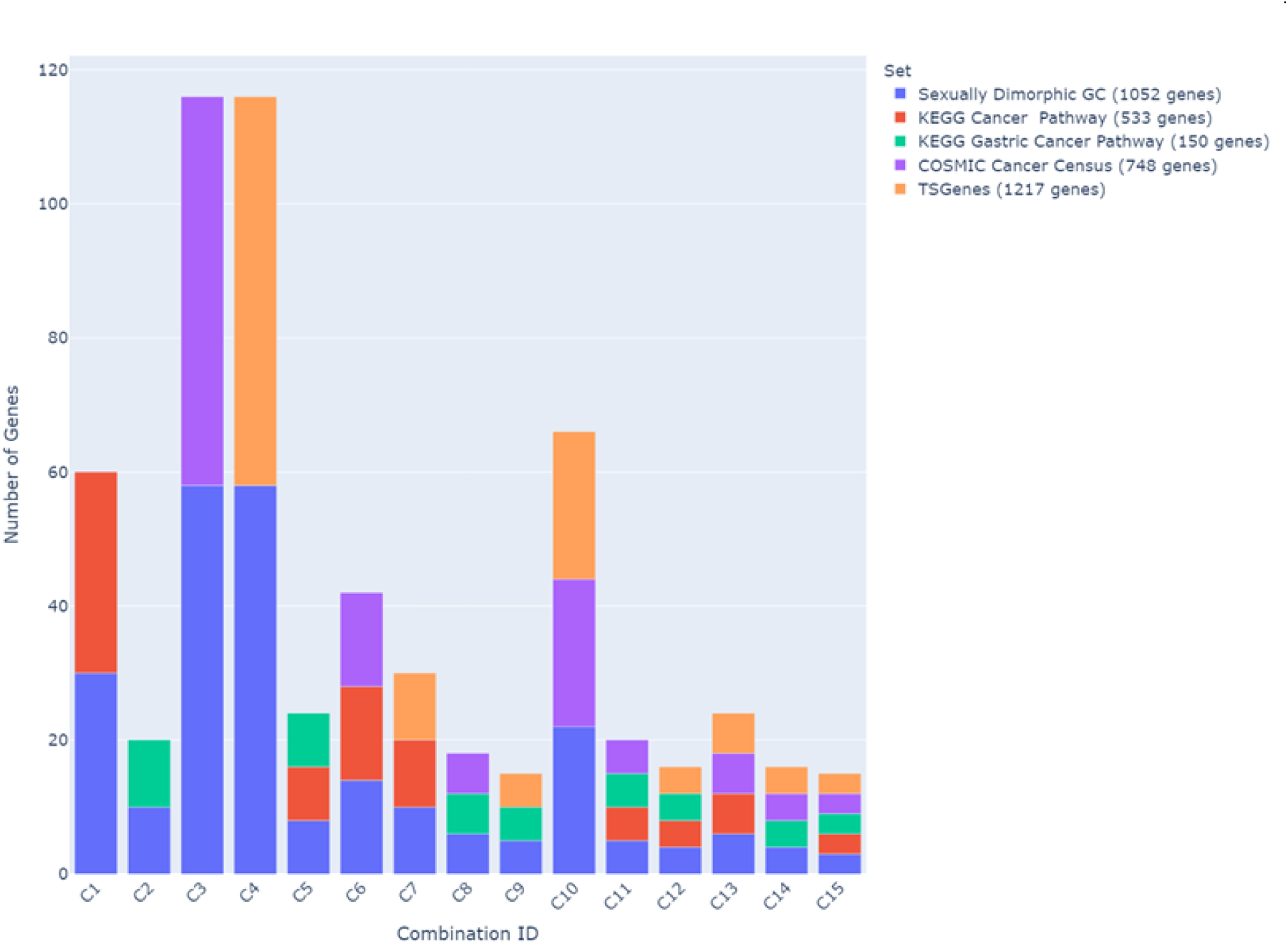
Upset plot, overlapping between sexually dimorphic GC mutated genes and the KEGG cancer pathway, the KEGG gastric cancer pathway, the COSMIC gene census, and the tumor suppressor genes databases. Number of overlapping genes in square brackets. C1: [30]; C2: [10]; C3: [58]; C4: [58]; C5: [8]; C6: [14]; C7: [10]; C8: [6]; C9: [5]; C10: [22]; C11: [5]; C12: [4]; C13: [6]; C14: [4]; C15: [3].

Fifty-eight genes overlap between sexually dimorphic GC and the COSMIC Cancer Census (C3), and a different set of fifty-eight genes overlap between sexually dimorphic GC and the Tumor Suppressor Genes set (C4) (notably, these two groups of genes are not identical). Twenty-two genes overlap among sexually dimorphic GC, the COSMIC Cancer Census, and the Tumor Suppressor Genes set (C10).

Fourteen genes overlap among sexually dimorphic GC, the KEGG Cancer Pathway, and the COSMIC Cancer Census (C6). Remarkably, all five datasets share three genes – Axin 1 (AXIN1), Glycogen Synthase Kinase 3 Beta (GSK3B), and RB Transcriptional Corepressor 1 (RB1) (C15), which are known for their critical roles in cancer development and progression. These genes may serve as key biomarkers or therapeutic targets, underscoring the importance of sexually dimorphic factors in gastric tumorigenesis.

### Gastric Cancer Co-Mutated Sexually Dimorphic Gene Patterns Differ Between the Sexes

The number of driver mutations required to initiate tumorigenesis has been extensively studied, with estimates ranging from three to approximately ten mutations, depending on the type of cancer, patient age, and the specific models or methodologies employed [21–24]. By investigating co-mutation patterns among sexually dimorphic mutated genes, we aimed to identify specific combinations of mutations that may synergistically drive tumorigenesis in a sex-specific context. We evaluated two-gene, three-gene, four-gene, and five-gene combinations among our sexually biased genes in males and females. The top ten co-mutation patterns are summarized (Fig. 5, detailed list in Table S4).

**Fig. 5:**
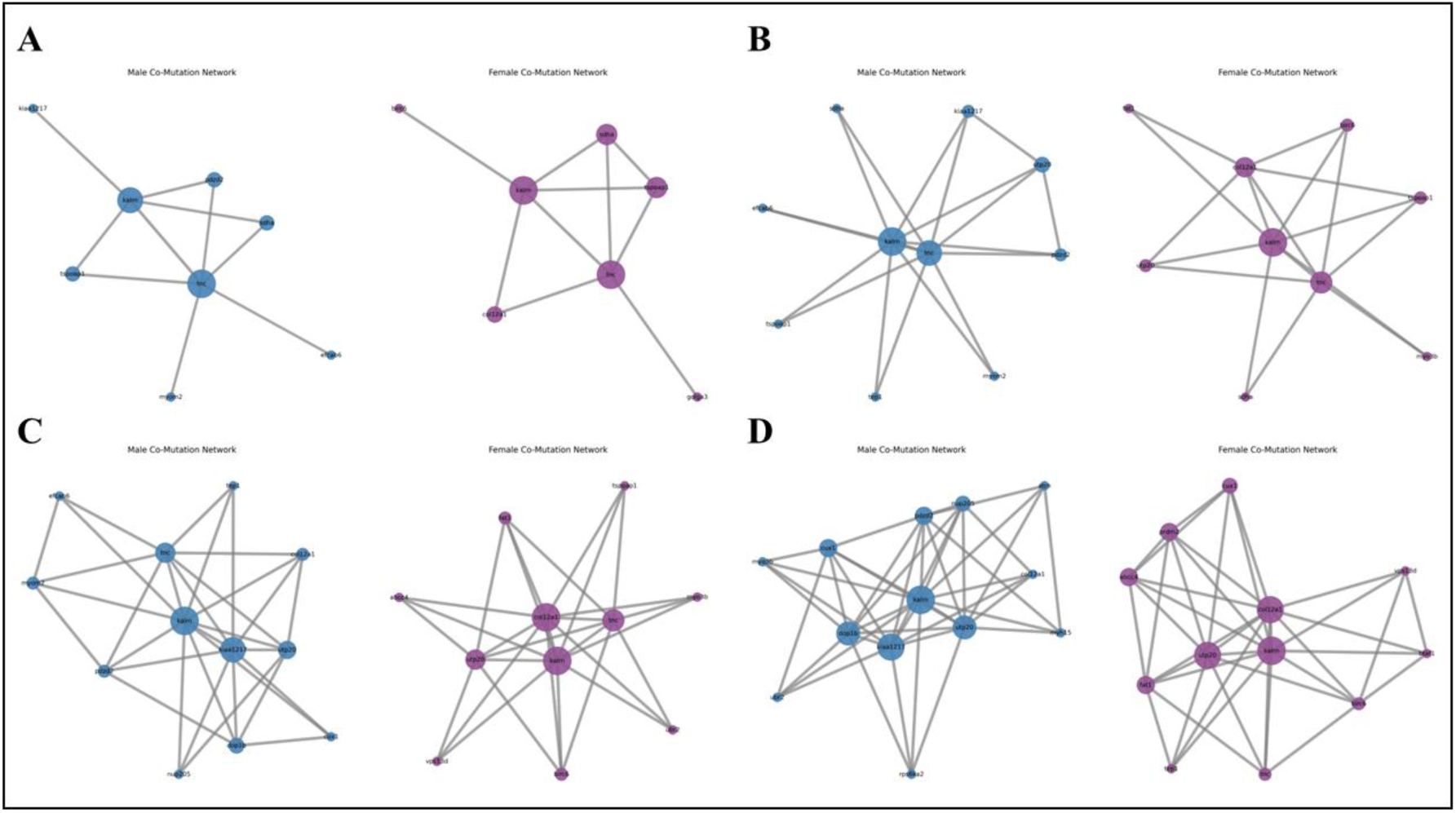
Sexual dimorphism in co-mutated gene patterns. A network plot visualization of the top ten co-mutated gene combinations for males (blue nodes) and females (purple nodes), including two-gene (A), three-gene (B), four-gene (C), and five-gene combinations (D).

Generally, males present higher co-mutation counts in the two-gene and three-gene levels. Yet, females tend to have higher counts in their top co-mutated gene combinations across four-gene to five-gene levels, suggesting more stable or frequent co-mutational events among these genes, which may have implications for disease progression and treatment responses.

Further, results indicate that male and female tumors differ in the most prevailed co-mutated gene patterns. Tenascin C (TNC), involved in extracellular matrix remodeling, and Kalirin RhoGEF Kinase (KALRN), playing a role in cytoskeleton organization and cell migration, are the most frequently co-mutated genes in both males and females, suggesting a potentially critical role in tumorigenesis of both sexes [25]. However, differences in the prevalence of genes in co-mutated combinations between males and females are evident. Several genes highly prevail in the top co-mutated gene combinations of one sex but to a lesser degree or none in the opposite sex (e.g., Collagen Type XII Alpha 1 Chain (COL12A1) highly prevails in females but to a lesser degree in males; PDZ Domain Containing 2 (PDZD2) appears in males’ top co-mutated combinations but is entirely absent from the female top list. The opposite behavior is detected for the multi-drug-resistant ATP Binding Cassette Subfamily C Member 4 (ABCC4) and the tumor suppressor gene PR/SET Domain 2 (PRDM2); Cut Like Homeobox 1 (CUX1) and UTP20 Small Subunit Processome Component (UTP20) appear in co-mutations in both genders but with different partners and frequencies). Hence, while certain genes are involved in tumorigenesis in both males and females, their interactions and pathways differ. Distinctive co-mutation combination patterns in males and females point toward sex-specific co-mutational networks that could reflect tumor biology variances and differential cancer development and progression.

### Tumor Sex Prediction Based on Mutation Patterns

We investigated the possibility that sex-related disparities in GC mutated genes might point to fundamentally distinct molecular networks - potentially suggesting two genetically different diseases sharing the same name. Specifically, we aimed to establish whether the differences in mutated gene patterns between males and females are substantial enough to underlie sex-specific pathways in tumorigenesis and thus allow accurate sex prediction. To predict the sex of GC tumor samples based on their gene mutation profiles, we employed a supervised machine-learning approach. Upon evaluating multiple models, the Random Forest classifier emerged as the best model based on mean accuracy (see Methods). The model was implemented twice, first utilizing the full panel of genes (∼32,000 autosomal genes) and second on the sexually dimorphic mutated gene sub-dataset (1,052 autosomal genes). In the case of the sexually dimorphic mutated genes sub-dataset, the model achieved an accuracy of 68.54%, a ROC-AUC of 72.62%, and an F1-score of 76.67%, effectively identifying positive cases while maintaining a balance between precision and recall. These results demonstrate the model’s ability to distinguish between male and female samples based on their mutation profiles (Fig. 6). Similar yet less optimal results were obtained once utilizing the full panel of genes. In general, slightly lower accuracy and ROC-AUC and similar F1-Scores were achieved (Fig. S3).

**Fig. 6:**
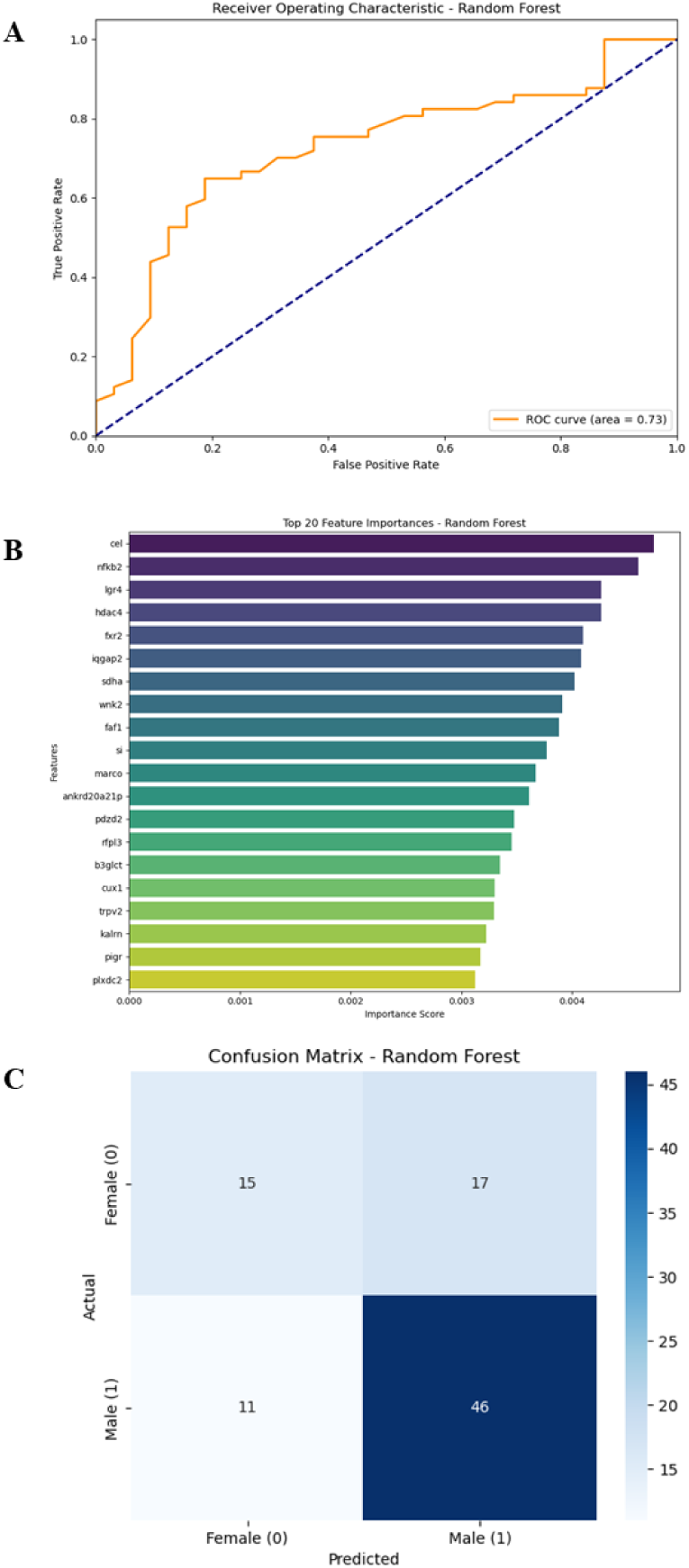
Performance of the Random Forest classifier in predicting tumor sample’s sex based on sexually dimorphic gene mutation profiles. **(A)** The Receiver Operating Characteristic (ROC) curve of the Random Forest model evaluated on the test set, illustrating its ability to distinguish between male and female gastric cancer samples, with an area under the curve (AUC) of 0.7262. **(B)** Feature importance plot displaying the top 20 genes ranked by their importance scores in the Random Forest model. **(C)** Confusion matrix summarizing the Random Forest model classification results on the test set.

## Discussion

Cancer prevalence, progression, and patients’ responsiveness to treatments differ between men and women, leading to significant gaps in cancer survival and mortality between the sexes [26–28]. This underscores the critical need for sex-specific personalized cancer characterization. In this study, we report the substantial differences in GC genomic alterations between men and women. By investigating single-nucleotide genomic disparities in GC, we aimed to gain insights into the molecular mechanisms underlying cancer development in each sex. This knowledge can inform the development of sex-tailored diagnostic markers and therapeutic strategies, potentially improving patient outcomes.

Analyzing single-nucleotide mutations documented for the 158 females and 283 males in the TCGA-STAD dataset, we characterized various aspects of genomic mutations, their associated genes, and their interactions. Our findings demonstrate that although both men and women are diagnosed with GC from a pathological standpoint, on a molecular level, they may exhibit distinct molecular pathways - suggesting that GC in males and females might be considered two separate yet similar diseases.

Autosomal TMB analysis of STAD-TCGA dataset indicates significant differences between men and women in the number of single-nucleotide, non-silent mutations identified in GC tumor genomes. This finding suggests that the utilization of TMB as a biomarker for ICI success must be reassessed in the context of patient sex.

A pertinent question arises regarding the source of TMB differences between males and females (Fig. 1C). Do female cells require a larger number of accumulating mutations before cancer occurs, or do they survive longer, allowing the number of accompanying mutations to grow? Our results indicate that the TMB value gap between the sexes is pronounced in post-menopausal years (Fig. 1C, age groups 60 to 80 years), supporting the notion of differences in DNA repair mechanisms contributing to gastric tumor development in males and females.

DNA repair mechanisms are essential for maintaining genomic stability by preventing DNA damage caused by environmental factors, metabolic stress, and replication errors. Sex-based differences in DNA repair efficiency have been documented, with females generally exhibiting more robust repair capabilities than males, partly due to the influence of sex steroid hormones (SSHs) [29]. The interplay between SSHs and DNA repair is complex; androgens regulate components essential for DNA double-strand break repair, while estrogens enhance repair capacity by upregulating DNA repair genes, promoting cell proliferation and enabling tolerance to chemotherapy and radiotherapy, yet contributing to hormone-dependent cancer progression. The increase in female TMB values during post-menopausal age may hint at the protective role of estrogen against gastric cancer in younger women. Although most women begin the menopausal transition between the ages of 45 and 55, many are prescribed hormone replacement therapy (HRT), potentially delaying the TMB-related postmenopausal effect in older ages. Unfortunately, TCGA clinical records lack information on HRT consumption, preventing the accurate incorporation of this parameter into our analysis.

We identified 1,052 sexually dimorphic mutated genes showing significant differences in mutation occurrence rate, with 460 genes exhibiting higher mutation rates in males and 592 genes in females. These two groups significantly differ, potentially reflecting underlying biological differences in mutation patterns between the sexes. GO enrichment analysis revealed sex-specific enrichments: the female group showed enrichment in categories related to positive regulation of DNA repair, aligning with documented differences in DNA repair efficacy and higher TMB in females.

Sex-based genomic differences in GC are not limited to single-nucleotide mutation occurrence rate. A recent study exploring structural variant signatures across cancer genomes – specifically focusing on transcription-replication collision footprints – reported an abundance of large tandem duplications in female-enriched upper gastrointestinal tract cancers, including GC [30]. Suggesting that genome integrity, as a whole, may be more compromised in female GC patients.

Many prevalent cancer projects utilize specific and limited gene panels (e.g., DFCI-Oncopanel - 275 to 447 genes; MSK-IMPACT 342 to 468 genes, and FoundationOne 324to 511 genes), which may not capture the sexually dimorphic biomarkers identified in our study, constituting approximately merely 3% of the TCGA whole-gene panel. If sexual dimorphism is to be considered a factor in molecular cancer characterization, gene panels should be broadened and updated accordingly.

We found that sexually dimorphic GC mutated genes overlap with authoritative cancer-related pathways - the KEGG Cancer Pathway, the KEGG Gastric Cancer Pathway, the COSMIC Gene Census, and the Tumor Suppressor Genes database - underscoring their potential role in tumorigenesis and emphasizing their importance in cancer biology. Notably, three genes - the G-protein signaling regulator (AXIN1), the negative regulator of the cell cycle (RB1), and the negative regulator in the hormonal control of glucose homeostasis (GSK3B) – were common to all datasets. AXIN1 and RB1, more mutated in males, and GSK3B, more mutated in females, are well-documented to their central roles in cancer formation. Interestingly AXIN1 and GSK3B interact within the Wnt signaling pathway, known to be associated with various cancers and responsive to SSHs [31].

Analyzing co-mutations among sexually dimorphic genes, we observed that male and female tumors differ in the most prevalent co-mutated genes. Females showed a tendency for higher counts of co-mutated gene combinations at the three-gene and four-gene levels, suggesting more stable or frequent co-mutational events. This may have implications for disease progression and highlights the potential value of identifying female-specific genomic profiles for tailored treatment development.

Tailored treatments in cancer, such as immunotherapy, targeted radiotherapy, and microbiome-based therapies, are increasingly focusing on molecular classification rather than tumor origin [32–34]. Clinical trials have demonstrated improved outcomes based on molecular characteristics of metastatic tumors [35]. Our approach suggests that females not only present a more conserved genomic profile in GC but also exhibit sex-unique mutation combinations. This supports the notion that GC in males and females may be considered distinct diseases deserving separate characterization and treatment strategies.

Utilizing a supervised machine learning approach, we employed sexually dimorphic mutated gene patterns to predict tumor sex. The Random Forest classifier effectively identified sample sex with an accuracy of 68.54%, demonstrating that these mutation patterns are central to GC sex-biased characteristics and can be used to differentiate male and female samples. Given that cancerous cell mutations accumulate during tumor progression, we do not expect mutation patterns to be entirely uniform. However, the presence of male-like and female-like characteristic patterns suggests that integrating sexual dimorphism genomic alteration analysis in cancer diagnosis could enhance personalized treatment, regardless of patient sex.

These differences may underlie observed disparities in cancer susceptibility, progression, and therapeutic response between the sexes. Addressing male and female GC separately in tumor characterization and classification is a critical step toward more effective treatments for GC patients. Moreover, this study emphasizes the need for further research to elucidate sex-specific contributions to tumorigenesis and how sex-biased genomic alterations can serve as potential biomarkers or therapeutic targets not only in GC but also in other cancers.

## Methods

### Data Downloading and Processing

Gastrointestinal cancer datasets were downloaded from The Cancer Genome Atlas (TCGA) via the Genomic Data Commons (GDC) data portal [36]. Mutation Annotation Format (MAF) files for the following projects were obtained and analyzed: stomach adenocarcinoma (STAD), colon adenocarcinoma (COAD), rectum adenocarcinoma (READ), hepatocellular carcinoma (LIHC), and pancreatic adenocarcinoma (PAAD). Case records were excluded if the sex field was missing or indicated as “other.” The final dataset included: **STAD:** 283 males and 158 females; **COAD:** 221 males and 206 females; **READ:** 87 males and 72 females; **LIHC:** 254 males and 121 females; **PAAD:** 102 males and 83 females. Esophageal carcinoma (ESCA) was excluded due to the small number of female samples (157 males and 27 females).

### Tumor Mutational Burden Analysis

Tumor mutational burden (TMB) values were calculated separately for males and females. Only non-silent mutations in autosomal genes were included. The total number of mutations per sample was divided by 35.8 megabases (Mb), representing the size of the autosomal exome, to obtain the TMB value for each sample.

### Statistical Analysis

All statistical analyses were performed using Python (version 3.12) and R (version 4.4.2). Python was utilized for data processing and model development, while R was used for data visualization.

### TMB Comparison

A two-tailed t-test for unequal variances was applied to compare TMB values between males and females, followed by Benjamini-Hochberg false discovery rate (FDR) correction to adjust for multiple comparisons.

### Identification of Sexually Dimorphic Mutated Genes

The chi-square test was used to identify genes with significant differences in mutation rates between sexes, followed by FDR correction.

### Distribution Analysis

The Mann-Whitney U test compared the distributions of statistical significance between genes with higher mutation percentages in males versus females. Normality was assessed using the Shapiro-Wilk test.

### Significance Threshold

Statistical significance was determined at a P-value < 0.05.

### Mutated Gene Sexual Dimorphism Analysis

Mutation occurrence was calculated for all autosomal genes (n = 32,817) in the TCGA panel. A gene was considered mutated if it contained one or more mutations in a sample. Genes showing significant differences in mutation rates between sexes after FDR correction were classified as sexually dimorphic.

Gene location data were obtained from the UCSC Genome Browser (human genome assembly January 2022). Scatter plots were created to visualize the distribution of sexually dimorphic genes across chromosomes. The genetic map illustrating the locations of these genes was generated using the MG2C online tool [37].

### Cancer-Related Gene Lists

Gene lists were obtained from authoritative cancer databases to identify overlaps with sexually dimorphic mutated genes: KEGG Cancer Pathway and KEGG Gastric Cancer Pathway [18], COSMIC Gene Census [19], and Tumor Suppressor Genes Database [20].

### Gene Ontology Analysis

Gene Ontology (GO) enrichment analysis was performed using the ClusterProfiler tool [17] to identify biological processes and pathways significantly associated with the sexually dimorphic mutated genes.

### Co-Mutation Combination Analysis

The matrix of 1,052 sexually dimorphic mutated genes, containing mutation information for each gene in each tumor sample, was used for co-mutation analysis. Two-gene, three-gene, four-gene, and five-gene co-mutation combinations were identified. Due to the exponential increase in possible unique combinations (∼552,000 two-gene combinations; ∼193 million three-gene combinations; ∼53 billion four-gene combinations; ∼11 trillion five-gene combinations), the analysis of four-gene and five-gene combinations focused on the top 10% most frequently mutated genes to reduce computational complexity.

### Tumor Sex Prediction Analysis

A supervised machine learning approach was employed to predict the sex of gastric cancer samples based on gene mutation data. All models were implemented using scikit-learn (version 1.0.2). The dataset consisted of mutation profiles for either 1,052 sexually dimorphic autosomal genes or the full set of 32,817 autosomal genes, labeled as ‘female’ or ‘male’.

### Data Splitting

The dataset was divided into training and test sets using an 80:20 ratio, stratified by sex to maintain class balance.

### Feature Selection

The SelectKBest method with mutual information (mutual_info_classif) as the scoring function was applied to select the top 1,000 genes.

### Model Evaluation

Four machine learning models were initially evaluated: Support Vector Machine (SVM) with a linear kernel, Random Forest Classifier, XGBoost and Logistic Regression [38].

### Cross-Validation

Stratified 5-fold cross-validation was performed on the training set to consistently evaluate model performance using metrics such as mean accuracy, area under the receiver operating characteristic curve (ROC-AUC), and F1-score.

### Model Selection and Tuning

The Random Forest Classifier demonstrated superior performance and was selected as the optimal model. Hyperparameter tuning was conducted using GridSearchCV with 5-fold cross-validation on the training set to optimize parameters like the number of estimators, maximum depth, and minimum samples required for splitting.

### Model Evaluation on Test Set

The optimized Random Forest model was evaluated on the independent test set. Performance metrics included accuracy, ROC-AUC, and F1-score. Visualizations of the ROC curve and confusion matrix were generated to illustrate model performance.

### Feature Importance Analysis

The Random Forest model’s feature importances were extracted to identify the most influential genes in predicting sex, highlighting potential biomarkers for sex-specific differences in gastric cancer.

## List of abbreviations

TMB: Tumor mutational burden
GC: Gastric Cancer
TCGA-STAD: The Cancer Genome Atlas Stomach Adenocarcinoma
ICIs: Immune checkpoint inhibitors
GO: Gene Ontology
SSHs: Sex Steroid Hormones
HRT: Hormone Replacement Therapy

## Declarations

### Ethics approval and consent to participate

Not applicable

### Consent for publication

Not applicable.

### Availability of data and materials

The TCGA datasets that support this study’s findings are available at The Cancer Genome Atlas (TCGA) via the Genomic Data Commons (GDC) data portal (https://portal.gdc.cancer.gov/projects/TCGA-STAD). All other relevant data are in the manuscript and Supporting Information files.

### Authors’ contributions

D.S. - Study conception and design; N.K. - data collection and algorithms programming; N.K. and D.S. - data analysis and interpretation of results; D.S. - manuscript preparation; All authors reviewed the manuscript.

## Acknowledgments

We thank A. Ben-Yehuda for critically reading the manuscript, and D. Dahary for fruitful discussions.

## Supplementary Materials

Figs. S1 to S3

Tables S1 to S4

