## Supplemental Data 1 for "Sex Disparities in Gastric Cancer Tumor Mutation Burden and Gene Mutation Patterns"

Supplementary Materials:

- Figs. S1 to S3
- Tables S1 to S4

**Fig. S1: Autosomal scatter of the sexually dimorphic genes.** Male and female prevail sexually dimorphic genes by sex (black for males, purple for females) by chromosome genomic location.

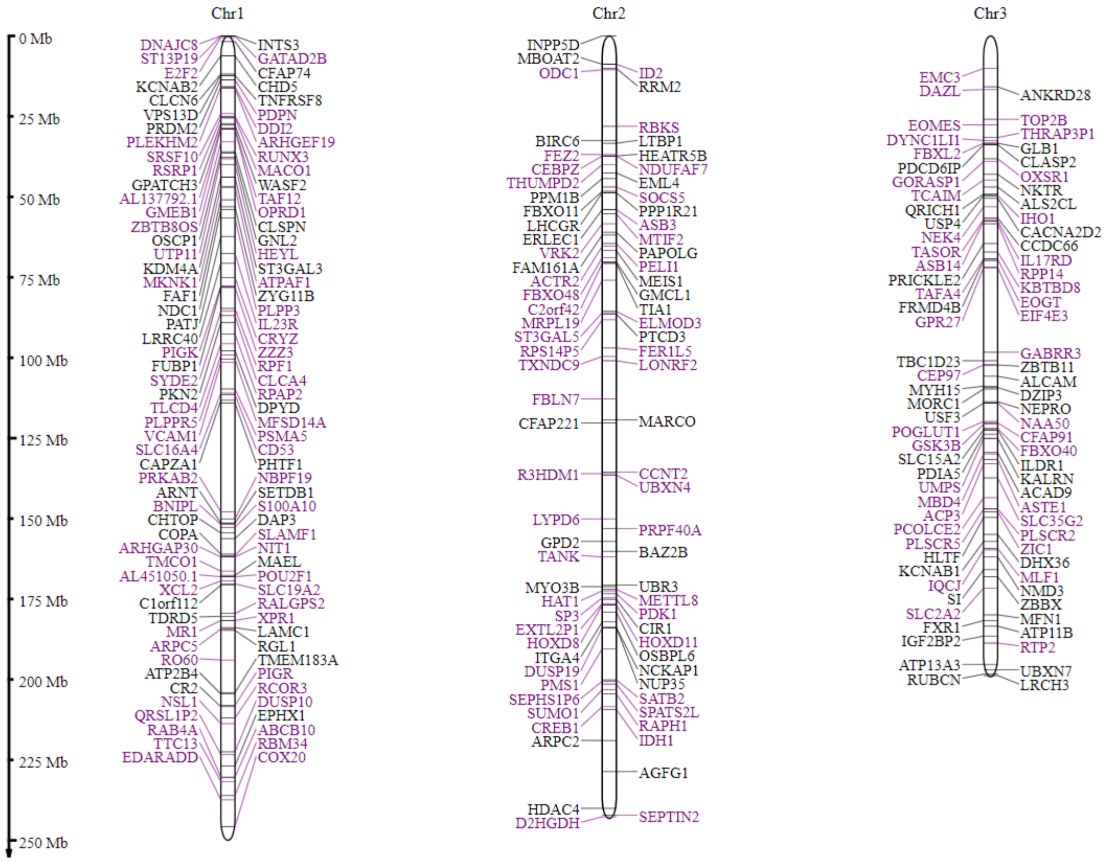

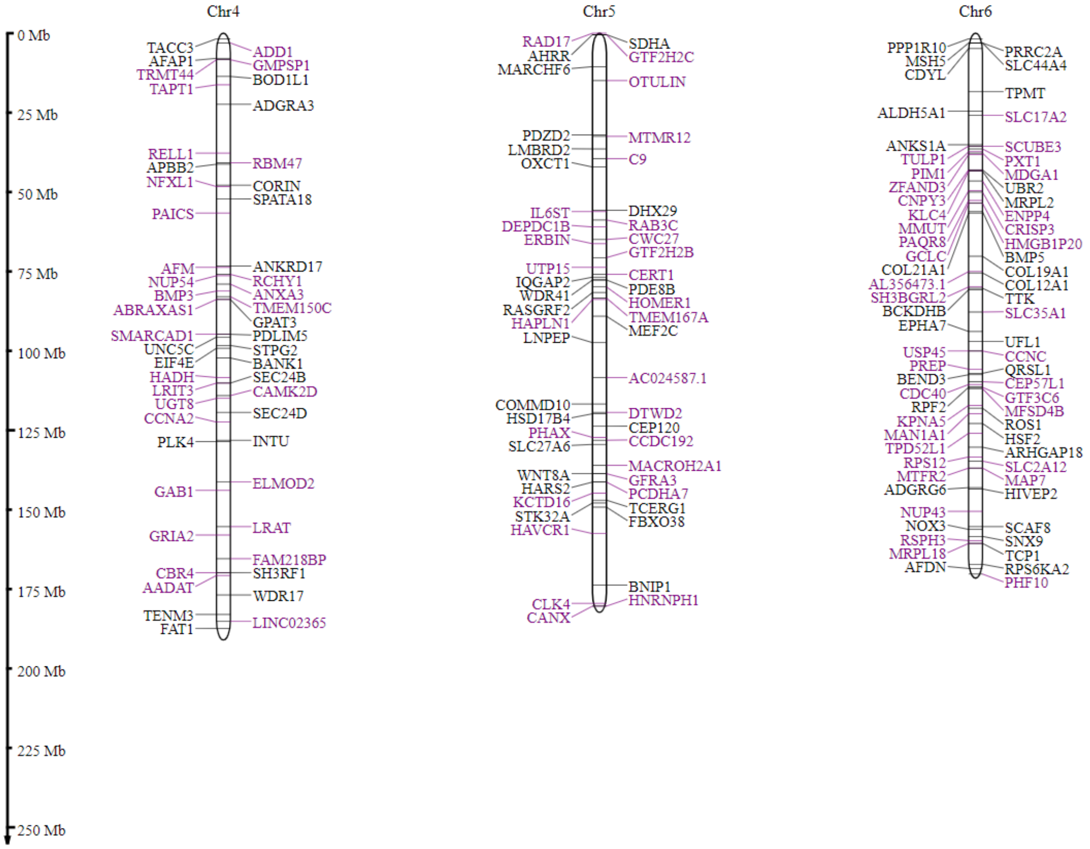
0 Mb
25 Mb
50 Mb
75 Mb
100 Mb
125 Mb
150 Mb
175 Mb
200 Mb
225 Mb
250 Mb
Chr4
TACC3
AFAP1
TRMT44
TAPT1
RELL1
APBB2
NFXL1
PAICS
AFM
NUP54
BMP3
ABRAXAS1
SMARCAD1
UNC5C
EIF4E
HADH
LRIT3
UGT8
CCNA2
PLK4
GAB1
GRIA2
CBR4
AADAT
TENM3
FAT1
ADD1
GMPSP1
BOD1L1
ADGRA3
RBM47
CORIN
SPATA18
ANKRD17
RCHY1
ANXA3
TMEM150C
GPAT3
PDLIM5
STPG2
BANK1
SEC24B
CAMK2D
SEC24D
INTU
ELMOD2
LRAT
FAM218BP
SH3RF1
WDR17
LINC02365
Chr5
RAD17
AHRR
MARCHF6
PDZD2
LMBRD2
OXCT1
IL6ST
DEPDC1B
ERBIN
UTP15
IQGAP2
WDR41
RASGRF2
HAPLN1
LNPEP
COMMD10
HSD17B4
PHAX
SLC27A6
WNT8A
HARS2
KCTD16
STK32A
HAVCR1
CLK4
CANX
SDHA
GTF2H2C
OTULIN
MTMR12
C9
DHX29
RAB3C
CWC27
GTF2H2B
CERT1
PDE8B
HOMER1
TMEM167A
MEF2C
AC024587.1
DTWD2
CEP120
CCDC192
MACROH2A1
GFRA3
PCDHA7
TCERG1
FBXO38
BNIP1
HNRNPH1
Chr6
PPP1R10
MSH5
CDYL
ALDH5A1
ANKS1A
TULP1
PIM1
ZFAND3
CNPY3
KLC4
MMUT
PAQR8
GCLC
COL21A1
AL356473.1
SH3BGRL2
BCKDHB
EPHA7
USP45
PREP
BEND3
CDC40
RPF2
KPNA5
MAN1A1
TPD52L1
RPS12
MTFR2
ADGRG6
NUP43
NOX3
RSPH3
MRPL18
AFDN
PRRC2A
SLC44A4
TPMT
SLC17A2
SCUBE3
PXT1
MDGA1
UBR2
MRPL2
ENPP4
CRISP3
HMGB1P20
BMP5
COL19A1
COL12A1
TTK
SLC35A1
UFL1
CCNC
QRSL1
CEP57L1
GTF3C6
MFSD4B
ROS1
HSF2
ARHGAP18
SLC2A12
MAP7
HIVEP2
SCAF8
SNX9
TCP1
RPS6KA2
PHF10

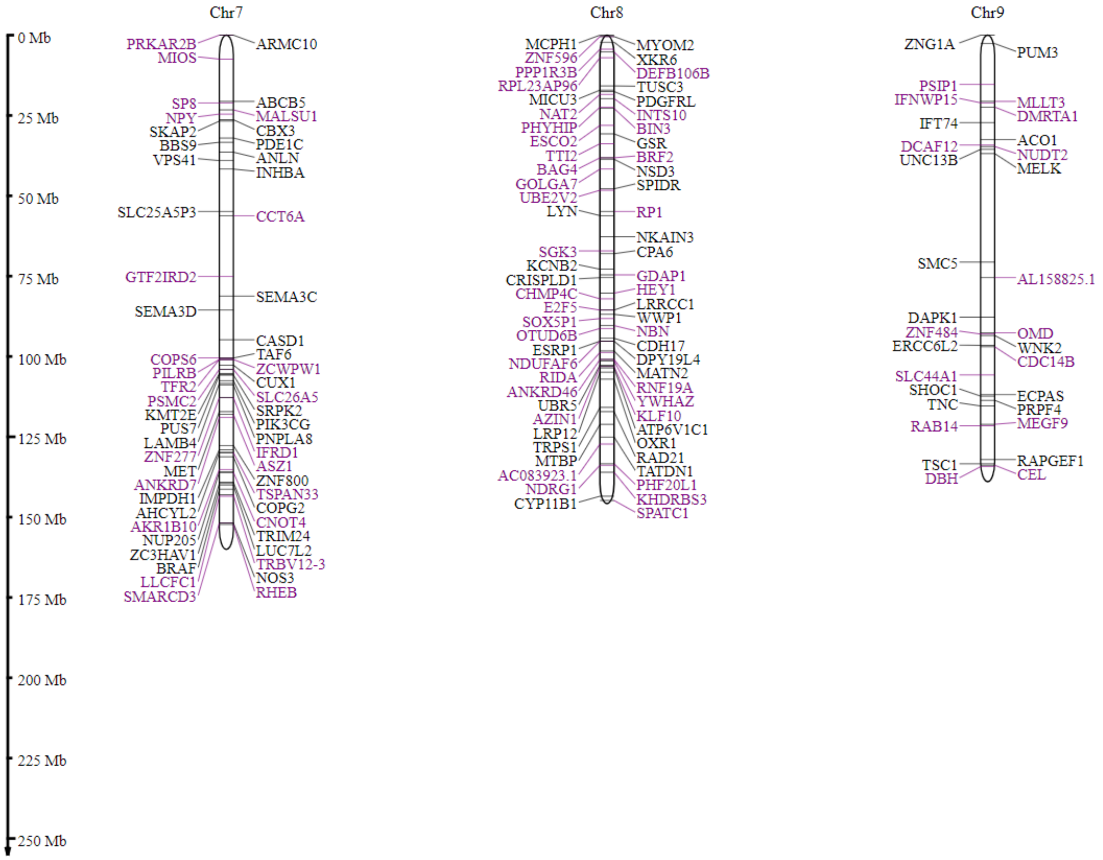
Chr7
Chr8
Chr9
0 Mb
25 Mb
50 Mb
75 Mb
100 Mb
125 Mb
150 Mb
175 Mb
200 Mb
225 Mb
250 Mb
PRKAR2B
MIOS
ARMC10
SP8
NPY
SKAP2
BBS9
VPS41
ABCB5
MALSU1
CBX3
PDE1C
ANLN
INHBA
SLC25A5P3
CCT6A
GTF2IRD2
SEMA3C
SEMA3D
CASD1
TAF6
COPS6
PILRB
TFR2
PSMC2
KMT2E
PUS7
LAMB4
ZNF277
MET
ANKRD7
IMPDH1
AHCYL2
AKR1B10
NUP205
ZC3HAV1
BRAF
LLCFC1
SMARCD3
ZCWPW1
CUX1
SLC26A5
SRPK2
PIK3CG
PNPLA8
IFRD1
ASZ1
ZNF800
TSPAN33
COPG2
CNOT4
TRIM24
LUC7L2
TRBV12-3
NOS3
RHEB
MCPH1
ZNF596
PPP1R3B
RPL23AP96
MICU3
NAT2
PHYHIP
ESCO2
TTI2
BAG4
GOLGA7
UBE2V2
LYN
MYOM2
XKR6
DEFB106B
TUSC3
PDGFRL
INTS10
BIN3
GSR
BRF2
NSD3
SPIDR
RP1
NKAIN3
CPA6
SGK3
KCNB2
CRISPLD1
CHMP4C
E2F5
SOX5P1
OTUD6B
ESRP1
NDUFAF6
RIDA
ANKRD46
UBR5
AZIN1
LRP12
TRPS1
MTBP
AC083923.1
NDRG1
CYP11B1
GDAP1
HEY1
LRRCC1
WWP1
NBN
CDH17
DPY19L4
MATN2
RNF19A
YWHAZ
KLF10
ATP6V1C1
OXR1
RAD21
TATDN1
PHF20L1
KHDRBS3
SPATC1
ZNG1A
PUM3
PSIP1
IFNWP15
IFT74
DCAF12
UNC13B
MLLT3
DMRTA1
ACO1
NUDT2
MELK
SMC5
AL158825.1
DAPK1
ZNF484
ERCC6L2
OMD
WNK2
CDC14B
SLC44A1
SHOC1
TNC
ECPAS
PRPF4
RAB14
MEGF9
TSC1
DBH
RAPGEF1
CEL

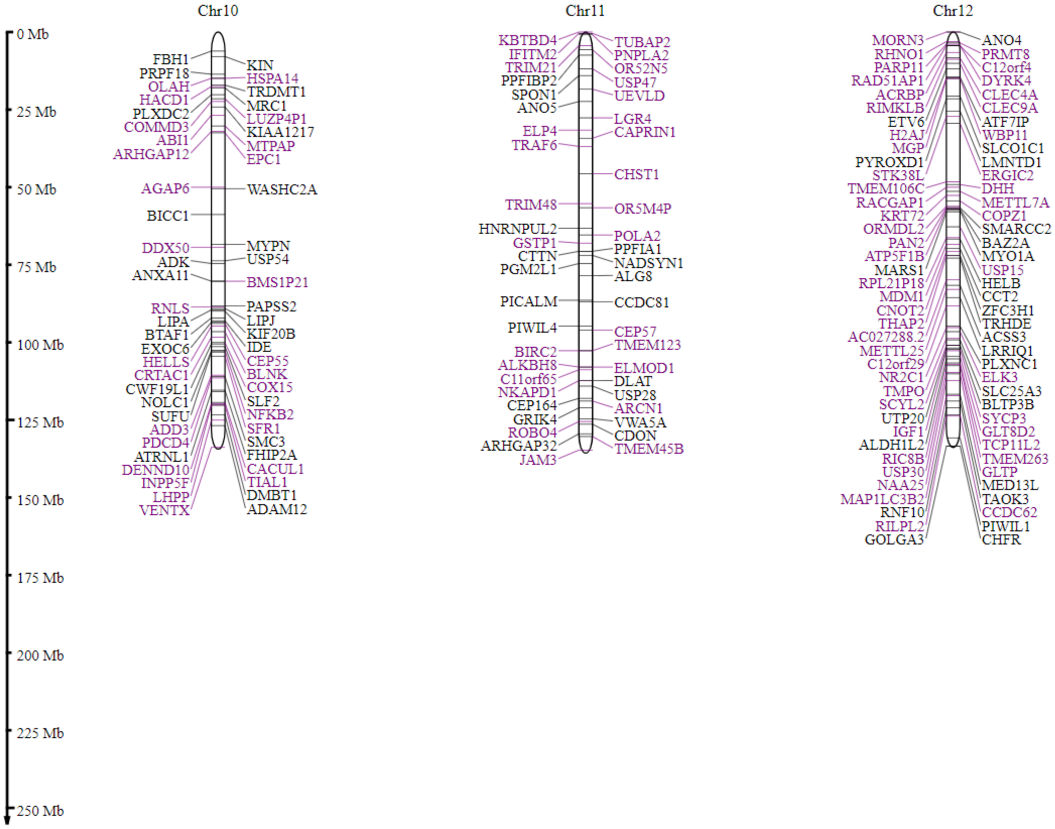
Chr10
Chr11
Chr12
0 Mb
25 Mb
50 Mb
75 Mb
100 Mb
125 Mb
150 Mb
175 Mb
200 Mb
225 Mb
250 Mb
FBH1
PRPF18
OLAH
HACD1
PLXDC2
COMMD3
ABI1
ARHGAP12
KIN
HSPA14
TRDMT1
MRC1
LUZP4P1
KIAA1217
MTPAP
EPC1
AGAP6
WASHC2A
BICC1
DDX50
ADK
ANXA11
MYPN
USP54
BMS1P21
RNLS
LIPA
BTAF1
EXOC6
HELLS
CRTAC1
CWF19L1
NOLC1
SUFU
ADD3
PDCD4
ATRNL1
DENND10
INPP5F
LHPP
VENTX
PAPSS2
LIPJ
KIF20B
IDE
CEP55
BLNK
COX15
SLF2
NFKB2
SFR1
SMC3
FHIP2A
CACUL1
TIAL1
DMBT1
ADAM12
KBTBD4
IFITM2
TRIM21
PPFIBP2
SPON1
ANO5
TUBAP2
PNPLA2
OR52N5
USP47
UEVLD
LGR4
CAPRIN1
ELP4
TRAF6
CHST1
TRIM48
OR5M4P
HNRNPUL2
GSTP1
CTTN
PGM2L1
POLA2
PPFIA1
NADSYN1
ALG8
PICALM
CCDC81
PIWIL4
CEP57
TMEM123
BIRC2
ALKBH8
C11orf65
NKAPD1
CEP164
GRIK4
ROBO4
ARHGAP32
JAM3
ELMOD1
DLAT
USP28
ARCN1
VWA5A
CDON
TMEM45B
MORN3
RHNO1
PARP11
RAD51AP1
ACRBP
RIMKLB
ETV6
H2AJ
MGP
PYROXD1
STK38L
TMEM106C
RACGAP1
KRT72
ORMDL2
PAN2
ATP5F1B
MARS1
RPL21P18
MDM1
CNOT2
THAP2
AC027288.2
METTL25
C12orf29
NR2C1
TMPO
SCYL2
UTP20
IGF1
ALDH1L2
RIC8B
USP30
NAA25
MAP1LC3B2
RNF10
RILPL2
GOLGA3
ANO4
PRMT8
C12orf4
DYRK4
CLEC4A
CLEC9A
ATF7IP
WBP11
SLCO1C1
LMNTD1
ERGIC2
DHH
METTL7A
COPZ1
SMARCC2
BAZ2A
MYO1A
USP15
HELB
CCT2
ZFC3H1
TRHDE
ACSS3
LRRIQ1
PLXNC1
ELK3
SLC25A3
BLTP3B
SYCP3
GLT8D2
TCP11L2
TMEM263
GLTP
MED13L
TAOK3
CCDC62
PIWIL1
CHFR

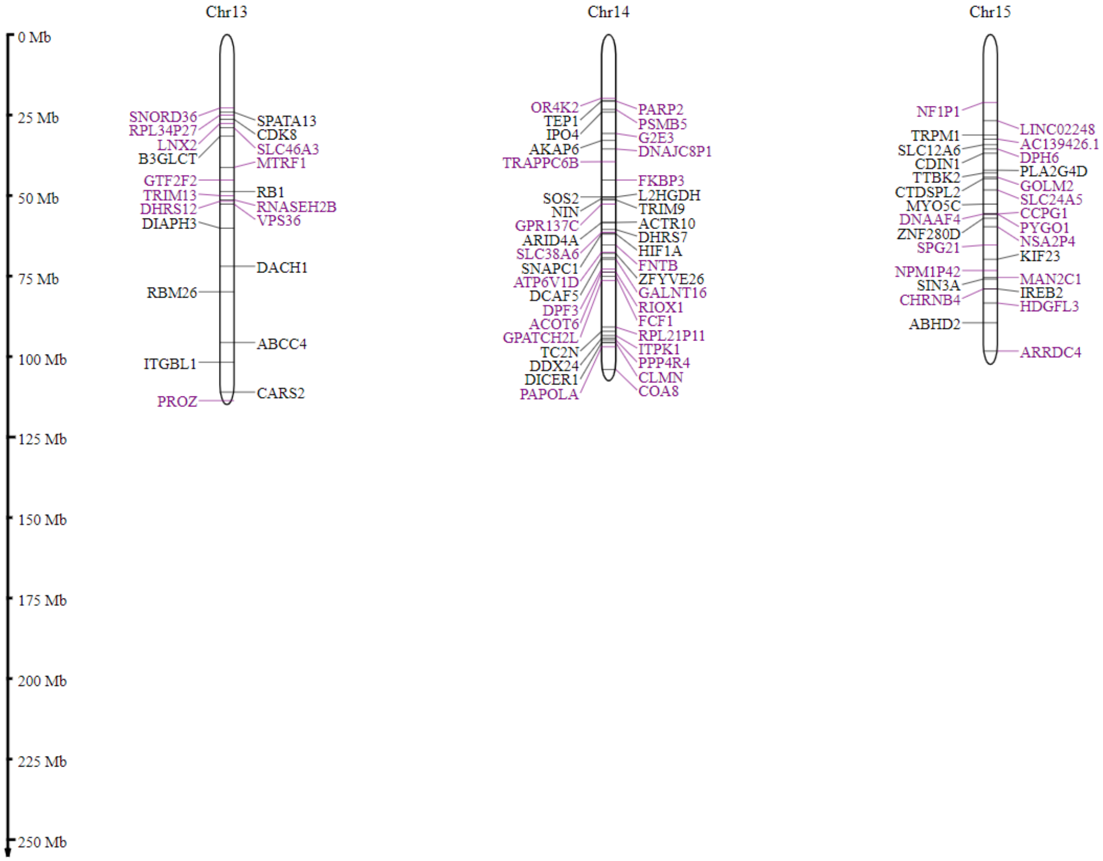

Chr13
Chr14
Chr15
0 Mb
25 Mb
50 Mb
75 Mb
100 Mb
125 Mb
150 Mb
175 Mb
200 Mb
225 Mb
250 Mb
SNORD36
RPL34P27
LNX2
B3GLCT
SPATA13
CDK8
SLC46A3
MTRF1
GTF2F2
TRIM13
DHRS12
DIAPH3
RB1
RNASEH2B
VPS36
DACH1
RBM26
ABCC4
ITGBL1
CARS2
PROZ
OR4K2
TEP1
IPO4
AKAP6
TRAPPC6B
PARP2
PSMB5
G2E3
DNAJC8P1
FKBP3
SOS2
NIN
GPR137C
ARID4A
SLC38A6
SNAPC1
ATP6V1D
DCAF5
DPF3
ACOT6
GPATCH2L
TC2N
DDX24
DICER1
PAPOLA
L2HGDH
TRIM9
ACTR10
DHRS7
HIF1A
FNTB
ZFYVE26
GALNT16
RIOX1
FCF1
RPL21P11
ITPK1
PPP4R4
CLMN
COA8
NF1P1
TRPM1
SLC12A6
CDIN1
TTBK2
CTDSPL2
MYO5C
DNAAF4
ZNF280D
SPG21
NPM1P42
SIN3A
CHRNB4
ABHD2
LINC02248
AC139426.1
DPH6
PLA2G4D
GOLM2
SLC24A5
CCPG1
PYGO1
NSA2P4
KIF23
MAN2C1
IREB2
HDGFL3
ARRDC4

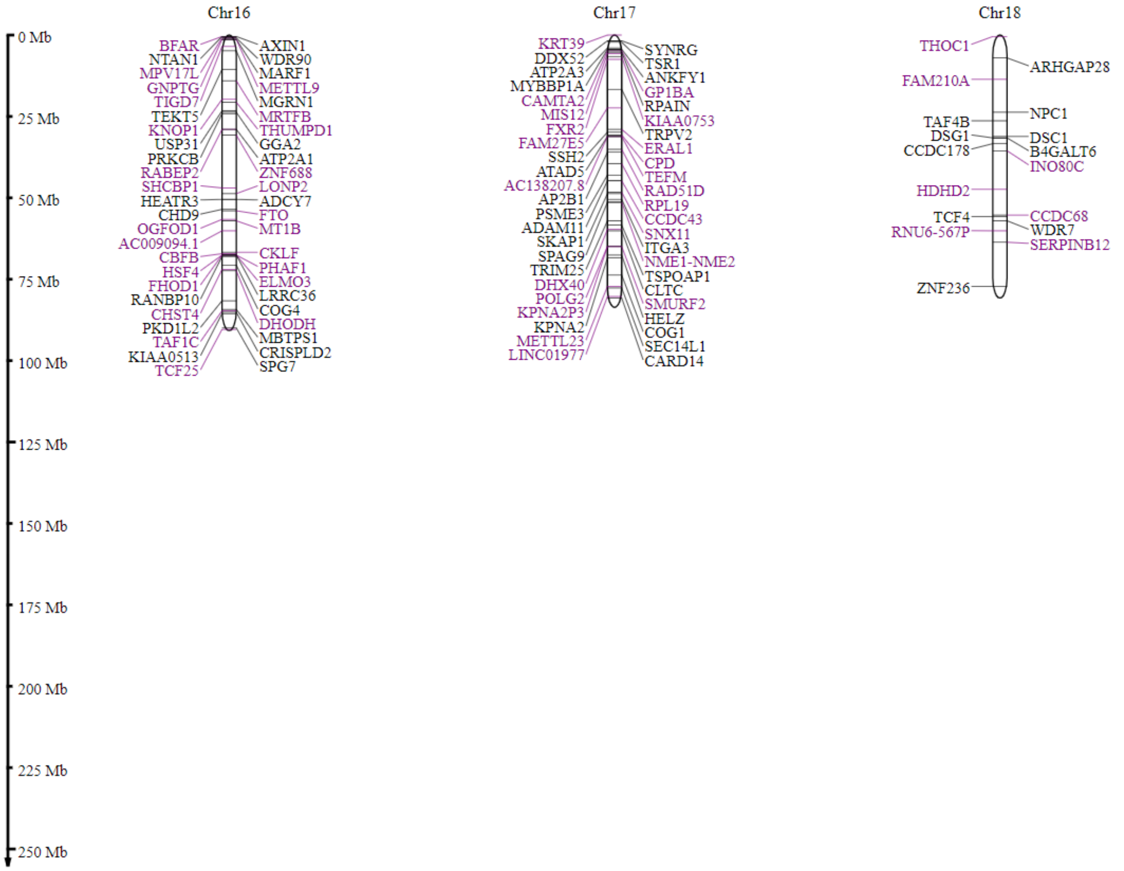
0 Mb
25 Mb
50 Mb
75 Mb
100 Mb
125 Mb
150 Mb
175 Mb
200 Mb
225 Mb
250 Mb
Chr16
BFAR
NTAN1
MPV17L
GNPTG
TIGD7
TEKT5
KNOP1
USP31
PRKCB
RABEP2
SHCBP1
HEATR3
CHD9
OGFOD1
AC009094.1
CBFB
HSF4
FHOD1
RANBP10
CHST4
PKD1L2
TAF1C
KIAA0513
TCF25
AXIN1
WDR90
MARF1
METTL9
MGRN1
MRTFB
THUMPD1
GGA2
ATP2A1
ZNF688
LONP2
ADCY7
FTO
MT1B
CKLF
PHAF1
ELMO3
LRRC36
COG4
DHODH
MBTPS1
CRISPLD2
SPG7
Chr17
KRT39
DDX52
ATP2A3
MYBBP1A
CAMTA2
MIS12
FXR2
FAM27E5
SSH2
ATAD5
AC138207.8
AP2B1
PSME3
ADAM11
SKAP1
SPAG9
TRIM25
DHX40
POLG2
KPNA2P3
KPNA2
METTL23
LINC01977
SYNRG
TSR1
ANKFY1
GP1BA
RPAIN
KIAA0753
TRPV2
ERAL1
CPD
TEFM
RAD51D
RPL19
CCDC43
SNX11
ITGA3
NME1-NME2
TSPOAP1
CLTC
SMURF2
HELZ
COG1
SEC14L1
CARD14
Chr18
THOC1
FAM210A
TAF4B
DSG1
CCDC178
HDHD2
TCF4
RNU6-567P
ZNF236
ARHGAP28
NPC1
DSC1
B4GALT6
INO80C
CCDC68
WDR7
SERPINB12

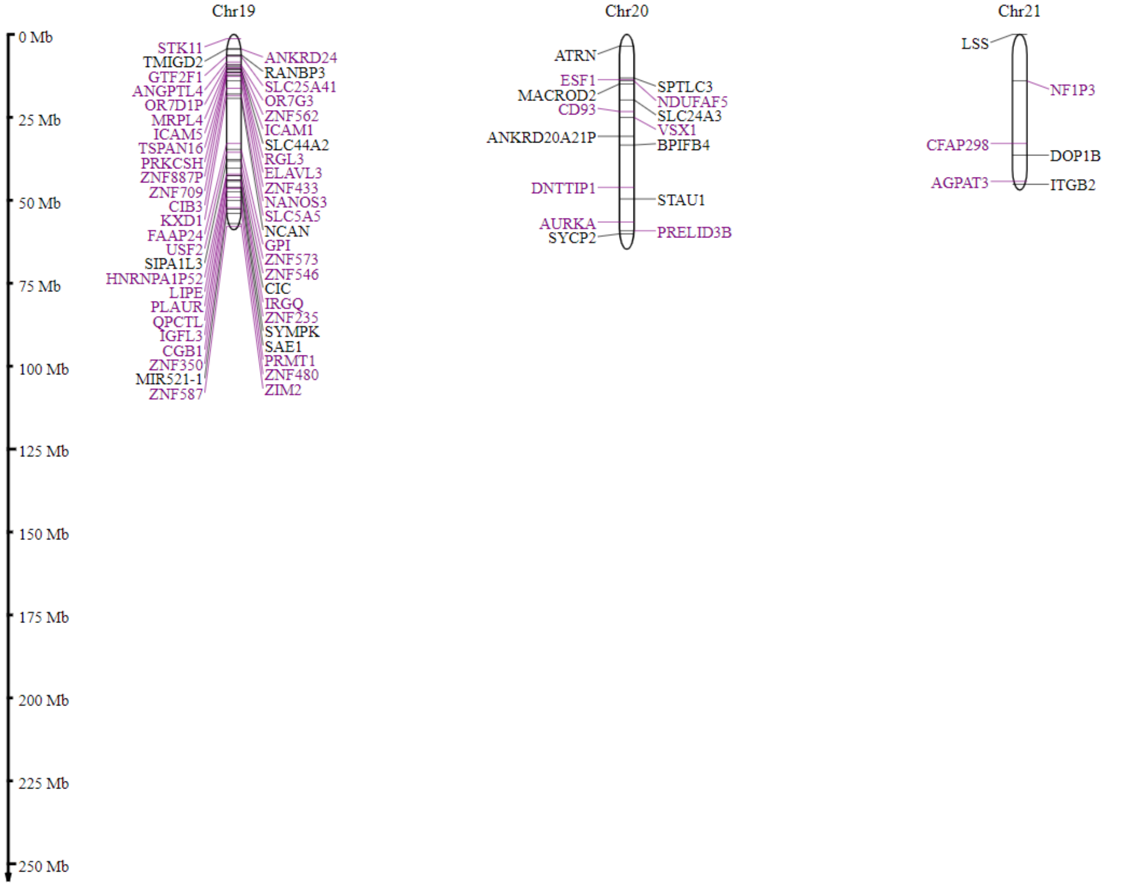
Chr19
Chr20
Chr21
0 Mb
25 Mb
50 Mb
75 Mb
100 Mb
125 Mb
150 Mb
175 Mb
200 Mb
225 Mb
250 Mb
STK11
TMIGD2
GTF2F1
ANGPTL4
OR7D1P
MRPL4
ICAM5
TSPAN16
PRKCSH
ZNF887P
ZNF709
CIB3
KXD1
FAAP24
USF2
SIPA1L3
HNRNPA1P52
LIPE
PLAUR
QPCTL
IGFL3
CGB1
ZNF350
MIR521-1
ZNF587
ANKRD24
RANBP3
SLC25A41
OR7G3
ZNF562
ICAM1
SLC44A2
RGL3
ELAVL3
ZNF433
NANOS3
SLC5A5
NCAN
GPI
ZNF573
ZNF546
CIC
IRGQ
ZNF235
SYMPK
SAE1
PRMT1
ZNF480
ZIM2
ATRN
ESF1
MACROD2
CD93
ANKRD20A21P
DNTTIP1
AURKA
SYCP2
SPTLC3
NDUFAF5
SLC24A3
VSX1
BPIFB4
STAU1
PRELID3B
LSS
NF1P3
CFAP298
DOP1B
AGPAT3
ITGB2

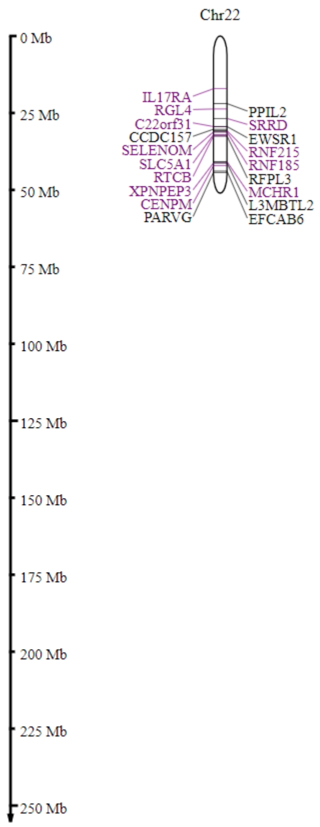
Chr22
0 Mb
25 Mb
50 Mb
75 Mb
100 Mb
125 Mb
150 Mb
175 Mb
200 Mb
225 Mb
250 Mb
IL17RA
RGL4
C22orf31
CCDC157
SELENOM
SLC5A1
RTCB
XPNPEP3
CENPM
PARVG
PPIL2
SRRD
EWSR1
RNF215
RNF185
RFPL3
MCHR1
L3MBTL2
EFCAB6

**Fig. S2:** Autosomal scatter of the sexually dimorphic genes**.** Biased and unbiased autosomal genes by chromosomal genomic location.

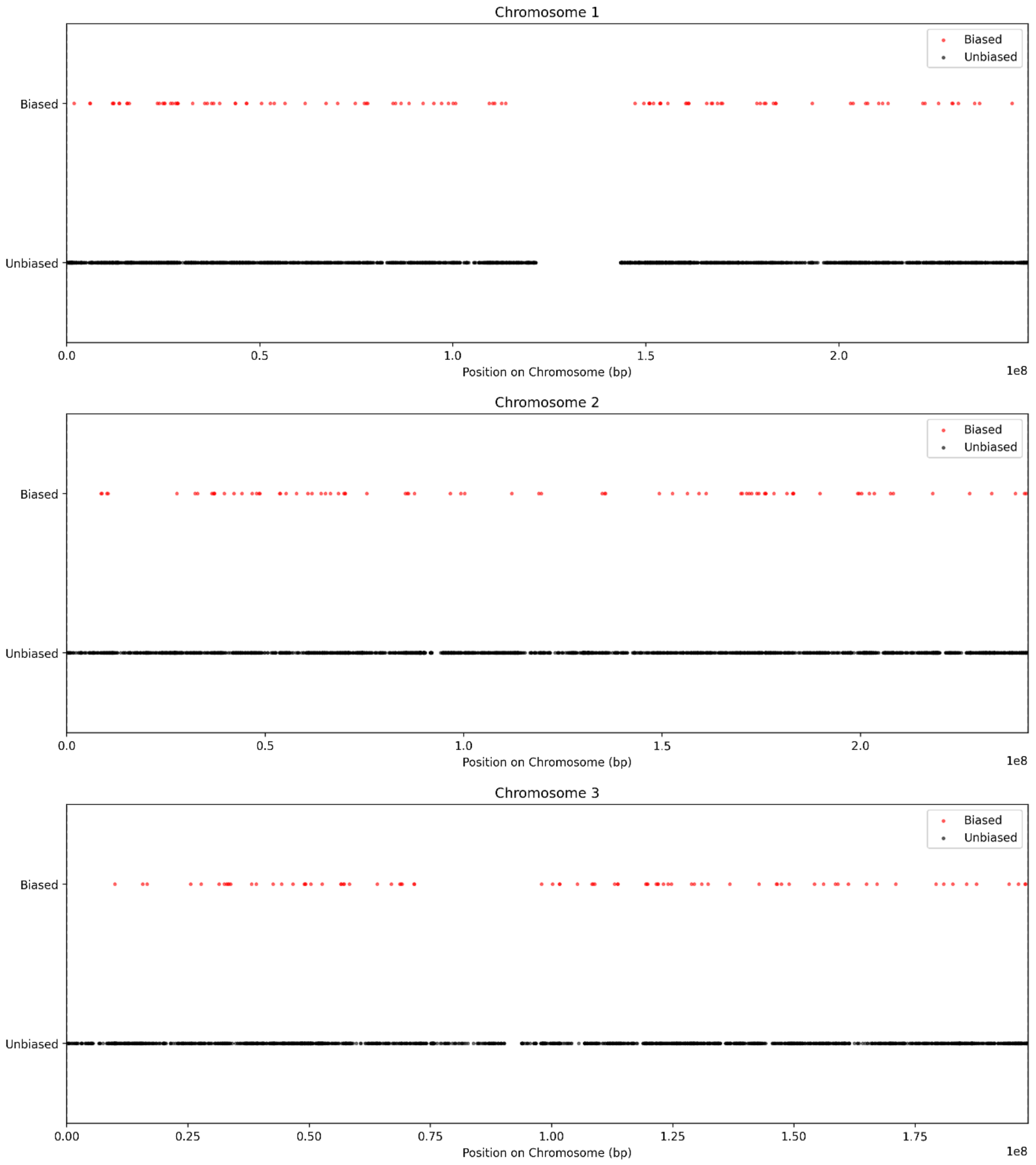

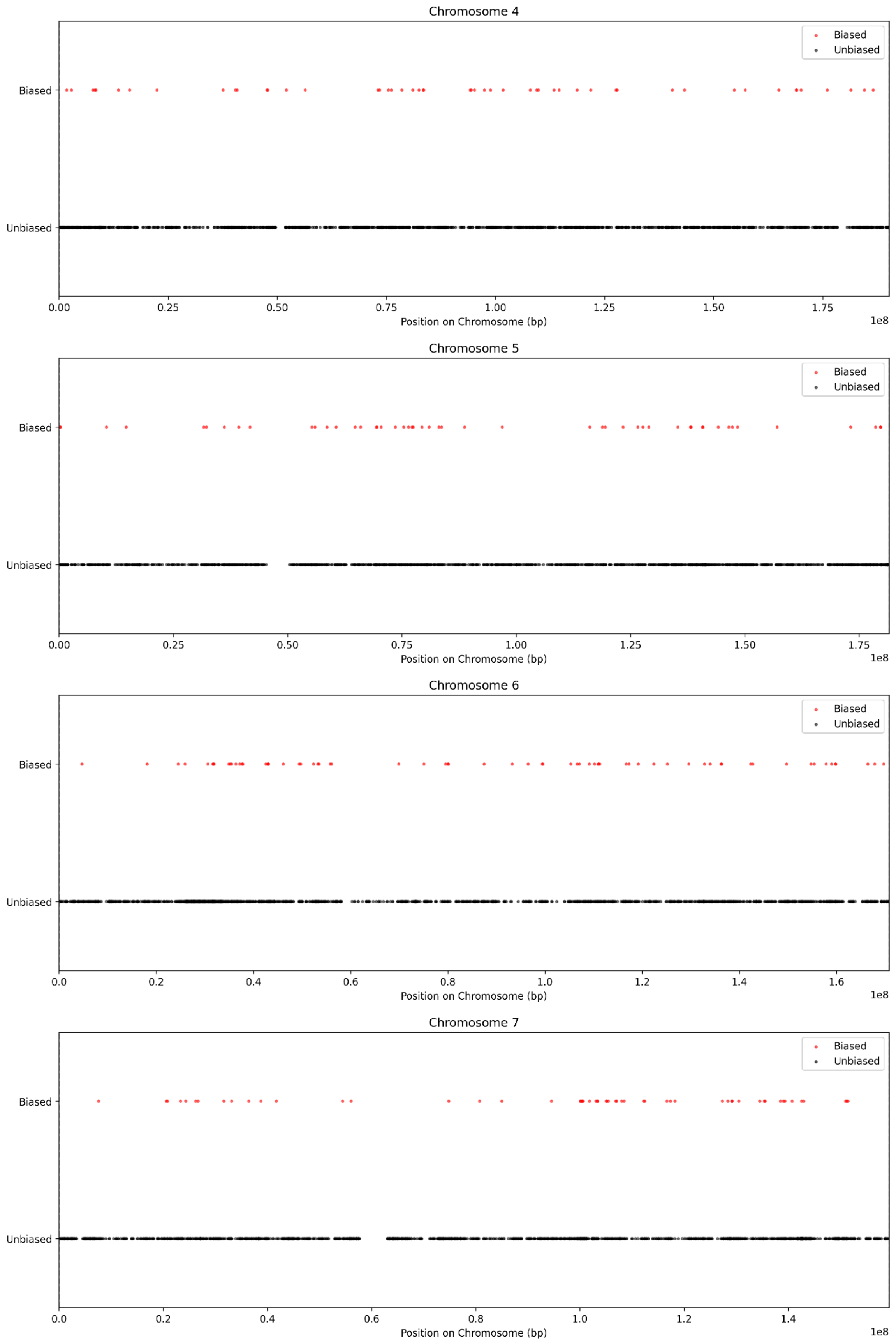
Chromosome 4
Biased
Unbiased
0.00
0.25
0.50
0.75
1.00
1.25
1.50
1.75
1e8
Position on Chromosome (bp)
Chromosome 5
Biased
Unbiased
0.00
0.25
0.50
0.75
1.00
1.25
1.50
1.75
1e8
Position on Chromosome (bp)
Chromosome 6
Biased
Unbiased
0.0
0.2
0.4
0.6
0.8
1.0
1.2
1.4
1.6
1e8
Position on Chromosome (bp)
Chromosome 7
Biased
Unbiased
0.0
0.2
0.4
0.6
0.8
1.0
1.2
1.4
1e8
Position on Chromosome (bp)

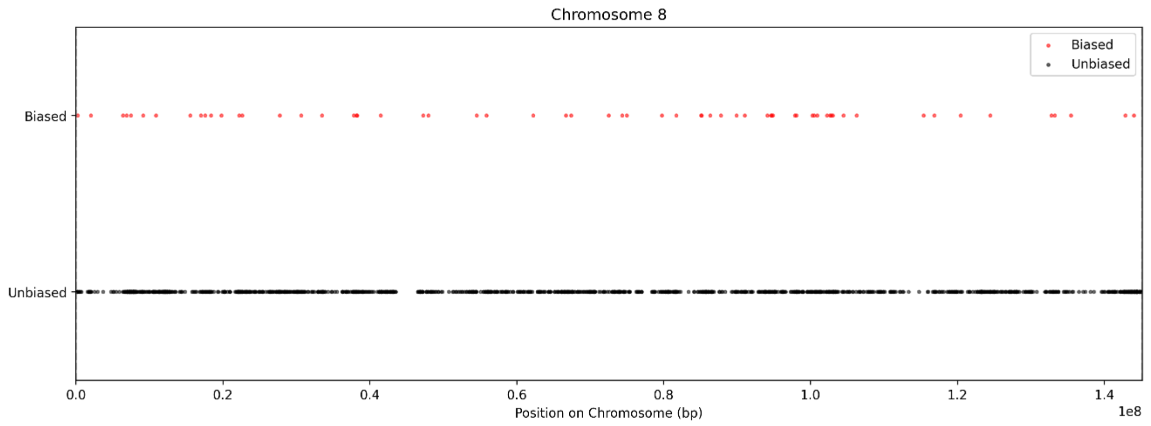
Chromosome 8
Biased
Unbiased
Biased
Unbiased
0.0
0.2
0.4
0.6
0.8
1.0
1.2
1.4
Position on Chromosome (bp)
1e8

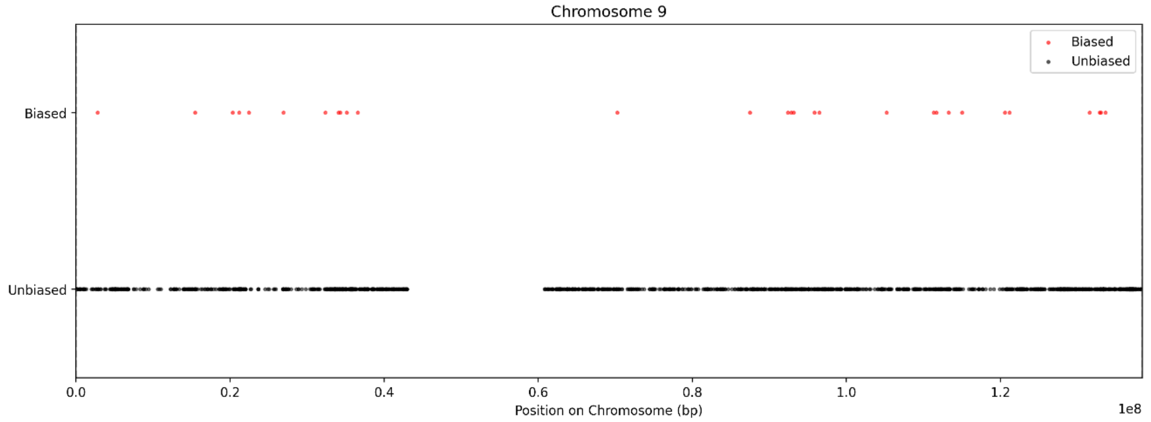
Chromosome 9
Biased
Unbiased
Biased
Unbiased
0.0
0.2
0.4
0.6
0.8
1.0
1.2
Position on Chromosome (bp)
1e8

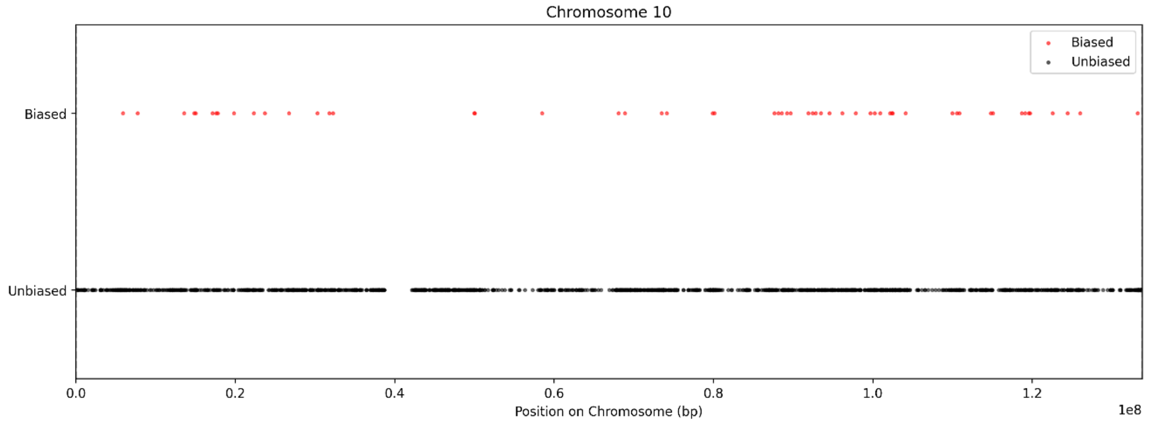
Chromosome 10
Biased
Unbiased
Biased
Unbiased
0.0
0.2
0.4
0.6
0.8
1.0
1.2
Position on Chromosome (bp)
1e8

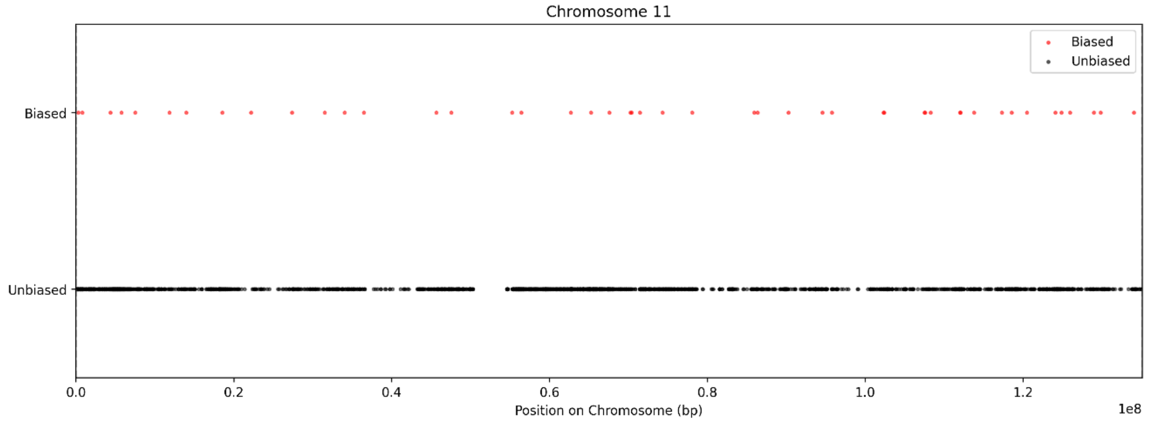
Chromosome 11
Biased
Unbiased
Biased
Unbiased
0.0
0.2
0.4
0.6
0.8
1.0
1.2
Position on Chromosome (bp)
1e8

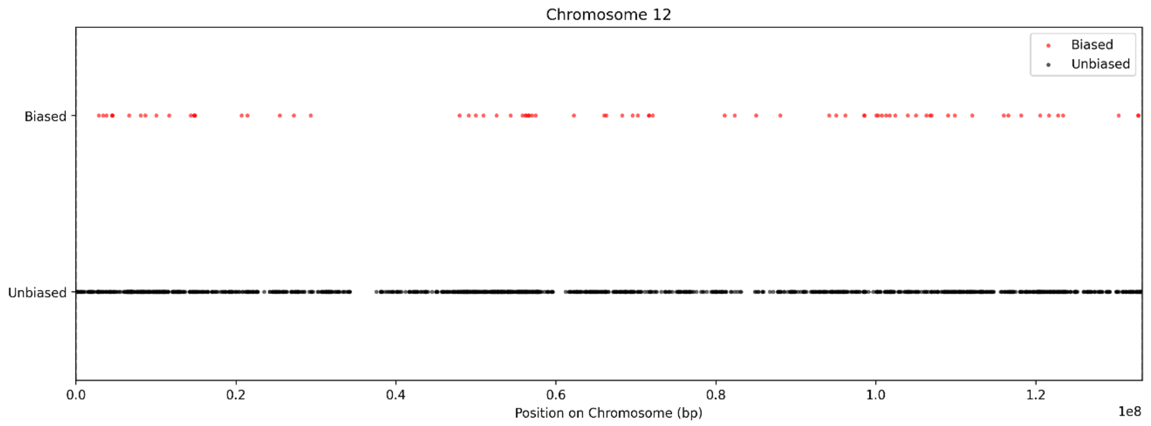
Chromosome 12
Biased
Unbiased
Biased
Unbiased
0.0
0.2
0.4
0.6
0.8
1.0
1.2
1e8
Position on Chromosome (bp)

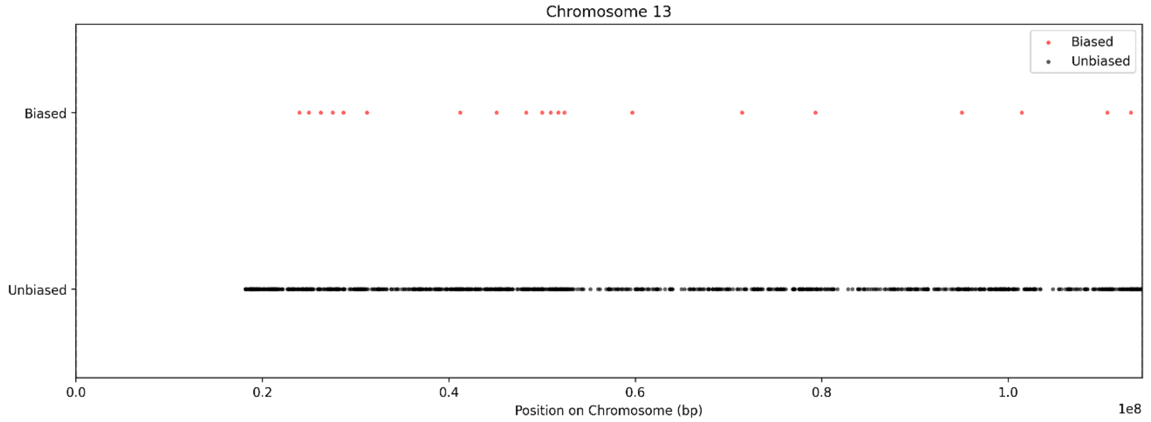
Chromosome 13
Biased
Unbiased
Biased
Unbiased
0.0
0.2
0.4
0.6
0.8
1.0
1e8
Position on Chromosome (bp)

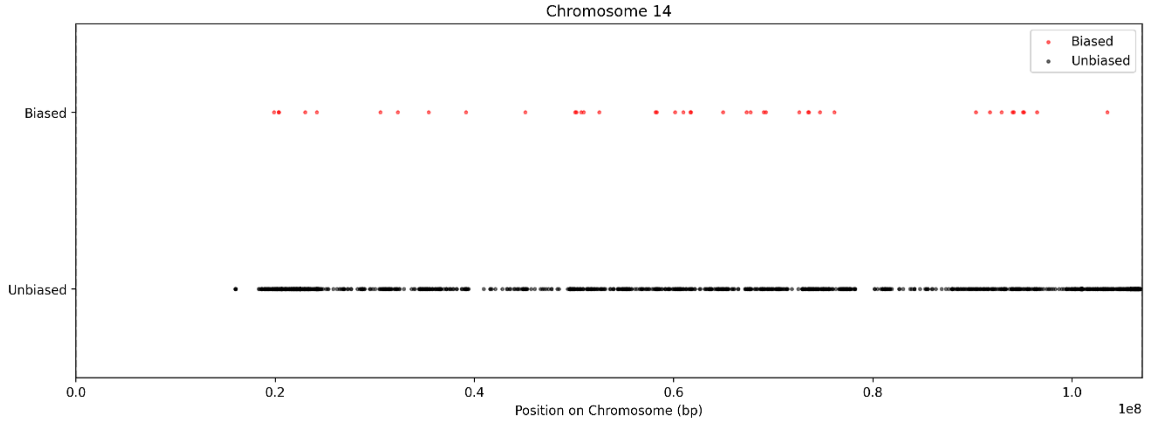
Chromosome 14
Biased
Unbiased
Biased
Unbiased
0.0
0.2
0.4
0.6
0.8
1.0
1e8
Position on Chromosome (bp)

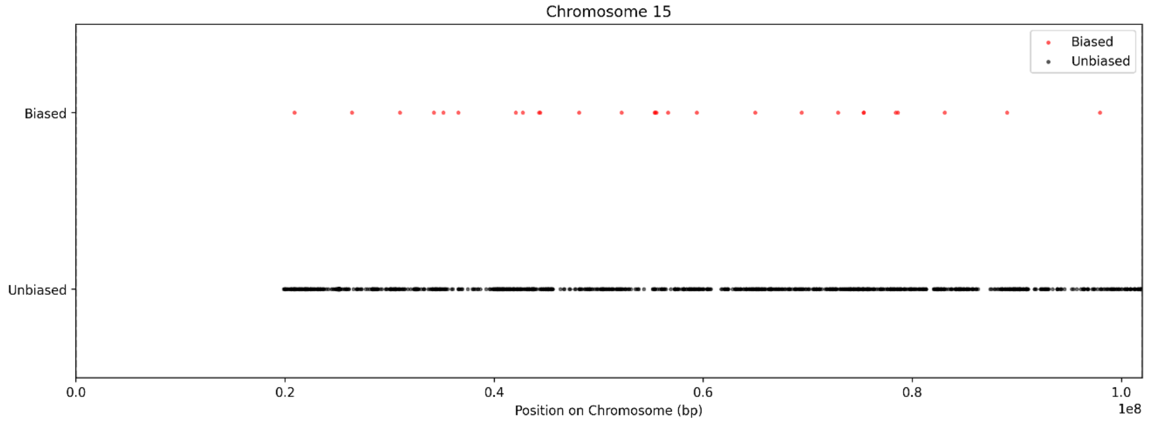
Chromosome 15
Biased
Unbiased
Biased
Unbiased
0.0
0.2
0.4
0.6
0.8
1.0
1e8
Position on Chromosome (bp)

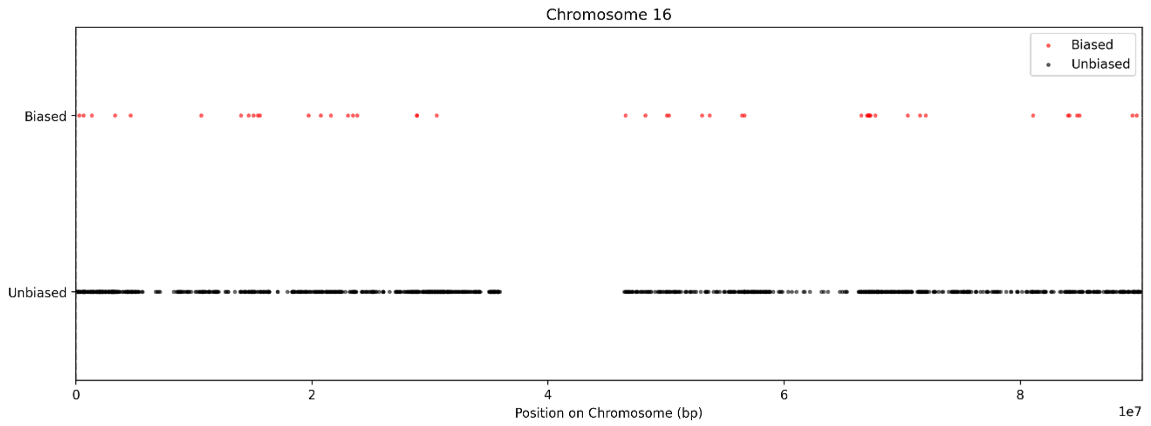
Chromosome 16
Biased
Unbiased
Biased
Unbiased
0
2
4
6
8
1e7
Position on Chromosome (bp)

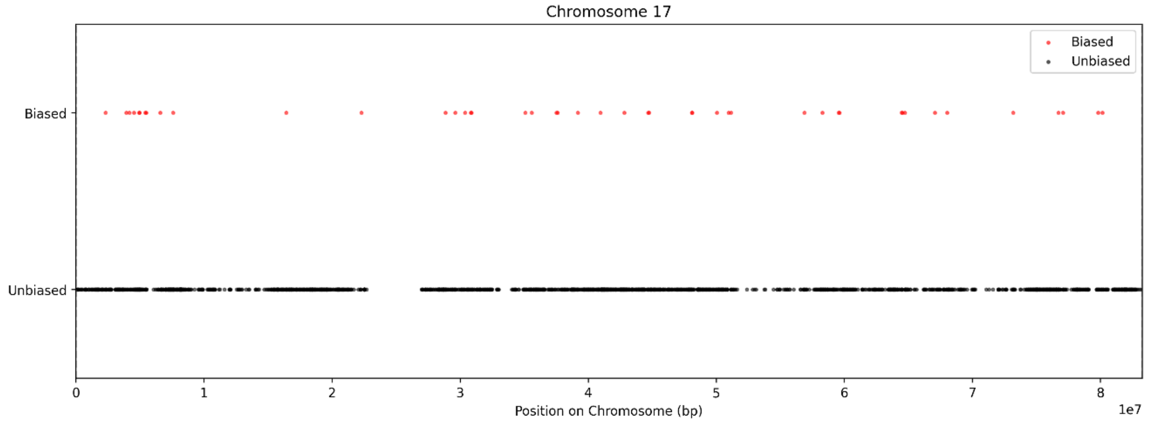
Chromosome 17
Biased
Unbiased
Biased
Unbiased
0
1
2
3
4
5
6
7
8
1e7
Position on Chromosome (bp)

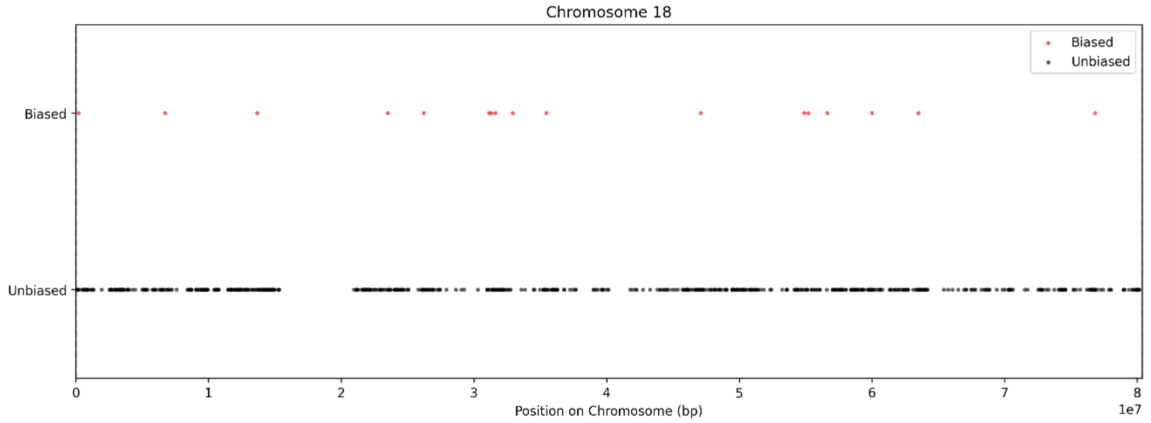
Chromosome 18
Biased
Unbiased
Biased
Unbiased
0
1
2
3
4
5
6
7
8
1e7
Position on Chromosome (bp)

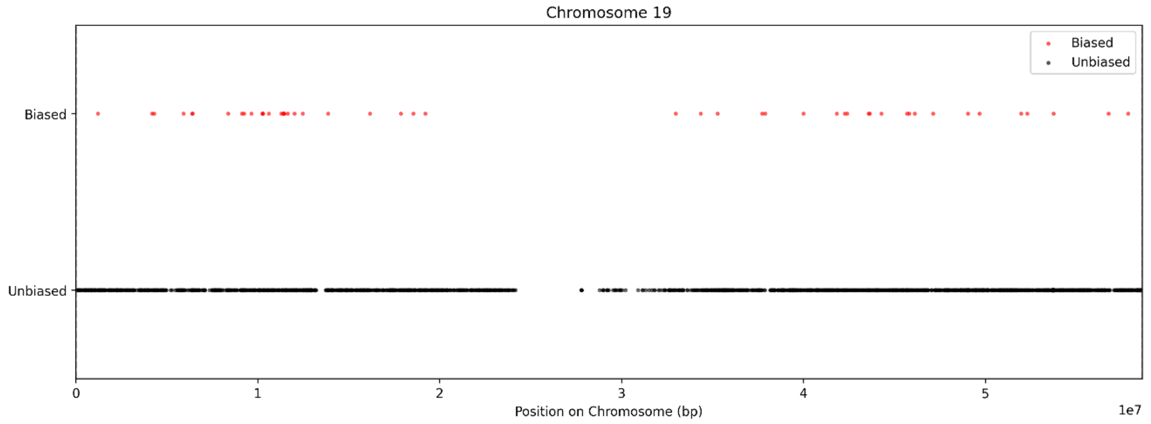
Chromosome 19
Biased
Unbiased
Biased
Unbiased
0
1
2
3
4
5
1e7
Position on Chromosome (bp)

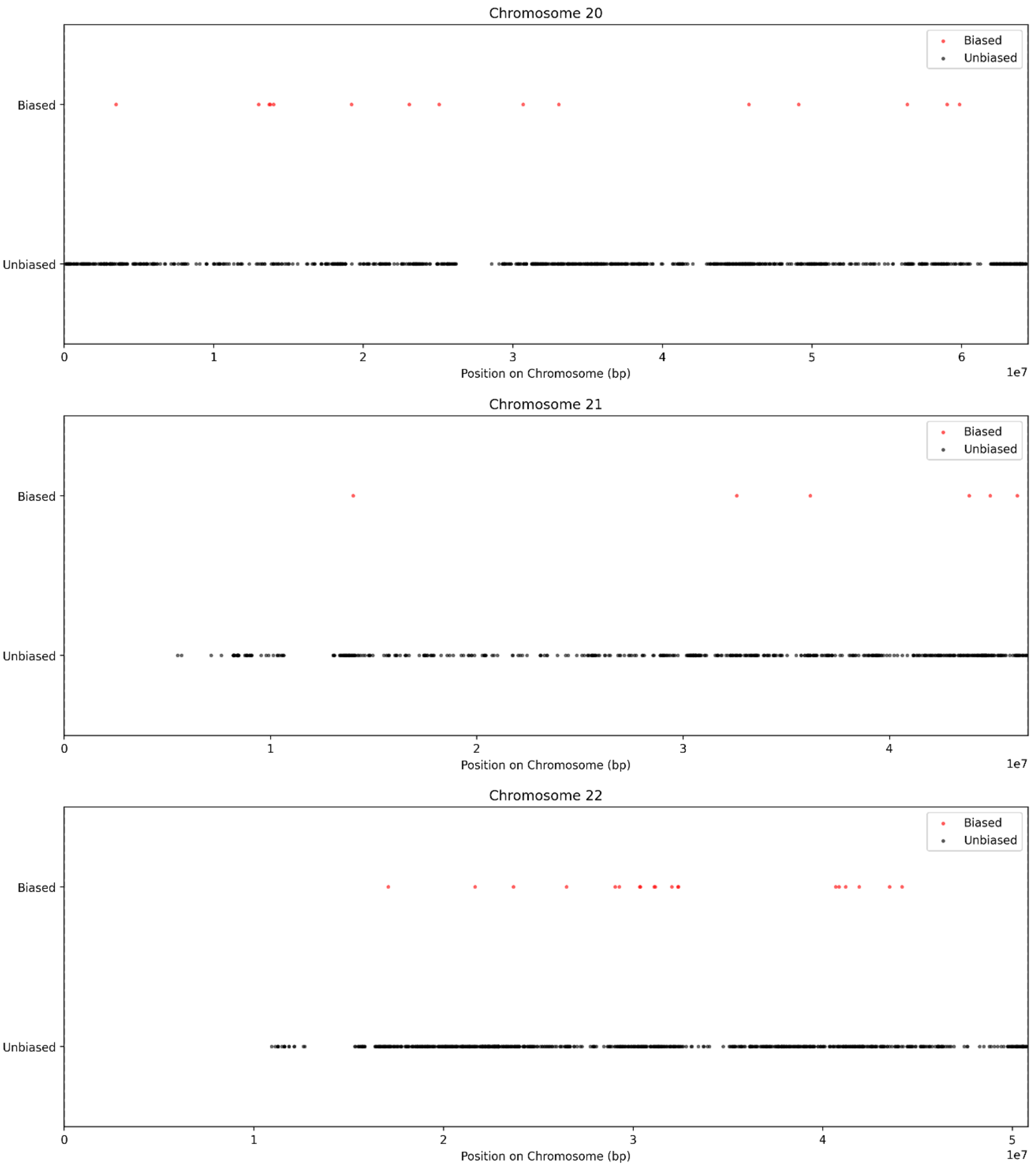

Chromosome 20
Biased
Unbiased
Biased
Unbiased
0
1
2
3
4
5
6
1e7
Position on Chromosome (bp)
Chromosome 21
Biased
Unbiased
Biased
Unbiased
0
1
2
3
4
1e7
Position on Chromosome (bp)
Chromosome 22
Biased
Unbiased
Biased
Unbiased
0
1
2
3
4
5
1e7
Position on Chromosome (bp)

**Fig, S3:** Performance of the Random Forest classifier in predicting tumor sample's sex based on autosomal gene mutation profiles**.**

**A**

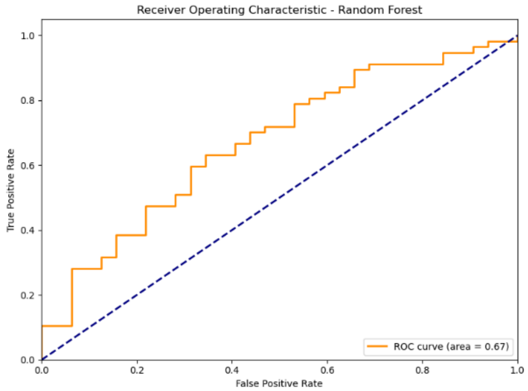

**B**

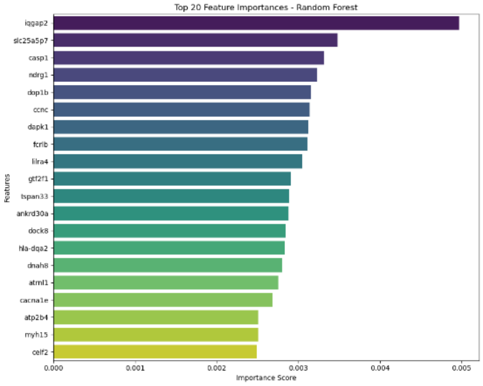

**C**

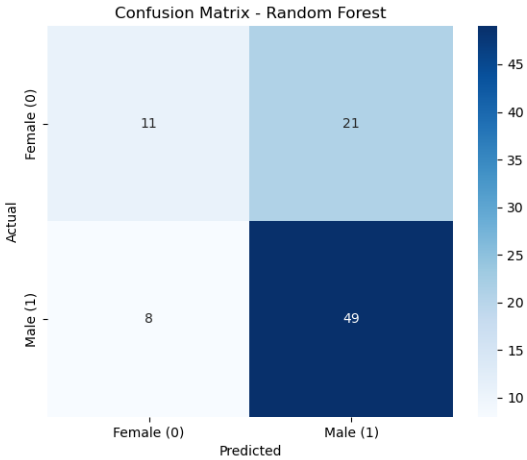

**(A)** The Receiver Operating Characteristic (ROC) curve of the Random Forest model evaluated on the test set, illustrating its ability to distinguish between male and female gastric cancer samples, with an area under the curve (AUC) of 0.7262. **(B)** Feature importance plot displaying the top 20 genes ranked by their importance scores in the Random Forest model. **(C)** Confusion matrix summarizing the Random Forest model classification results on the test set.

**Table S1**: 1052 sexually dimorphic mutated genes

| Gene Symbol | Chr | p Value | Corrected p value |
|---|---|---|---|
| B3GLCT | 13 | 8.347E-08 | 0.002016 |
| NDUFAF7 | 2 | 1.229E-07 | 0.002016 |
| CWC27 | 5 | 8.789E-07 | 0.005829 |
| OMD | 9 | 9.149E-07 | 0.005829 |
| E2F5 | 8 | 9.226E-07 | 0.005829 |
| PPP4R4 | 14 | 1.066E-06 | 0.005829 |
| TRMT44 | 4 | 1.407E-06 | 0.006063 |
| SMARCAD1 | 4 | 1.505E-06 | 0.006063 |
| GTF2F1 | 19 | 1.663E-06 | 0.006063 |
| PARP11 | 12 | 2.026E-06 | 0.006649 |
| BLNK | 10 | 2.472E-06 | 0.007361 |
| ZNF480 | 19 | 2.919E-06 | 0.007361 |
| LGR4 | 11 | 2.986E-06 | 0.007361 |
| NDRG1 | 8 | 3.172E-06 | 0.007361 |
| RPL23AP96 | 8 | 3.365E-06 | 0.007361 |
| ELAVL3 | 19 | 3.65E-06 | 0.007486 |
| DUSP19 | 2 | 4.536E-06 | 0.008318 |
| MMUT | 6 | 4.727E-06 | 0.008318 |
| WNK2 | 9 | 5.062E-06 | 0.008318 |
| ATP6V1D | 14 | 5.323E-06 | 0.008318 |
| NUP43 | 6 | 5.323E-06 | 0.008318 |
| ZFC3H1 | 12 | 5.883E-06 | 0.008776 |
| COA8 | 14 | 6.197E-06 | 0.008842 |
| SLAMF1 | 1 | 6.674E-06 | 0.009126 |
| CEL | 9 | 7.887E-06 | 0.009268 |
| QRSL1P2 | 1 | 8.168E-06 | 0.009268 |
| ZNF709 | 19 | 8.415E-06 | 0.009268 |
| SLC5A1 | 22 | 8.486E-06 | 0.009268 |
| ZBTB8OS | 1 | 8.577E-06 | 0.009268 |
| TIAL1 | 10 | 8.704E-06 | 0.009268 |
| DTWD2 | 5 | 8.8E-06 | 0.009268 |
| MARF1 | 16 | 9.171E-06 | 0.009268 |
| FNTB | 14 | 9.507E-06 | 0.009268 |
| MYH15 | 3 | 9.602E-06 | 0.009268 |
| RHEB | 7 | 1.006E-05 | 0.009430 |
| CERT1 | 5 | 1.063E-05 | 0.009686 |
| IL17RA | 22 | 1.143E-05 | 0.009894 |
| ZNF546 | 19 | 1.146E-05 | 0.009894 |
| SCUBE3 | 6 | 1.25E-05 | 0.010369 |
| ACOT6 | 14 | 1.264E-05 | 0.010369 |
| TANK | 2 | 1.408E-05 | 0.010513 |
| PHF10 | 6 | 1.408E-05 | 0.010513 |
| HELB | 12 | 1.47E-05 | 0.010513 |

| | | | |
|---|---|---|---|
| CLCA4 | 1 | 1.502E-05 | 0.010513 |
| PRKAR2B | 7 | 1.502E-05 | 0.010513 |
| TTC13 | 1 | 1.518E-05 | 0.010513 |
| EPC1 | 10 | 1.547E-05 | 0.010513 |
| FXR2 | 17 | 1.62E-05 | 0.010513 |
| STK38L | 12 | 1.642E-05 | 0.010513 |
| TMEM106C | 12 | 1.654E-05 | 0.010513 |
| SLC2A2 | 3 | 1.654E-05 | 0.010513 |
| DOP1B | 21 | 1.691E-05 | 0.010513 |
| CCNA2 | 4 | 1.723E-05 | 0.010513 |
| GTF2F2 | 13 | 1.8E-05 | 0.010513 |
| SGK3 | 8 | 1.874E-05 | 0.010513 |
| GRIA2 | 4 | 1.895E-05 | 0.010513 |
| HAPLN1 | 5 | 1.937E-05 | 0.010513 |
| NUDT2 | 9 | 1.937E-05 | 0.010513 |
| ELMOD3 | 2 | 1.972E-05 | 0.010513 |
| DMRTA1 | 9 | 1.972E-05 | 0.010513 |
| CYP11B1 | 8 | 1.994E-05 | 0.010513 |
| ABCC4 | 13 | 2.012E-05 | 0.010513 |
| SPTLC3 | 20 | 2.108E-05 | 0.010513 |
| AC027288.2 | 12 | 2.119E-05 | 0.010513 |
| ST13P19 | 1 | 2.136E-05 | 0.010513 |
| ARCN1 | 11 | 2.136E-05 | 0.010513 |
| COL19A1 | 6 | 2.178E-05 | 0.010513 |
| ROBO4 | 11 | 2.244E-05 | 0.010513 |
| RIOX1 | 14 | 2.264E-05 | 0.010513 |
| CAMTA2 | 17 | 2.27E-05 | 0.010513 |
| RUNX3 | 1 | 2.275E-05 | 0.010513 |
| ZNF484 | 9 | 2.53E-05 | 0.011452 |
| AC083923.1 | 8 | 2.639E-05 | 0.011452 |
| NUP54 | 4 | 2.64E-05 | 0.011452 |
| AL356473.1 | 6 | 2.726E-05 | 0.011452 |
| PAPOLA | 14 | 2.765E-05 | 0.011452 |
| ERBIN | 5 | 2.765E-05 | 0.011452 |
| MDM1 | 12 | 2.781E-05 | 0.011452 |
| PROZ | 13 | 2.783E-05 | 0.011452 |
| MAN1A1 | 6 | 2.792E-05 | 0.011452 |
| FBLN7 | 2 | 2.83E-05 | 0.011467 |
| CEBPZ | 2 | 2.894E-05 | 0.011583 |
| RIC8B | 12 | 2.989E-05 | 0.011599 |
| LNX2 | 13 | 2.996E-05 | 0.011599 |
| MYO5C | 15 | 3.005E-05 | 0.011599 |
| NSA2P4 | 15 | 3.096E-05 | 0.011599 |
| RB1 | 13 | 3.122E-05 | 0.011599 |
| GALNT16 | 14 | 3.165E-05 | 0.011599 |
| FAT1 | 4 | 3.19E-05 | 0.011599 |
| MARCO | 2 | 3.263E-05 | 0.011599 |
| PLPP3 | 1 | 3.274E-05 | 0.011599 |

| | | | |
|---|---|---|---|
| PHAF1 | 16 | 3.274E-05 | 0.011599 |
| QRICH1 | 3 | 3.296E-05 | 0.011599 |
| OXSR1 | 3 | 3.324E-05 | 0.011599 |
| HELLS | 10 | 3.358E-05 | 0.011599 |
| UBR5 | 8 | 3.484E-05 | 0.011721 |
| CENPM | 22 | 3.528E-05 | 0.011721 |
| RPP14 | 3 | 3.565E-05 | 0.011721 |
| TRIM48 | 11 | 3.572E-05 | 0.011721 |
| SMURF2 | 17 | 3.572E-05 | 0.011721 |
| LLCFC1 | 7 | 3.671E-05 | 0.011773 |
| FUBP1 | 1 | 3.747E-05 | 0.011773 |
| RIMKLB | 12 | 3.953E-05 | 0.011773 |
| USP47 | 11 | 4.045E-05 | 0.011773 |
| CDON | 11 | 4.114E-05 | 0.011773 |
| TAPT1 | 4 | 4.128E-05 | 0.011773 |
| SH3BGRL2 | 6 | 4.128E-05 | 0.011773 |
| SHOC1 | 9 | 4.13E-05 | 0.011773 |
| NKAPD1 | 11 | 4.132E-05 | 0.011773 |
| SCYL2 | 12 | 4.149E-05 | 0.011773 |
| PMS1 | 2 | 4.149E-05 | 0.011773 |
| DEPDC1B | 5 | 4.149E-05 | 0.011773 |
| FEZ2 | 2 | 4.225E-05 | 0.011773 |
| DDI2 | 1 | 4.233E-05 | 0.011773 |
| MRTFB | 16 | 4.233E-05 | 0.011773 |
| RTCB | 22 | 4.233E-05 | 0.011773 |
| GAB1 | 4 | 4.233E-05 | 0.011773 |
| C9 | 5 | 4.233E-05 | 0.011773 |
| HAT1 | 2 | 4.349E-05 | 0.011994 |
| AFM | 4 | 4.521E-05 | 0.012345 |
| IDE | 10 | 4.575E-05 | 0.012345 |
| SRSF10 | 1 | 4.614E-05 | 0.012345 |
| PSIP1 | 9 | 4.627E-05 | 0.012345 |
| PLXDC2 | 10 | 4.744E-05 | 0.012408 |
| TMPO | 12 | 4.862E-05 | 0.012408 |
| PARP2 | 14 | 4.862E-05 | 0.012408 |
| CPD | 17 | 4.927E-05 | 0.012408 |
| UEVLD | 11 | 4.928E-05 | 0.012408 |
| NAA25 | 12 | 4.928E-05 | 0.012408 |
| AGPAT3 | 21 | 4.934E-05 | 0.012408 |
| PRMT1 | 19 | 4.953E-05 | 0.012408 |
| ESCO2 | 8 | 5.044E-05 | 0.012539 |
| TIGD7 | 16 | 5.132E-05 | 0.012663 |
| PELI1 | 2 | 5.388E-05 | 0.013161 |
| GOLM2 | 15 | 5.459E-05 | 0.013161 |
| USF2 | 19 | 5.459E-05 | 0.013161 |
| SLC2A12 | 6 | 5.494E-05 | 0.013161 |
| SOS2 | 14 | 5.573E-05 | 0.013169 |
| RSRP1 | 1 | 5.624E-05 | 0.013169 |

| | | | |
|---|---|---|---|
| ICAM1 | 19 | 5.624E-05 | 0.013169 |
| TAOK3 | 12 | 5.672E-05 | 0.013169 |
| TTBK2 | 15 | 5.723E-05 | 0.013169 |
| IRGQ | 19 | 5.739E-05 | 0.013169 |
| RBM26 | 13 | 5.807E-05 | 0.013201 |
| ZNF573 | 19 | 5.833E-05 | 0.013201 |
| DPH6 | 15 | 5.976E-05 | 0.013432 |
| SPIDR | 8 | 6.249E-05 | 0.013951 |
| PREP | 6 | 6.429E-05 | 0.014255 |
| ADCY7 | 16 | 6.637E-05 | 0.014570 |
| INO80C | 18 | 6.659E-05 | 0.014570 |
| SYCP2 | 20 | 6.771E-05 | 0.014716 |
| INPP5F | 10 | 6.982E-05 | 0.014963 |
| CAMK2D | 4 | 7.066E-05 | 0.014963 |
| LINC02365 | 4 | 7.108E-05 | 0.014963 |
| ITGBL1 | 13 | 7.113E-05 | 0.014963 |
| NKTR | 3 | 7.113E-05 | 0.014963 |
| OTUD6B | 8 | 7.241E-05 | 0.015135 |
| BOD1L1 | 4 | 7.528E-05 | 0.015310 |
| AHRR | 5 | 7.544E-05 | 0.015310 |
| RPF1 | 1 | 7.556E-05 | 0.015310 |
| KHDRBS3 | 8 | 7.556E-05 | 0.015310 |
| LONP2 | 16 | 7.604E-05 | 0.015310 |
| CRISP3 | 6 | 7.604E-05 | 0.015310 |
| ZZZ3 | 1 | 7.797E-05 | 0.015507 |
| NR2C1 | 12 | 7.797E-05 | 0.015507 |
| HELZ | 17 | 7.901E-05 | 0.015526 |
| LNPEP | 5 | 7.901E-05 | 0.015526 |
| CNOT4 | 7 | 8.031E-05 | 0.015648 |
| RILPL2 | 12 | 8.076E-05 | 0.015648 |
| BMS1P21 | 10 | 8.106E-05 | 0.015648 |
| CCNC | 6 | 8.28E-05 | 0.015824 |
| MTPAP | 10 | 8.339E-05 | 0.015824 |
| CCNT2 | 2 | 8.342E-05 | 0.015824 |
| RP1 | 8 | 8.533E-05 | 0.016007 |
| SERPINB12 | 18 | 8.608E-05 | 0.016007 |
| NFXL1 | 4 | 8.608E-05 | 0.016007 |
| ADGRA3 | 4 | 8.666E-05 | 0.016007 |
| RHNO1 | 12 | 8.723E-05 | 0.016007 |
| PDE8B | 5 | 8.731E-05 | 0.016007 |
| PIGK | 1 | 8.863E-05 | 0.016064 |
| GPATCH2L | 14 | 8.863E-05 | 0.016064 |
| AADAT | 4 | 8.909E-05 | 0.016064 |
| ID2 | 2 | 9.047E-05 | 0.016198 |
| SLC46A3 | 13 | 9.082E-05 | 0.016198 |
| CDIN1 | 15 | 9.157E-05 | 0.016244 |
| MPV17L | 16 | 9.444E-05 | 0.016551 |
| SOCS5 | 2 | 9.447E-05 | 0.016551 |

| | | | |
|---|---|---|---|
| DNAAF4 | 15 | 9.546E-05 | 0.016551 |
| PKN2 | 1 | 9.583E-05 | 0.016551 |
| SPG7 | 16 | 9.583E-05 | 0.016551 |
| GATAD2B | 1 | 9.823E-05 | 0.016573 |
| OR4K2 | 14 | 9.823E-05 | 0.016573 |
| SLC38A6 | 14 | 9.823E-05 | 0.016573 |
| METTL8 | 2 | 9.823E-05 | 0.016573 |
| KPNA2 | 17 | 9.926E-05 | 0.016573 |
| NPY | 7 | 0.000101 | 0.016573 |
| RAPGEF1 | 9 | 0.0001017 | 0.016573 |
| PHF20L1 | 8 | 0.000102 | 0.016573 |
| SYDE2 | 1 | 0.0001027 | 0.016573 |
| ATP5F1B | 12 | 0.0001027 | 0.016573 |
| THUMPD2 | 2 | 0.0001027 | 0.016573 |
| ARMC10 | 7 | 0.0001031 | 0.016573 |
| MAP7 | 6 | 0.0001032 | 0.016573 |
| COX20 | 1 | 0.0001035 | 0.016573 |
| RPL21P18 | 12 | 0.0001035 | 0.016573 |
| MIS12 | 17 | 0.0001075 | 0.017036 |
| IGFL3 | 19 | 0.0001075 | 0.017036 |
| C22orf31 | 22 | 0.0001113 | 0.017564 |
| R3HDM1 | 2 | 0.0001133 | 0.017698 |
| ZNF277 | 7 | 0.0001133 | 0.017698 |
| RPAP2 | 1 | 0.0001155 | 0.017781 |
| ASB3 | 2 | 0.0001155 | 0.017781 |
| TTI2 | 8 | 0.0001155 | 0.017781 |
| WWP1 | 8 | 0.0001159 | 0.017781 |
| MEGF9 | 9 | 0.0001181 | 0.017787 |
| DHX29 | 5 | 0.000119 | 0.017787 |
| ITGA3 | 17 | 0.0001198 | 0.017787 |
| B4GALT6 | 18 | 0.0001198 | 0.017787 |
| FAM161A | 2 | 0.0001198 | 0.017787 |
| BRAF | 7 | 0.0001198 | 0.017787 |
| CDC40 | 6 | 0.0001212 | 0.017787 |
| HADH | 4 | 0.0001218 | 0.017787 |
| SLC26A5 | 7 | 0.0001218 | 0.017787 |
| FCF1 | 14 | 0.0001218 | 0.017787 |
| MELK | 9 | 0.0001224 | 0.017787 |
| ODC1 | 2 | 0.0001225 | 0.017787 |
| RAD17 | 5 | 0.0001236 | 0.017862 |
| RALGPS2 | 1 | 0.0001257 | 0.018092 |
| MSH5 | 6 | 0.000128 | 0.018344 |
| MYOM2 | 8 | 0.0001313 | 0.018737 |
| TMEM123 | 11 | 0.0001355 | 0.019255 |
| THRAP3P1 | 3 | 0.0001378 | 0.019487 |
| SLCO1C1 | 12 | 0.00014 | 0.019567 |
| MARCHF6 | 5 | 0.00014 | 0.019567 |
| RCOR3 | 1 | 0.0001406 | 0.019567 |

| | | | |
|---|---|---|---|
| QPCTL | 19 | 0.0001418 | 0.019567 |
| ADAM12 | 10 | 0.0001432 | 0.019567 |
| BIRC6 | 2 | 0.0001433 | 0.019567 |
| TUSC3 | 8 | 0.0001442 | 0.019567 |
| LHCGR | 2 | 0.0001466 | 0.019567 |
| DCAF5 | 14 | 0.000147 | 0.019567 |
| BCKDHB | 6 | 0.000147 | 0.019567 |
| VPS36 | 13 | 0.0001476 | 0.019567 |
| HAVCR1 | 5 | 0.0001481 | 0.019567 |
| BLTP3B | 12 | 0.0001482 | 0.019567 |
| EML4 | 2 | 0.0001482 | 0.019567 |
| STAU1 | 20 | 0.0001482 | 0.019567 |
| MCPH1 | 8 | 0.0001482 | 0.019567 |
| PRDM2 | 1 | 0.0001485 | 0.019567 |
| RNU6-567P | 18 | 0.00015 | 0.019688 |
| HSF4 | 16 | 0.0001552 | 0.020088 |
| LIPE | 19 | 0.0001552 | 0.020088 |
| AP2B1 | 17 | 0.0001555 | 0.020088 |
| LRCH3 | 3 | 0.0001555 | 0.020088 |
| SHCBP1 | 16 | 0.0001588 | 0.020355 |
| RAPH1 | 2 | 0.0001588 | 0.020355 |
| TRHDE | 12 | 0.0001609 | 0.020498 |
| SI | 3 | 0.0001612 | 0.020498 |
| RO60 | 1 | 0.0001646 | 0.020618 |
| DNAJC8P1 | 14 | 0.0001646 | 0.020618 |
| UBE2V2 | 8 | 0.0001646 | 0.020618 |
| SLC44A1 | 9 | 0.0001646 | 0.020618 |
| MDGA1 | 6 | 0.0001667 | 0.020797 |
| RGL1 | 1 | 0.0001686 | 0.020808 |
| ZNF688 | 16 | 0.0001712 | 0.020808 |
| TCF25 | 16 | 0.0001725 | 0.020808 |
| SETDB1 | 1 | 0.0001732 | 0.020808 |
| SYCP3 | 12 | 0.0001741 | 0.020808 |
| IFRD1 | 7 | 0.0001741 | 0.020808 |
| EWSR1 | 22 | 0.0001751 | 0.020808 |
| USP30 | 12 | 0.0001759 | 0.020808 |
| MLLT3 | 9 | 0.0001759 | 0.020808 |
| DLAT | 11 | 0.0001762 | 0.020808 |
| ZBTB11 | 3 | 0.0001762 | 0.020808 |
| FBXO38 | 5 | 0.0001762 | 0.020808 |
| BIN3 | 8 | 0.0001771 | 0.020808 |
| MAEL | 1 | 0.0001771 | 0.020808 |
| FHIP2A | 10 | 0.0001771 | 0.020808 |
| MBTPS1 | 16 | 0.0001775 | 0.020808 |
| CDH17 | 8 | 0.0001775 | 0.020808 |
| TSPOAP1 | 17 | 0.0001842 | 0.021309 |
| TRBV12-3 | 7 | 0.0001842 | 0.021309 |
| CFAP91 | 3 | 0.0001857 | 0.021309 |

| | | | |
|---|---|---|---|
| SLC35A1 | 6 | 0.0001857 | 0.021309 |
| XPR1 | 1 | 0.000187 | 0.021309 |
| DPF3 | 14 | 0.000187 | 0.021309 |
| PCDHA7 | 5 | 0.000187 | 0.021309 |
| RPS12 | 6 | 0.000187 | 0.021309 |
| MYO3B | 2 | 0.0001887 | 0.021382 |
| GTF2IRD2 | 7 | 0.000189 | 0.021382 |
| SPATS2L | 2 | 0.0001913 | 0.021495 |
| CFAP298 | 21 | 0.0001913 | 0.021495 |
| ICAM5 | 19 | 0.0001954 | 0.021720 |
| PDCD6IP | 3 | 0.0001957 | 0.021720 |
| KCNAB1 | 3 | 0.0001971 | 0.021720 |
| VRK2 | 2 | 0.0001972 | 0.021720 |
| TASOR | 3 | 0.0001972 | 0.021720 |
| NBN | 8 | 0.0001972 | 0.021720 |
| PPFIA1 | 11 | 0.0001986 | 0.021795 |
| THOC1 | 18 | 0.0002018 | 0.022080 |
| TUBAP2 | 11 | 0.0002035 | 0.022111 |
| AC138207.8 | 17 | 0.0002035 | 0.022111 |
| ABCB10 | 1 | 0.0002047 | 0.022168 |
| CDK8 | 13 | 0.0002077 | 0.022346 |
| SLC12A6 | 15 | 0.0002077 | 0.022346 |
| CCDC62 | 12 | 0.0002093 | 0.022365 |
| OSCP1 | 1 | 0.0002107 | 0.022365 |
| MEIS1 | 2 | 0.0002107 | 0.022365 |
| BNIP1 | 5 | 0.0002113 | 0.022365 |
| NOS3 | 7 | 0.0002113 | 0.022365 |
| UBR2 | 6 | 0.0002137 | 0.022503 |
| RANBP10 | 16 | 0.0002145 | 0.022503 |
| BAG4 | 8 | 0.0002154 | 0.022503 |
| NDUFAF5 | 20 | 0.0002167 | 0.022503 |
| MGP | 12 | 0.0002171 | 0.022503 |
| GPR137C | 14 | 0.0002174 | 0.022503 |
| NME1-NME2 | 17 | 0.0002174 | 0.022503 |
| RELL1 | 4 | 0.0002181 | 0.022503 |
| ATPAF1 | 1 | 0.0002218 | 0.022747 |
| MLF1 | 3 | 0.0002218 | 0.022747 |
| LYPD6 | 2 | 0.0002308 | 0.023600 |
| CLEC9A | 12 | 0.000233 | 0.023672 |
| PCOLCE2 | 3 | 0.000233 | 0.023672 |
| OR7G3 | 19 | 0.0002342 | 0.023717 |
| COMMD10 | 5 | 0.0002353 | 0.023756 |
| PAICS | 4 | 0.0002387 | 0.023759 |
| MKNK1 | 1 | 0.0002395 | 0.023759 |
| TEFM | 17 | 0.0002396 | 0.023759 |
| SLC24A5 | 15 | 0.0002414 | 0.023759 |
| SELENOM | 22 | 0.0002414 | 0.023759 |
| COMMD3 | 10 | 0.0002424 | 0.023759 |

| | | | |
|---|---|---|---|
| ACTR2 | 2 | 0.0002424 | 0.023759 |
| EMC3 | 3 | 0.0002424 | 0.023759 |
| YWHAZ | 8 | 0.0002424 | 0.023759 |
| FAF1 | 1 | 0.0002429 | 0.023759 |
| VWA5A | 11 | 0.0002453 | 0.023759 |
| USP28 | 11 | 0.0002485 | 0.023759 |
| ARHGAP12 | 10 | 0.0002486 | 0.023759 |
| OR5M4P | 11 | 0.0002486 | 0.023759 |
| COL12A1 | 6 | 0.0002491 | 0.023759 |
| NAA50 | 3 | 0.0002497 | 0.023759 |
| ZIC1 | 3 | 0.0002497 | 0.023759 |
| KIF23 | 15 | 0.0002514 | 0.023759 |
| ERLEC1 | 2 | 0.0002514 | 0.023759 |
| COL21A1 | 6 | 0.0002515 | 0.023759 |
| ADD3 | 10 | 0.0002525 | 0.023759 |
| CIB3 | 19 | 0.0002525 | 0.023759 |
| USP45 | 6 | 0.0002525 | 0.023759 |
| TMEM263 | 12 | 0.0002567 | 0.023759 |
| THUMPD1 | 16 | 0.0002567 | 0.023759 |
| OXCT1 | 5 | 0.000258 | 0.023759 |
| ECPAS | 9 | 0.000258 | 0.023759 |
| PIWIL1 | 12 | 0.0002586 | 0.023759 |
| SPATA18 | 4 | 0.0002586 | 0.023759 |
| RBM34 | 1 | 0.0002589 | 0.023759 |
| RBKS | 2 | 0.0002599 | 0.023759 |
| C2orf42 | 2 | 0.0002599 | 0.023759 |
| FER1L5 | 2 | 0.0002599 | 0.023759 |
| PLSCR5 | 3 | 0.0002599 | 0.023759 |
| ATP2B4 | 1 | 0.0002661 | 0.024255 |
| SUMO1 | 2 | 0.0002682 | 0.024385 |
| SPAG9 | 17 | 0.0002714 | 0.024466 |
| INPP5D | 2 | 0.0002714 | 0.024466 |
| SCAF8 | 6 | 0.0002714 | 0.024466 |
| TMCO1 | 1 | 0.0002747 | 0.024500 |
| DDX50 | 10 | 0.0002747 | 0.024500 |
| RABEP2 | 16 | 0.0002747 | 0.024500 |
| UBXN4 | 2 | 0.0002747 | 0.024500 |
| UTP20 | 12 | 0.0002755 | 0.024506 |
| ACP3 | 3 | 0.0002801 | 0.024778 |
| RAB3C | 5 | 0.0002801 | 0.024778 |
| OXR1 | 8 | 0.0002822 | 0.024815 |
| H2AJ | 12 | 0.0002855 | 0.024815 |
| NF1P1 | 15 | 0.0002855 | 0.024815 |
| LINC02248 | 15 | 0.0002855 | 0.024815 |
| ELK3 | 12 | 0.000286 | 0.024815 |
| GLT8D2 | 12 | 0.000286 | 0.024815 |
| ZNF433 | 19 | 0.000286 | 0.024815 |
| NDC1 | 1 | 0.0002903 | 0.024815 |

| | | | |
|---|---|---|---|
| ATRNL1 | 10 | 0.0002903 | 0.024815 |
| PRPF18 | 10 | 0.0002908 | 0.024815 |
| PSME3 | 17 | 0.0002908 | 0.024815 |
| TRIM13 | 13 | 0.0002937 | 0.024815 |
| EOMES | 3 | 0.0002937 | 0.024815 |
| CHRNB4 | 15 | 0.0002937 | 0.024815 |
| ZNF236 | 18 | 0.0002946 | 0.024815 |
| CGB1 | 19 | 0.0002955 | 0.024815 |
| SAE1 | 19 | 0.0002966 | 0.024815 |
| HACD1 | 10 | 0.0002966 | 0.024815 |
| COPZ1 | 12 | 0.0002966 | 0.024815 |
| WDR7 | 18 | 0.0002989 | 0.024815 |
| SLC27A6 | 5 | 0.0002989 | 0.024815 |
| HARS2 | 5 | 0.0002989 | 0.024815 |
| RPF2 | 6 | 0.0002989 | 0.024815 |
| ZNF280D | 15 | 0.0002994 | 0.024815 |
| PAPOLG | 2 | 0.0002994 | 0.024815 |
| PLXNC1 | 12 | 0.0003033 | 0.024872 |
| C12orf4 | 12 | 0.0003039 | 0.024872 |
| RAD51D | 17 | 0.0003039 | 0.024872 |
| SATB2 | 2 | 0.0003039 | 0.024872 |
| KPNA5 | 6 | 0.0003039 | 0.024872 |
| INTS3 | 1 | 0.000316 | 0.025795 |
| PDGFRL | 8 | 0.0003227 | 0.025992 |
| ARPC5 | 1 | 0.000324 | 0.025992 |
| MACROD2 | 20 | 0.0003254 | 0.025992 |
| SLC25A5P3 | 7 | 0.0003254 | 0.025992 |
| MARS1 | 12 | 0.0003256 | 0.025992 |
| ANO4 | 12 | 0.0003268 | 0.025992 |
| BNIPL | 1 | 0.000327 | 0.025992 |
| IFITM2 | 11 | 0.000327 | 0.025992 |
| RNF215 | 22 | 0.000327 | 0.025992 |
| GDAP1 | 8 | 0.000327 | 0.025992 |
| FAM210A | 18 | 0.0003271 | 0.025992 |
| ORMDL2 | 12 | 0.000331 | 0.026173 |
| COPS6 | 7 | 0.000331 | 0.026173 |
| MYPN | 10 | 0.0003378 | 0.026648 |
| COX15 | 10 | 0.000341 | 0.026773 |
| THAP2 | 12 | 0.000341 | 0.026773 |
| AZIN1 | 8 | 0.0003421 | 0.026792 |
| GGA2 | 16 | 0.0003472 | 0.027114 |
| ELMOD1 | 11 | 0.0003487 | 0.027114 |
| CLMN | 14 | 0.0003487 | 0.027114 |
| TRPM1 | 15 | 0.0003497 | 0.027114 |
| OPRD1 | 1 | 0.0003533 | 0.027114 |
| RANBP3 | 19 | 0.0003534 | 0.027114 |
| LRP12 | 8 | 0.0003534 | 0.027114 |
| HEATR3 | 16 | 0.0003545 | 0.027114 |

| | | | |
|---|---|---|---|
| LUC7L2 | 7 | 0.0003545 | 0.027114 |
| GLB1 | 3 | 0.000356 | 0.027114 |
| FKBP3 | 14 | 0.0003561 | 0.027114 |
| ELMOD2 | 4 | 0.0003561 | 0.027114 |
| BBS9 | 7 | 0.00036 | 0.027346 |
| AC009094.1 | 16 | 0.0003718 | 0.028181 |
| SMARCD3 | 7 | 0.0003801 | 0.028561 |
| PSMA5 | 1 | 0.0003813 | 0.028561 |
| ZNF562 | 19 | 0.0003813 | 0.028561 |
| TOP2B | 3 | 0.0003813 | 0.028561 |
| CBR4 | 4 | 0.0003813 | 0.028561 |
| RPL34P27 | 13 | 0.0003829 | 0.028561 |
| FBXO48 | 2 | 0.0003829 | 0.028561 |
| ERCC6L2 | 9 | 0.0003898 | 0.028980 |
| PLEKHM2 | 1 | 0.0003937 | 0.028980 |
| ELP4 | 11 | 0.0003937 | 0.028980 |
| TMEM167A | 5 | 0.0003937 | 0.028980 |
| PDE1C | 7 | 0.000395 | 0.028980 |
| MORN3 | 12 | 0.0003953 | 0.028980 |
| UMPS | 3 | 0.0003953 | 0.028980 |
| AKAP6 | 14 | 0.0003965 | 0.028980 |
| PRKCB | 16 | 0.0003965 | 0.028980 |
| PRELID3B | 20 | 0.0003987 | 0.029053 |
| CUX1 | 7 | 0.0003993 | 0.029053 |
| NCKAP1 | 2 | 0.0004015 | 0.029154 |
| ITGB2 | 21 | 0.0004035 | 0.029165 |
| ADGRG6 | 6 | 0.0004035 | 0.029165 |
| DAZL | 3 | 0.0004119 | 0.029708 |
| BMP5 | 6 | 0.0004138 | 0.029747 |
| GPATCH3 | 1 | 0.0004197 | 0.029747 |
| HNRNPUL2 | 11 | 0.0004197 | 0.029747 |
| IREB2 | 15 | 0.0004197 | 0.029747 |
| TRPS1 | 8 | 0.0004197 | 0.029747 |
| PHTF1 | 1 | 0.0004203 | 0.029747 |
| SKAP1 | 17 | 0.0004203 | 0.029747 |
| TPMT | 6 | 0.0004203 | 0.029747 |
| ATF7IP | 12 | 0.0004218 | 0.029747 |
| PIWIL4 | 11 | 0.0004224 | 0.029747 |
| TATDN1 | 8 | 0.0004224 | 0.029747 |
| HNRNPH1 | 5 | 0.0004246 | 0.029759 |
| AC024587.1 | 5 | 0.0004246 | 0.029759 |
| ANKFY1 | 17 | 0.0004253 | 0.029759 |
| GSTP1 | 11 | 0.0004404 | 0.030551 |
| LRIT3 | 4 | 0.0004404 | 0.030551 |
| POU2F1 | 1 | 0.0004438 | 0.030551 |
| DUSP10 | 1 | 0.0004438 | 0.030551 |
| METTL7A | 12 | 0.0004438 | 0.030551 |
| TRAPPC6B | 14 | 0.0004438 | 0.030551 |

| | | | |
|---|---|---|---|
| SPATC1 | 8 | 0.0004438 | 0.030551 |
| USP15 | 12 | 0.0004459 | 0.030551 |
| RNASEH2B | 13 | 0.0004459 | 0.030551 |
| HDGFL3 | 15 | 0.0004459 | 0.030551 |
| NCAN | 19 | 0.0004469 | 0.030551 |
| KNOP1 | 16 | 0.0004523 | 0.030854 |
| NIN | 14 | 0.0004532 | 0.030854 |
| SNORD36 | 13 | 0.0004559 | 0.030882 |
| CKLF | 16 | 0.0004559 | 0.030882 |
| MFSD14A | 1 | 0.0004573 | 0.030882 |
| SLC19A2 | 1 | 0.0004573 | 0.030882 |
| GORASP1 | 3 | 0.0004646 | 0.031109 |
| PHYHIP | 8 | 0.0004646 | 0.031109 |
| ACAD9 | 3 | 0.0004656 | 0.031109 |
| RNF185 | 22 | 0.0004664 | 0.031109 |
| LRAT | 4 | 0.0004664 | 0.031109 |
| PAQR8 | 6 | 0.0004664 | 0.031109 |
| ABI1 | 10 | 0.000469 | 0.031157 |
| NDUFAF6 | 8 | 0.000469 | 0.031157 |
| MACO1 | 1 | 0.0004857 | 0.031677 |
| NBPF19 | 1 | 0.0004857 | 0.031677 |
| EDARADD | 1 | 0.0004857 | 0.031677 |
| BFAR | 16 | 0.0004857 | 0.031677 |
| ESF1 | 20 | 0.0004857 | 0.031677 |
| WBP11 | 12 | 0.0004865 | 0.031677 |
| MAN2C1 | 15 | 0.0004865 | 0.031677 |
| KPNA2P3 | 17 | 0.0004865 | 0.031677 |
| RGL3 | 19 | 0.0004865 | 0.031677 |
| ANKRD7 | 7 | 0.0004865 | 0.031677 |
| CLCN6 | 1 | 0.0004922 | 0.031860 |
| SUFU | 10 | 0.0004922 | 0.031860 |
| CFAP221 | 2 | 0.0004922 | 0.031860 |
| DDX52 | 17 | 0.000496 | 0.031981 |
| NEPRO | 3 | 0.000496 | 0.031981 |
| GPAT3 | 4 | 0.0004989 | 0.032004 |
| MTBP | 8 | 0.0004989 | 0.032004 |
| PYROXD1 | 12 | 0.0005003 | 0.032004 |
| AGFG1 | 2 | 0.0005003 | 0.032004 |
| GNPTG | 16 | 0.0005066 | 0.032074 |
| HNRNPA1P52 | 19 | 0.0005066 | 0.032074 |
| EXTL2P1 | 2 | 0.0005066 | 0.032074 |
| DDX24 | 14 | 0.0005075 | 0.032074 |
| MGRN1 | 16 | 0.0005075 | 0.032074 |
| LSS | 21 | 0.0005075 | 0.032074 |
| ANGPTL4 | 19 | 0.0005157 | 0.032074 |
| PRKAB2 | 1 | 0.0005166 | 0.032074 |
| MR1 | 1 | 0.0005166 | 0.032074 |
| RAB4A | 1 | 0.0005166 | 0.032074 |

| | | | |
|---|---|---|---|
| SRRD | 22 | 0.0005166 | 0.032074 |
| GTF2H2B | 5 | 0.0005166 | 0.032074 |
| SEPHS1P6 | 2 | 0.0005176 | 0.032074 |
| GLTP | 12 | 0.0005186 | 0.032074 |
| XPNPEP3 | 22 | 0.0005186 | 0.032074 |
| SLC35G2 | 3 | 0.0005186 | 0.032074 |
| HEYL | 1 | 0.00052 | 0.032074 |
| DHH | 12 | 0.00052 | 0.032074 |
| IGF1 | 12 | 0.00052 | 0.032074 |
| PRMT8 | 12 | 0.0005223 | 0.032089 |
| GPD2 | 2 | 0.0005231 | 0.032089 |
| CORIN | 4 | 0.0005231 | 0.032089 |
| UTP15 | 5 | 0.0005283 | 0.032203 |
| GOLGA7 | 8 | 0.0005283 | 0.032203 |
| ANKRD17 | 4 | 0.0005286 | 0.032203 |
| GOLGA3 | 12 | 0.0005319 | 0.032203 |
| COG4 | 16 | 0.0005338 | 0.032203 |
| ALS2CL | 3 | 0.0005338 | 0.032203 |
| ZBBX | 3 | 0.0005338 | 0.032203 |
| CASD1 | 7 | 0.0005338 | 0.032203 |
| ATP6V1C1 | 8 | 0.0005338 | 0.032203 |
| HIVEP2 | 6 | 0.0005432 | 0.032707 |
| OTULIN | 5 | 0.000546 | 0.032820 |
| CACUL1 | 10 | 0.0005508 | 0.032900 |
| ARRDC4 | 15 | 0.0005508 | 0.032900 |
| DNTTIP1 | 20 | 0.0005508 | 0.032900 |
| PDZD2 | 5 | 0.0005514 | 0.032900 |
| TRIM24 | 7 | 0.0005665 | 0.033466 |
| DHX40 | 17 | 0.0005673 | 0.033466 |
| PLSCR2 | 3 | 0.0005673 | 0.033466 |
| IL6ST | 5 | 0.0005673 | 0.033466 |
| CANX | 5 | 0.0005718 | 0.033466 |
| INTS10 | 8 | 0.0005718 | 0.033466 |
| UNC5C | 4 | 0.0005721 | 0.033466 |
| CBX3 | 7 | 0.0005721 | 0.033466 |
| E2F2 | 1 | 0.0005721 | 0.033466 |
| TRAF6 | 11 | 0.0005721 | 0.033466 |
| KRT39 | 17 | 0.0005721 | 0.033466 |
| INHBA | 7 | 0.000577 | 0.033694 |
| MED13L | 12 | 0.0005788 | 0.033738 |
| COG1 | 17 | 0.0005843 | 0.033931 |
| ANLN | 7 | 0.0005843 | 0.033931 |
| AXIN1 | 16 | 0.0005852 | 0.033931 |
| ARNT | 1 | 0.000591 | 0.034051 |
| PICALM | 11 | 0.000591 | 0.034051 |
| PPP1R21 | 2 | 0.0005915 | 0.034051 |
| BEND3 | 6 | 0.0005915 | 0.034051 |
| OLAH | 10 | 0.0005935 | 0.034051 |

| | | | |
|---|---|---|---|
| AC139426.1 | 15 | 0.0005935 | 0.034051 |
| TENM3 | 4 | 0.0005957 | 0.034088 |
| TRIM21 | 11 | 0.0005983 | 0.034088 |
| KRT72 | 12 | 0.0005983 | 0.034088 |
| MTRF1 | 13 | 0.0005983 | 0.034088 |
| ANXA11 | 10 | 0.0006084 | 0.034209 |
| NSD3 | 8 | 0.0006084 | 0.034209 |
| LUZP4P1 | 10 | 0.0006092 | 0.034209 |
| RNLS | 10 | 0.0006109 | 0.034209 |
| C11orf65 | 11 | 0.0006109 | 0.034209 |
| KIAA0753 | 17 | 0.0006109 | 0.034209 |
| TSPAN16 | 19 | 0.0006109 | 0.034209 |
| GPI | 19 | 0.0006109 | 0.034209 |
| HOXD8 | 2 | 0.0006109 | 0.034209 |
| EOGT | 3 | 0.0006109 | 0.034209 |
| HDHD2 | 18 | 0.0006169 | 0.034372 |
| CCDC68 | 18 | 0.0006169 | 0.034372 |
| GTF3C6 | 6 | 0.0006169 | 0.034372 |
| PDIA5 | 3 | 0.00062 | 0.034487 |
| PRRC2A | 6 | 0.0006295 | 0.034954 |
| KBTBD8 | 3 | 0.0006308 | 0.034968 |
| DZIP3 | 3 | 0.0006344 | 0.035108 |
| MATN2 | 8 | 0.0006364 | 0.035158 |
| TLCD4 | 1 | 0.0006458 | 0.035440 |
| PRPF40A | 2 | 0.0006458 | 0.035440 |
| POGLUT1 | 3 | 0.0006458 | 0.035440 |
| ABRAXAS1 | 4 | 0.0006458 | 0.035440 |
| PAPSS2 | 10 | 0.0006504 | 0.035440 |
| SMC3 | 10 | 0.0006504 | 0.035440 |
| SYNRG | 17 | 0.0006504 | 0.035440 |
| TEP1 | 14 | 0.0006551 | 0.035440 |
| EFCAB6 | 22 | 0.0006551 | 0.035440 |
| IFNWP15 | 9 | 0.00066 | 0.035440 |
| LMBRD2 | 5 | 0.0006603 | 0.035440 |
| SEMA3D | 7 | 0.0006603 | 0.035440 |
| ZNF350 | 19 | 0.0006604 | 0.035440 |
| PSMC2 | 7 | 0.0006604 | 0.035440 |
| HEY1 | 8 | 0.0006604 | 0.035440 |
| ANKRD46 | 8 | 0.0006604 | 0.035440 |
| CRTAC1 | 10 | 0.000672 | 0.035440 |
| CEP57 | 11 | 0.000672 | 0.035440 |
| MTIF2 | 2 | 0.000672 | 0.035440 |
| SEPTIN2 | 2 | 0.000672 | 0.035440 |
| MTMR12 | 5 | 0.000672 | 0.035440 |
| AKR1B10 | 7 | 0.000672 | 0.035440 |
| JAM3 | 11 | 0.0006778 | 0.035440 |
| FBH1 | 10 | 0.0006792 | 0.035440 |
| ADK | 10 | 0.0006792 | 0.035440 |

| | | | |
|---|---|---|---|
| SNAPC1 | 14 | 0.0006792 | 0.035440 |
| TTK | 6 | 0.0006792 | 0.035440 |
| DICER1 | 14 | 0.0006799 | 0.035440 |
| TCF4 | 18 | 0.0006799 | 0.035440 |
| WASHC2A | 10 | 0.0006868 | 0.035440 |
| TRPV2 | 17 | 0.0006887 | 0.035440 |
| GNL2 | 1 | 0.0006893 | 0.035440 |
| KIN | 10 | 0.0006893 | 0.035440 |
| RPAIN | 17 | 0.0006893 | 0.035440 |
| SIPA1L3 | 19 | 0.0006913 | 0.035440 |
| ATRN | 20 | 0.0006913 | 0.035440 |
| DNAJC8 | 1 | 0.0006922 | 0.035440 |
| GP1BA | 17 | 0.0006922 | 0.035440 |
| ERAL1 | 17 | 0.0006922 | 0.035440 |
| ASB14 | 3 | 0.0006922 | 0.035440 |
| TDRD5 | 1 | 0.0006923 | 0.035440 |
| PUM3 | 9 | 0.0006923 | 0.035440 |
| FAAP24 | 19 | 0.000693 | 0.035440 |
| ASTE1 | 3 | 0.000693 | 0.035440 |
| AL158825.1 | 9 | 0.000693 | 0.035440 |
| RCHY1 | 4 | 0.0006951 | 0.035440 |
| ASZ1 | 7 | 0.0006951 | 0.035440 |
| KCNAB2 | 1 | 0.0006981 | 0.035440 |
| SLF2 | 10 | 0.0006981 | 0.035440 |
| TMEM183A | 1 | 0.0006989 | 0.035440 |
| CIR1 | 2 | 0.0006989 | 0.035440 |
| WNT8A | 5 | 0.0006989 | 0.035440 |
| RAD21 | 8 | 0.0006989 | 0.035440 |
| KALRN | 3 | 0.0006998 | 0.035440 |
| CD93 | 20 | 0.0007012 | 0.035458 |
| ARHGAP30 | 1 | 0.0007133 | 0.035901 |
| CCPG1 | 15 | 0.0007133 | 0.035901 |
| SP8 | 7 | 0.0007133 | 0.035901 |
| LRRCC1 | 8 | 0.0007252 | 0.036445 |
| WDR17 | 4 | 0.0007555 | 0.037465 |
| KMT2E | 7 | 0.0007555 | 0.037465 |
| C12orf29 | 12 | 0.0007556 | 0.037465 |
| XCL2 | 1 | 0.0007561 | 0.037465 |
| FTO | 16 | 0.0007561 | 0.037465 |
| DYNC1LI1 | 3 | 0.0007561 | 0.037465 |
| CDC14B | 9 | 0.0007561 | 0.037465 |
| DIAPH3 | 13 | 0.000757 | 0.037465 |
| ATP2A3 | 17 | 0.000757 | 0.037465 |
| PTCD3 | 2 | 0.000757 | 0.037465 |
| LINC01977 | 17 | 0.000758 | 0.037465 |
| RASGRF2 | 5 | 0.0007657 | 0.037714 |
| SLC16A4 | 1 | 0.0007677 | 0.037714 |
| ITPK1 | 14 | 0.0007677 | 0.037714 |

| | | | |
|---|---|---|---|
| KCTD16 | 5 | 0.0007677 | 0.037714 |
| PHAX | 5 | 0.0007726 | 0.037898 |
| BANK1 | 4 | 0.0007769 | 0.037939 |
| VPS41 | 7 | 0.0007769 | 0.037939 |
| OR52N5 | 11 | 0.000781 | 0.037939 |
| ZNF235 | 19 | 0.000781 | 0.037939 |
| TNFRSF8 | 1 | 0.0007839 | 0.037939 |
| PPFIBP2 | 11 | 0.0007839 | 0.037939 |
| CLTC | 17 | 0.0007839 | 0.037939 |
| RACGAP1 | 12 | 0.0007884 | 0.037939 |
| OGFOD1 | 16 | 0.0007884 | 0.037939 |
| PRKCSH | 19 | 0.0007884 | 0.037939 |
| TCAIM | 3 | 0.0007884 | 0.037939 |
| NEK4 | 3 | 0.0007884 | 0.037939 |
| MBD4 | 3 | 0.0007884 | 0.037939 |
| ANXA3 | 4 | 0.0008014 | 0.038175 |
| HOMER1 | 5 | 0.0008014 | 0.038175 |
| C1orf112 | 1 | 0.0008048 | 0.038175 |
| DACH1 | 13 | 0.0008048 | 0.038175 |
| NUP35 | 2 | 0.0008048 | 0.038175 |
| RFPL3 | 22 | 0.0008048 | 0.038175 |
| STK32A | 5 | 0.0008048 | 0.038175 |
| ESRP1 | 8 | 0.0008048 | 0.038175 |
| PKD1L2 | 16 | 0.0008051 | 0.038175 |
| ZNF587 | 19 | 0.0008058 | 0.038175 |
| LIPJ | 10 | 0.0008105 | 0.038175 |
| CARD14 | 17 | 0.0008105 | 0.038175 |
| IQGAP2 | 5 | 0.0008127 | 0.038175 |
| CAPZA1 | 1 | 0.0008188 | 0.038175 |
| CTTN | 11 | 0.0008188 | 0.038175 |
| CACNA2D2 | 3 | 0.0008188 | 0.038175 |
| AHCYL2 | 7 | 0.0008188 | 0.038175 |
| CHST4 | 16 | 0.0008198 | 0.038175 |
| CCDC43 | 17 | 0.0008198 | 0.038175 |
| HMGB1P20 | 6 | 0.0008198 | 0.038175 |
| VPS13D | 1 | 0.0008214 | 0.038175 |
| CLSPN | 1 | 0.0008224 | 0.038175 |
| TRIM25 | 17 | 0.0008224 | 0.038175 |
| PPM1B | 2 | 0.0008224 | 0.038175 |
| MEF2C | 5 | 0.0008224 | 0.038175 |
| SEC14L1 | 17 | 0.000825 | 0.038185 |
| NMD3 | 3 | 0.000825 | 0.038185 |
| UTP11 | 1 | 0.0008316 | 0.038201 |
| ANKRD24 | 19 | 0.0008316 | 0.038201 |
| ST3GAL5 | 2 | 0.0008316 | 0.038201 |
| SP3 | 2 | 0.0008316 | 0.038201 |
| CLK4 | 5 | 0.0008316 | 0.038201 |
| PATJ | 1 | 0.0008323 | 0.038201 |

| | | | |
|---|---|---|---|
| CIC | 19 | 0.0008344 | 0.038245 |
| SEC24B | 4 | 0.0008626 | 0.039427 |
| SLC44A4 | 6 | 0.0008626 | 0.039427 |
| NKAIN3 | 8 | 0.0008712 | 0.039763 |
| CFAP74 | 1 | 0.000877 | 0.039837 |
| PILRB | 7 | 0.0008787 | 0.039837 |
| MCHR1 | 22 | 0.0008836 | 0.039837 |
| ENPP4 | 6 | 0.0008836 | 0.039837 |
| IL23R | 1 | 0.0008837 | 0.039837 |
| CRYZ | 1 | 0.0008837 | 0.039837 |
| HSPA14 | 10 | 0.0008837 | 0.039837 |
| PAN2 | 12 | 0.0008837 | 0.039837 |
| LONRF2 | 2 | 0.0008837 | 0.039837 |
| METTL23 | 17 | 0.000891 | 0.040055 |
| RNF19A | 8 | 0.000891 | 0.040055 |
| ARHGAP18 | 6 | 0.0008931 | 0.040096 |
| L2HGDH | 14 | 0.0008987 | 0.040180 |
| CDYL | 6 | 0.0008987 | 0.040180 |
| MICU3 | 8 | 0.0008987 | 0.040180 |
| COPG2 | 7 | 0.0009045 | 0.040384 |
| AGAP6 | 10 | 0.0009237 | 0.040704 |
| POLA2 | 11 | 0.0009237 | 0.040704 |
| IDH1 | 2 | 0.0009237 | 0.040704 |
| ADD1 | 4 | 0.0009237 | 0.040704 |
| TFR2 | 7 | 0.0009237 | 0.040704 |
| DBH | 9 | 0.0009237 | 0.040704 |
| STK11 | 19 | 0.0009244 | 0.040704 |
| AURKA | 20 | 0.0009244 | 0.040704 |
| GABRR3 | 3 | 0.0009244 | 0.040704 |
| RBM47 | 4 | 0.0009253 | 0.040704 |
| CEP57L1 | 6 | 0.0009253 | 0.040704 |
| HSD17B4 | 5 | 0.0009288 | 0.040804 |
| CHD5 | 1 | 0.0009338 | 0.040867 |
| HDAC4 | 2 | 0.0009338 | 0.040867 |
| DPYD | 1 | 0.0009435 | 0.040867 |
| CEP164 | 11 | 0.0009435 | 0.040867 |
| SLC25A41 | 19 | 0.0009451 | 0.040867 |
| NAT2 | 8 | 0.0009451 | 0.040867 |
| CHMP4C | 8 | 0.0009451 | 0.040867 |
| AL137792.1 | 1 | 0.0009475 | 0.040867 |
| SLC25A3 | 12 | 0.0009514 | 0.040867 |
| GMCL1 | 2 | 0.0009514 | 0.040867 |
| QRSL1 | 6 | 0.0009514 | 0.040867 |
| PNPLA8 | 7 | 0.0009514 | 0.040867 |
| TC2N | 14 | 0.0009518 | 0.040867 |
| ATP13A3 | 3 | 0.0009526 | 0.040867 |
| ST3GAL3 | 1 | 0.0009666 | 0.040867 |
| USP54 | 10 | 0.0009666 | 0.040867 |

| | | | |
|---|---|---|---|
| MYO1A | 12 | 0.0009666 | 0.040867 |
| LYN | 8 | 0.0009666 | 0.040867 |
| DAP3 | 1 | 0.0009673 | 0.040867 |
| LMNTD1 | 12 | 0.0009673 | 0.040867 |
| HIF1A | 14 | 0.0009673 | 0.040867 |
| STPG2 | 4 | 0.0009673 | 0.040867 |
| TCP1 | 6 | 0.0009673 | 0.040867 |
| KCNB2 | 8 | 0.0009673 | 0.040867 |
| DENND10 | 10 | 0.0009681 | 0.040867 |
| DYRK4 | 12 | 0.0009681 | 0.040867 |
| METTL9 | 16 | 0.0009681 | 0.040867 |
| FHOD1 | 16 | 0.0009681 | 0.040867 |
| TXNDC9 | 2 | 0.0009681 | 0.040867 |
| TMIGD2 | 19 | 0.0009721 | 0.040867 |
| TBC1D23 | 3 | 0.0009721 | 0.040867 |
| HSF2 | 6 | 0.0009721 | 0.040867 |
| IFT74 | 9 | 0.0009721 | 0.040867 |
| ANKRD20A21P | 20 | 0.0009726 | 0.040867 |
| NPM1P42 | 15 | 0.0009769 | 0.040906 |
| TEKT5 | 16 | 0.0009773 | 0.040906 |
| PIK3CG | 7 | 0.0009773 | 0.040906 |
| CHST1 | 11 | 0.0009973 | 0.041672 |
| KIF20B | 10 | 0.0010007 | 0.041672 |
| ALDH1L2 | 12 | 0.0010007 | 0.041672 |
| MET | 7 | 0.0010007 | 0.041672 |
| POLG2 | 17 | 0.0010019 | 0.041672 |
| WDR90 | 16 | 0.0010111 | 0.041935 |
| MRC1 | 10 | 0.0010122 | 0.041935 |
| ROS1 | 6 | 0.0010133 | 0.041935 |
| RPS6KA2 | 6 | 0.0010133 | 0.041935 |
| ZYG11B | 1 | 0.0010239 | 0.042166 |
| BICC1 | 10 | 0.0010239 | 0.042166 |
| SEMA3C | 7 | 0.0010239 | 0.042166 |
| ZCWPW1 | 7 | 0.0010314 | 0.042166 |
| ARHGEF19 | 1 | 0.0010326 | 0.042166 |
| TMEM45B | 11 | 0.0010326 | 0.042166 |
| RPL19 | 17 | 0.0010326 | 0.042166 |
| ZIM2 | 19 | 0.0010326 | 0.042166 |
| MRPL19 | 2 | 0.0010326 | 0.042166 |
| TAFA4 | 3 | 0.0010326 | 0.042166 |
| GPR27 | 3 | 0.0010367 | 0.042166 |
| PPP1R3B | 8 | 0.0010367 | 0.042166 |
| SPON1 | 11 | 0.0010382 | 0.042166 |
| TIA1 | 2 | 0.0010382 | 0.042166 |
| OSBPL6 | 2 | 0.0010382 | 0.042166 |
| TCP11L2 | 12 | 0.0010586 | 0.042785 |
| CBFB | 16 | 0.0010586 | 0.042785 |
| TPD52L1 | 6 | 0.0010586 | 0.042785 |

| | | | |
|---|---|---|---|
| RIDA | 8 | 0.0010586 | 0.042785 |
| CRISPLD1 | 8 | 0.0010647 | 0.042951 |
| NSL1 | 1 | 0.0010667 | 0.042951 |
| ALKBH8 | 11 | 0.0010667 | 0.042951 |
| ARHGAP28 | 18 | 0.0010801 | 0.043291 |
| PIGR | 1 | 0.0010804 | 0.043291 |
| G2E3 | 14 | 0.0010804 | 0.043291 |
| EIF4E3 | 3 | 0.0010804 | 0.043291 |
| PNPLA2 | 11 | 0.0010871 | 0.043384 |
| MRPL18 | 6 | 0.0010871 | 0.043384 |
| TSPAN33 | 7 | 0.0010871 | 0.043384 |
| OR7D1P | 19 | 0.0010907 | 0.043384 |
| RPS14P5 | 2 | 0.0010907 | 0.043384 |
| HOXD11 | 2 | 0.0010907 | 0.043384 |
| RNF10 | 12 | 0.0010984 | 0.043639 |
| MORC1 | 3 | 0.0011026 | 0.043754 |
| ZNF887P | 19 | 0.0011139 | 0.044148 |
| CWF19L1 | 10 | 0.0011236 | 0.044150 |
| DHRS7 | 14 | 0.0011236 | 0.044150 |
| MFN1 | 3 | 0.0011236 | 0.044150 |
| SDHA | 5 | 0.0011239 | 0.044150 |
| PYGO1 | 15 | 0.0011253 | 0.044150 |
| TMEM150C | 4 | 0.0011253 | 0.044150 |
| MACROH2A1 | 5 | 0.0011253 | 0.044150 |
| DCAF12 | 9 | 0.0011253 | 0.044150 |
| TAF12 | 1 | 0.0011294 | 0.044150 |
| KBTBD4 | 11 | 0.0011294 | 0.044150 |
| CTDSPL2 | 15 | 0.0011359 | 0.044150 |
| NTAN1 | 16 | 0.0011359 | 0.044150 |
| TSR1 | 17 | 0.0011359 | 0.044150 |
| CCT2 | 12 | 0.0011391 | 0.044150 |
| CCDC66 | 3 | 0.0011391 | 0.044150 |
| FRMD4B | 3 | 0.0011391 | 0.044150 |
| TACC3 | 4 | 0.0011391 | 0.044150 |
| EPHX1 | 1 | 0.0011435 | 0.044150 |
| ETV6 | 12 | 0.0011435 | 0.044150 |
| SLC44A2 | 19 | 0.0011435 | 0.044150 |
| RRM2 | 2 | 0.0011435 | 0.044150 |
| PRPF4 | 9 | 0.0011435 | 0.044150 |
| NADSYN1 | 11 | 0.0011585 | 0.044413 |
| USP31 | 16 | 0.0011585 | 0.044413 |
| CLASP2 | 3 | 0.0011585 | 0.044413 |
| AFAP1 | 4 | 0.0011585 | 0.044413 |
| LRRIQ1 | 12 | 0.0011598 | 0.044413 |
| PPIL2 | 22 | 0.0011598 | 0.044413 |
| DHX36 | 3 | 0.0011598 | 0.044413 |
| ATAD5 | 17 | 0.0011811 | 0.044724 |
| KIAA1217 | 10 | 0.0011941 | 0.044724 |

| | | | |
|---|---|---|---|
| SRPK2 | 7 | 0.0011941 | 0.044724 |
| ATP11B | 3 | 0.0011949 | 0.044724 |
| VCAM1 | 1 | 0.0011949 | 0.044724 |
| GTF2H2C | 5 | 0.0011949 | 0.044724 |
| MFSD4B | 6 | 0.0011949 | 0.044724 |
| MIOS | 7 | 0.0011949 | 0.044724 |
| GMEB1 | 1 | 0.001202 | 0.044724 |
| ELMO3 | 16 | 0.001202 | 0.044724 |
| TAF1C | 16 | 0.001202 | 0.044724 |
| SLC5A5 | 19 | 0.001202 | 0.044724 |
| PDK1 | 2 | 0.001202 | 0.044724 |
| IHO1 | 3 | 0.001202 | 0.044724 |
| TULP1 | 6 | 0.001202 | 0.044724 |
| PXT1 | 6 | 0.001202 | 0.044724 |
| RAB14 | 9 | 0.001202 | 0.044724 |
| ACRBP | 12 | 0.0012051 | 0.044724 |
| RPL21P11 | 14 | 0.0012051 | 0.044724 |
| ZFYVE26 | 14 | 0.0012053 | 0.044724 |
| ABCB5 | 7 | 0.0012053 | 0.044724 |
| DAPK1 | 9 | 0.0012053 | 0.044724 |
| CAPRIN1 | 11 | 0.0012102 | 0.044724 |
| BIRC2 | 11 | 0.0012102 | 0.044724 |
| VSX1 | 20 | 0.0012102 | 0.044724 |
| GMPSP1 | 4 | 0.0012102 | 0.044724 |
| CNPY3 | 6 | 0.0012102 | 0.044724 |
| NPC1 | 18 | 0.0012103 | 0.044724 |
| LAMB4 | 7 | 0.0012103 | 0.044724 |
| AL451050.1 | 1 | 0.0012124 | 0.044724 |
| FAM218BP | 4 | 0.0012124 | 0.044724 |
| CCDC178 | 18 | 0.0012129 | 0.044724 |
| ALCAM | 3 | 0.0012129 | 0.044724 |
| BAZ2B | 2 | 0.0012227 | 0.045035 |
| PDPN | 1 | 0.0012278 | 0.045069 |
| RAD51AP1 | 12 | 0.0012278 | 0.045069 |
| CCDC192 | 5 | 0.0012278 | 0.045069 |
| CHD9 | 16 | 0.0012344 | 0.045262 |
| ANKRD28 | 3 | 0.001259 | 0.045858 |
| PDCD4 | 10 | 0.0012618 | 0.045858 |
| SPG21 | 15 | 0.0012618 | 0.045858 |
| DHODH | 16 | 0.0012618 | 0.045858 |
| PLAUR | 19 | 0.0012618 | 0.045858 |
| D2HGDH | 2 | 0.0012618 | 0.045858 |
| FBXL2 | 3 | 0.0012618 | 0.045858 |
| CCT6A | 7 | 0.0012618 | 0.045858 |
| AFDN | 6 | 0.0012749 | 0.046167 |
| NIT1 | 1 | 0.001276 | 0.046167 |
| SFR1 | 10 | 0.001276 | 0.046167 |
| SOX5P1 | 8 | 0.001276 | 0.046167 |

| | | | |
|---|---|---|---|
| CRISPLD2 | 16 | 0.0012868 | 0.046356 |
| TCERG1 | 5 | 0.0012868 | 0.046356 |
| ABHD2 | 15 | 0.0012965 | 0.046356 |
| UBR3 | 2 | 0.0012965 | 0.046356 |
| L3MBTL2 | 22 | 0.0012965 | 0.046356 |
| MRPL2 | 6 | 0.0012965 | 0.046356 |
| LRRC40 | 1 | 0.0013049 | 0.046356 |
| EIF4E | 4 | 0.0013049 | 0.046356 |
| ZNF800 | 7 | 0.0013049 | 0.046356 |
| CPA6 | 8 | 0.0013049 | 0.046356 |
| LAMC1 | 1 | 0.001305 | 0.046356 |
| SMARCC2 | 12 | 0.001305 | 0.046356 |
| CEP55 | 10 | 0.0013062 | 0.046356 |
| NFKB2 | 10 | 0.0013062 | 0.046356 |
| KLF10 | 8 | 0.0013062 | 0.046356 |
| SLC24A3 | 20 | 0.0013066 | 0.046356 |
| SEC24D | 4 | 0.0013066 | 0.046356 |
| DPY19L4 | 8 | 0.0013066 | 0.046356 |
| COPA | 1 | 0.0013165 | 0.046604 |
| ANO5 | 11 | 0.0013165 | 0.046604 |
| LIPA | 10 | 0.001324 | 0.046686 |
| SIN3A | 15 | 0.001324 | 0.046686 |
| MRPL4 | 19 | 0.0013305 | 0.046686 |
| FBXO40 | 3 | 0.0013305 | 0.046686 |
| ACSS3 | 12 | 0.0013317 | 0.046686 |
| XKR6 | 8 | 0.0013317 | 0.046686 |
| KDM4A | 1 | 0.00134 | 0.046686 |
| CR2 | 1 | 0.00134 | 0.046686 |
| ACTR10 | 14 | 0.00134 | 0.046686 |
| PRICKLE2 | 3 | 0.00134 | 0.046686 |
| PLK4 | 4 | 0.00134 | 0.046686 |
| PUS7 | 7 | 0.00134 | 0.046686 |
| WASF2 | 1 | 0.001343 | 0.046686 |
| IPO4 | 14 | 0.001343 | 0.046686 |
| ALDH5A1 | 6 | 0.001343 | 0.046686 |
| IMPDH1 | 7 | 0.001343 | 0.046686 |
| SMC5 | 9 | 0.001343 | 0.046686 |
| EXOC6 | 10 | 0.0013736 | 0.047188 |
| TRIM9 | 14 | 0.0013736 | 0.047188 |
| WDR41 | 5 | 0.0013736 | 0.047188 |
| BAZ2A | 12 | 0.0013771 | 0.047188 |
| RUBCN | 3 | 0.0013771 | 0.047188 |
| EPHA7 | 6 | 0.0013771 | 0.047188 |
| PSMB5 | 14 | 0.0013808 | 0.047188 |
| MT1B | 16 | 0.0013808 | 0.047188 |
| SNX11 | 17 | 0.0013808 | 0.047188 |
| CEP97 | 3 | 0.0013808 | 0.047188 |
| PIM1 | 6 | 0.0013808 | 0.047188 |

| | | | |
|---|---|---|---|
| MTFR2 | 6 | 0.0013808 | 0.047188 |
| BRF2 | 8 | 0.0013808 | 0.047188 |
| PLPPR5 | 1 | 0.001381 | 0.047188 |
| ERGIC2 | 12 | 0.001381 | 0.047188 |
| SLC17A2 | 6 | 0.001381 | 0.047188 |
| BTAF1 | 10 | 0.0013818 | 0.047188 |
| LTBP1 | 2 | 0.0013963 | 0.047518 |
| DMBT1 | 10 | 0.0013968 | 0.047518 |
| IL17RD | 3 | 0.0013987 | 0.047518 |
| BMP3 | 4 | 0.0013987 | 0.047518 |
| UGT8 | 4 | 0.0013987 | 0.047518 |
| CREB1 | 2 | 0.001411 | 0.047836 |
| NF1P3 | 21 | 0.001411 | 0.047836 |
| PPP1R10 | 6 | 0.0014175 | 0.048003 |
| GRIK4 | 11 | 0.0014233 | 0.048003 |
| SPATA13 | 13 | 0.0014233 | 0.048003 |
| NOX3 | 6 | 0.0014233 | 0.048003 |
| ZNG1A[illegible] | 9 | 0.0014233 | 0.048003 |
| TRDMT1 | 10 | 0.0014339 | 0.048114 |
| KIAA0513 | 16 | 0.0014339 | 0.048114 |
| IGF2BP2 | 3 | 0.0014339 | 0.048114 |
| UFL1 | 6 | 0.0014339 | 0.048114 |
| ZC3HAV1 | 7 | 0.0014339 | 0.048114 |
| CLEC4A | 12 | 0.0014381 | 0.048206 |
| HEATR5B | 2 | 0.0014424 | 0.048303 |
| CEP120 | 5 | 0.0014448 | 0.048333 |
| VENTX | 10 | 0.0014623 | 0.048487 |
| FAM27E5 | 17 | 0.0014623 | 0.048487 |
| IQCJ | 3 | 0.0014623 | 0.048487 |
| RTP2 | 3 | 0.0014623 | 0.048487 |
| USP4 | 3 | 0.0014644 | 0.048487 |
| SNX9 | 6 | 0.0014644 | 0.048487 |
| UNC13B | 9 | 0.0014644 | 0.048487 |
| LHPP | 10 | 0.0014716 | 0.048487 |
| CNOT2 | 12 | 0.0014716 | 0.048487 |
| DHRS12 | 13 | 0.0014716 | 0.048487 |
| KXD1 | 19 | 0.0014716 | 0.048487 |
| RGL4 | 22 | 0.0014716 | 0.048487 |
| GSK3B | 3 | 0.0014716 | 0.048487 |
| KLC4 | 6 | 0.0014716 | 0.048487 |
| GCLC | 6 | 0.0014716 | 0.048487 |
| TNC | 9 | 0.0014785 | 0.048631 |
| PGM2L1 | 11 | 0.0014857 | 0.048631 |
| CHFR | 12 | 0.0014857 | 0.048631 |
| DSC1 | 18 | 0.0014857 | 0.048631 |
| BPIFB4 | 20 | 0.0014857 | 0.048631 |
| UBXN7 | 3 | 0.0014857 | 0.048631 |
| CARS2 | 13 | 0.0014863 | 0.048631 |

| | | | |
|---|---|---|---|
| METTL25 | 12 | 0.0015141 | 0.049306 |
| ZFAND3 | 6 | 0.0015141 | 0.049306 |
| ZNF596 | 8 | 0.0015141 | 0.049306 |
| ALG8 | 11 | 0.0015247 | 0.049306 |
| DSG1 | 18 | 0.0015247 | 0.049306 |
| MIR521-1 | 19 | 0.0015247 | 0.049306 |
| MBOAT2 | 2 | 0.0015247 | 0.049306 |
| ARPC2 | 2 | 0.0015247 | 0.049306 |
| ACO1 | 9 | 0.0015247 | 0.049306 |
| ARID4A | 14 | 0.0015275 | 0.049306 |
| PLA2G4D | 15 | 0.0015275 | 0.049306 |
| ANKS1A | 6 | 0.0015294 | 0.049306 |
| TAF6 | 7 | 0.0015294 | 0.049306 |
| NOLC1 | 10 | 0.0015471 | 0.049306 |
| SYMPK | 19 | 0.0015471 | 0.049306 |
| ITGA4 | 2 | 0.0015471 | 0.049306 |
| USF3 | 3 | 0.0015471 | 0.049306 |
| APBB2 | 4 | 0.0015471 | 0.049306 |
| PDLIM5 | 4 | 0.0015471 | 0.049306 |
| ATP2A1 | 16 | 0.0015573 | 0.049306 |
| LRRC36 | 16 | 0.0015573 | 0.049306 |
| TAF4B | 18 | 0.0015573 | 0.049306 |
| SLC15A2 | 3 | 0.0015573 | 0.049306 |
| SH3RF1 | 4 | 0.0015573 | 0.049306 |
| CHTOP | 1 | 0.0015587 | 0.049306 |
| ADAM11 | 17 | 0.0015587 | 0.049306 |
| HLTF | 3 | 0.0015587 | 0.049306 |
| INTU | 4 | 0.0015587 | 0.049306 |
| MYBBP1A | 17 | 0.0015668 | 0.049306 |
| SKAP2 | 7 | 0.0015668 | 0.049306 |
| TSC1 | 9 | 0.0015668 | 0.049306 |
| SSH2 | 17 | 0.0015678 | 0.049306 |
| CD53 | 1 | 0.001573 | 0.049306 |
| S100A10 | 1 | 0.001573 | 0.049306 |
| NANOS3 | 19 | 0.001573 | 0.049306 |
| MALSU1 | 7 | 0.001573 | 0.049306 |
| DEFB106B | 8 | 0.001573 | 0.049306 |
| FBXO11 | 2 | 0.0015736 | 0.049306 |
| CCDC157 | 22 | 0.0015736 | 0.049306 |
| PARVG | 22 | 0.0015736 | 0.049306 |
| FXR1 | 3 | 0.0015736 | 0.049306 |
| CCDC81 | 11 | 0.0015746 | 0.049306 |
| ARHGAP32 | 11 | 0.0015746 | 0.049306 |
| ILDR1 | 3 | 0.0015746 | 0.049306 |
| GSR | 8 | 0.0015746 | 0.049306 |
| NUP205 | 7 | 0.0015782 | 0.049374 |
| GFRA3 | 5 | 0.0015933 | 0.049797 |
| MAP1LC3B2 | 12 | 0.0015989 | 0.049878 |

| | | | |
|---|---|---|---|
| RSPH3 | 6 | 0.0015989 | 0.049878 |

**Table S1: Gastric Cancer Genes with Sexually Dimorphic Mutation Patterns**: The table lists 1,052 genes exhibiting sexually dimorphic mutation patterns. Each gene is presented with its gene symbol, chromosomal location, chi-square test p-value, and the p-value after Benjamini-Hochberg false discovery rate (FDR) correction. The genes are sorted by significance, with the smallest corrected p-values appearing at the top of the table.

**Table S2**: Overlapping between sexually dimorphic GC mutated genes and the KEGG cancer pathway, the KEGG gastric cancer pathway, the COSMIC gene census, and the tumor suppressor genes databases

| Combination ID/Number | C1 | C2 | C3 | C4 | C5 | C6 | C7 | C8 | C9 | C10 | C11 | C12 | C13 | C14 | C15 |
|---|---|---|---|---|---|---|---|---|---|---|---|---|---|---|---|
| 1 | ADCY7 | AXIN1 | ABI1 | AHRR | AXIN1 | ARNT | AXIN1 | AXIN1 | AXIN1 | AXIN1 | AXIN1 | AXIN1 | AXIN1 | AXIN1 | AXIN1 |
| 2 | ARNT | BRAF | AFDN | ANGPTL4 | BRAF | AXIN1 | DAPK1 | DAPK1 | CDH17 | CCNC | BRAF | E2F2 | GSK3B | CDH17 | GSK3B |
| 3 | AXIN1 | CDH17 | ARNT | ARMC10 | E2F2 | BRAF | E2F2 | E2F2 | E2F2 | CDH17 | GSK3B | GSK3B | HIF1A | GSK3B | RB1 |
| 4 | BIRC2 | E2F2 | AXIN1 | AXIN1 | GSK3B | EML4 | GSK3B | GSK3B | GSK3B | CIC | MET | RB1 | PRKCB | RB1 | |
| 5 | BRAF | GAB1 | BIRC6 | BLNK | MET | GSK3B | GSTP1 | GSTP1 | RB1 | CUX1 | RB1 | | RB1 | | |
| 6 | CAMK2D | GSK3B | BMP5 | CCNC | RB1 | HEY1 | HIF1A | HIF1A | | DICER1 | | | SUFU | | |
| 7 | CCNA2 | MET | BRAF | CDH17 | SOS2 | HIF1A | IGF1 | IGF1 | | ETV6 | | | | | |
| 8 | DAPK1 | RB1 | CBFB | CHD5 | WNT8A | IL6ST | PRKCB | PRKCB | | FAT1 | | | | | |
| 9 | E2F2 | SOS2 | CCNC | CHFR | | MET | RB1 | RB1 | | GSK3B | | | | | |
| 10 | EML4 | WNT8A | CDH17 | CIC | | NFKB2 | SUFU | SUFU | | HIF1A | | | | | |
| 11 | GSK3B | | CIC | CUX1 | | PIM1 | | | | IDH1 | | | | | |
| 12 | GSTP1 | | CLTC | DACH1 | | PRKCB | | | | NBN | | | | | |
| 13 | HEY1 | | CREB1 | DAPK1 | | RB1 | | | | NDRG1 | | | | | |
| 14 | HEYL | | CUX1 | DICER1 | | SUFU | | | | PRDM2 | | | | | |
| 15 | HIF1A | | DICER1 | DMBT1 | | | | | | PRKCB | | | | | |
| 16 | IGF1 | | EML4 | E2F2 | | | | | | RB1 | | | | | |
| 17 | IL23R | | EPHA7 | ESRP1 | | | | | | SDHA | | | | | |
| 18 | IL6ST | | ETV6 | ETV6 | | | | | | STK11 | | | | | |
| 19 | ITGA3 | | EWSR1 | FAT1 | | | | | | SUFU | | | | | |
| 20 | LAMB4 | | FAT1 | GSK3B | | | | | | TRIM24 | | | | | |
| 21 | LAMC1 | | FBXO11 | GSTP1 | | | | | | TSC1 | | | | | |
| 22 | MET | | FUBP1 | HIF1A | | | | | | WNK2 | | | | | |
| 23 | NFKB2 | | GSK3B | HLTF | | | | | | | | | | | |
| 24 | PIM1 | | HEY1 | IDH1 | | | | | | | | | | | |
| 25 | PRKCB | | HIF1A | IGF1 | | | | | | | | | | | |
| 26 | RB1 | | HOXD11 | IL17RD | | | | | | | | | | | |
| 27 | SOS2 | | IDH1 | IQGAP2 | | | | | | | | | | | |

| | | | |
|---|---|---|---|
| 28 | SUFU | IGF2BP2 | KLF10 |
| 29 | TRAF6 | IL6ST | LMNTD1 |
| 30 | WNT8A | LYN | MBD4 |
| 31 | | MET | MCPH1 |
| 32 | | MLF1 | MYBBP1A |
| 33 | | MLLT3 | MYO1A |
| 34 | | NBN | NBN |
| 35 | | NDRG1 | NDRG1 |
| 36 | | NFKB2 | OSCP1 |
| 37 | | NIN | PDCD4 |
| 38 | | NSD3 | PDGFRL |
| 39 | | OMD | PLXNC1 |
| 40 | | PICALM | PRDM2 |
| 41 | | PIM1 | PRKCB |
| 42 | | PMS1 | RB1 |
| 43 | | PRDM2 | RCHY1 |
| 44 | | PRKCB | RPS6KA2 |
| 45 | | PSIP1 | RUNX3 |
| 46 | | RAD17 | SDHA |
| 47 | | RAD21 | STK11 |
| 48 | | RB1 | SUFU |
| 49 | | ROS1 | TANK |
| 50 | | SDHA | TCF4 |
| 51 | | SETDB1 | TRIM13 |
| 52 | | STK11 | TRIM24 |
| 53 | | SUFU | TSC1 |
| 54 | | TNC | UFL1 |
| 55 | | TRIM24 | UNC5C |
| 56 | | TSC1 | VWA5A |
| 57 | | UBR5 | WNK2 |
| 58 | | WNK2 | ZIC1 |

**Table S2**: Overlapping between sexually dimorphic GC mutated genes and the KEGG cancer pathway, the KEGG gastric cancer pathway, the COSMIC gene census, and the tumor suppressor genes databases. Number of overlapping genes in square brackets: **C1**: Sexually Dimorphic GC & KEGG Cancer Pathway [30]; **C2**: Sexually Dimorphic GC & KEGG Gastric Cancer Pathway [10]; **C3**: Sexually Dimorphic GC & COSMIC Cancer Census [58]; **C4** Sexually Dimorphic GC & TSGenes [58]; **C5**: Sexually Dimorphic GC & KEGG Cancer Pathway & KEGG Gastric Cancer Pathway [8]; **C6**: Sexually Dimorphic GC & KEGG Cancer Pathway & COSMIC Cancer Census [14]; **C7**: Sexually Dimorphic GC & KEGG Cancer Pathway & TSGenes [10]; **C8** Sexually Dimorphic GC & KEGG Gastric Cancer Pathway & COSMIC Cancer Census [6]; **C9**: Sexually Dimorphic GC & KEGG Gastric Cancer Pathway & TSGenes [5]; **C10**: Sexually Dimorphic GC & COSMIC Cancer Census & TSGenes [22]; **C11**: Sexually Dimorphic GC & KEGG Cancer Pathway & KEGG Gastric Cancer Pathway & COSMIC Cancer Census [5]; **C12**: Sexually Dimorphic GC & KEGG Cancer Pathway & KEGG Gastric Cancer Pathway & TSGenes [4]; **C13**: Sexually Dimorphic GC & KEGG Cancer Pathway & COSMIC Cancer Census & TSGenes [6]; **C14**: Sexually Dimorphic GC & KEGG Gastric Cancer Pathway & COSMIC Cancer Census & TSGenes [4]; **C15**: Sexually Dimorphic GC & KEGG Cancer Pathway & KEGG Gastric Cancer Pathway & COSMIC Cancer C;ensus & TSGenes [3].

**Table S3**: Enrichment analyses of Gene Ontology terms in male and female higher mutation rate gene groups

| Cluster | ID | Description | GeneRatio | BgRatio | RichFactc | FoldEn | zScore | pvalue | p.adjust | qvalue | geneID | Count |
|---|---|---|---|---|---|---|---|---|---|---|---|---|
| Male | GO:0141005 | retrotransposon silencing by heterochromatin formation | 6/434 | 18/18986 | 0.33 | 14.58 | 8.817588 | 2.03E-06 | 0.01 | 0.01 | MAEL/PIWIL1/PIWIL4/SETDB1/MORC1/TDRD5 | 6 |
| Male | GO:0031507 | heterochromatin formation | 15/434 | 161/18986 | 0.09 | 4.08 | 5.9945058 | 4.50E-06 | 0.01 | 0.01 | RB1/BAZ2A/MAEL/UBR2/UBR5/PIWIL1/PIWIL4/SIN3A/BEND3/SETDB1/CDYL/MORC1/CBX3/TDRD5/ATF7IP | 15 |
| Male | GO:0045814 | negative regulation of gene expression, epigenetic | 16/434 | 192/18986 | 0.08 | 3.65 | 5.6352217 | 9.17E-06 | 0.01 | 0.01 | RB1/BAZ2A/MAEL/UBR2/UBR5/PIWIL1/PIWIL4/SIN3A/HDAC4/BEND3/SETDB1/CDYL/MORC1/CBX3/TDRD5/ATF7IP | 16 |
| Male | GO:0010526 | retrotransposon silencing | 7/434 | 36/18986 | 0.19 | 8.51 | 6.8948649 | 1.46E-05 | 0.01 | 0.01 | MAEL/UBR2/PIWIL1/PIWIL4/SETDB1/MORC1/TDRD5 | 7 |
| Male | GO:0032197 | retrotransposition | 7/434 | 38/18986 | 0.18 | 8.06 | 6.6616509 | 2.12E-05 | 0.02 | 0.02 | MAEL/UBR2/PIWIL1/PIWIL4/SETDB1/MORC1/TDRD5 | 7 |
| Male | GO:0032196 | transposition | 7/434 | 40/18986 | 0.18 | 7.66 | 6.4449001 | 3.02E-05 | 0.02 | 0.02 | MAEL/UBR2/PIWIL1/PIWIL4/SETDB1/MORC1/TDRD5 | 7 |
| Male | GO:0034101 | erythrocyte homeostasis | 13/434 | 150/18986 | 0.09 | 3.79 | 5.2495617 | 4.18E-05 | 0.02 | 0.02 | RB1/HIF1A/LYN/CDIN1/KMT2E/IREB2/ARNT/INPP5D/HEATR3/ARID4A/INHBA/DIAPH3/UFL1 | 13 |

| | | | | | | | | | | | | |
|---|---|---|---|---|---|---|---|---|---|---|---|---|
| Male | GO:0040029 | epigenetic regulation of gene expression | 19/434 | 295/18986 | 0.06 | 2.82 | 4.8121775 | 5.30E-05 | 0.03 | 0.02 | RB1/BAZ2A/MAEL/KMT2E/UBR2/UBR5/PIWIL1/PIWIL4/SIN3A/HDAC4/BEND3/SETDB1/ARID4A/CDYL/MORC1/CBX3/TDRD5/AXIN1/ATF7IP | 19 |
| Male | GO:0002244 | hematopoietic progenitor cell differentiation | 12/434 | 135/18986 | 0.09 | 3.89 | 5.1515644 | 6.41E-05 | 0.03 | 0.03 | LIPA/LYN/BRAF/SIN3A/WDR7/SOS2/DHX36/ANLN/SIPA1L3/PUS7/INHBA/UFL1 | 12 |
| Male | GO:0002262 | myeloid cell homeostasis | 14/434 | 183/18986 | 0.08 | 3.35 | 4.8789899 | 8.41E-05 | 0.03 | 0.03 | RB1/LIPA/HIF1A/LYN/CDIN1/KMT2E/IREB2/ARNT/INPP5D/HEATR3/ARID4A/INHBA/DIAPH3/UFL1 | 14 |
| Male | GO:0030218 | erythrocyte differentiation | 12/434 | 139/18986 | 0.09 | 3.78 | 5.025357 | 8.51E-05 | 0.03 | 0.03 | RB1/HIF1A/LYN/CDIN1/KMT2E/ARNT/INPP5D/HEATR3/ARID4A/INHBA/DIAPH3/UFL1 | 12 |
| Male | GO:1901987 | regulation of cell cycle phase transition | 25/434 | 479/18986 | 0.05 | 2.28 | 4.3506738 | 0.0001194 | 0.04 | 0.04 | RB1/MTBP/PSME3/RAD21/LYN/ATAD5/TAOK3/ANKRD17/KMT2E/SYCP2/SIN3A/TTK/SMARCC2/INTS3/ANLN/PPP1R10/APBB2/CLSPN/INHBA/UFL1/MBTPS1/RRM2/ATP2B4/USP28/CHFR | 25 |
| Male | GO:0006346 | DNA methylation-dependent heterochromatin formation | 7/434 | 51/18986 | 0.14 | 6.00 | 5.4734561 | 0.0001509 | 0.04 | 0.04 | BAZ2A/MAEL/PIWIL4/BEND3/SETDB1/TDRD5/ATF7IP | 7 |

| | | | | | | | | | | | | |
|---|---|---|---|---|---|---|---|---|---|---|---|---|
| Male | GO:0046885 | regulation of hormone biosynthetic process | 5/434 | 24/18986 | 0.21 | 9.11 | 6.0833952 | 0.000181 | 0.05 | 0.05 | HIF1A/LHCGR/ARNT/PDE8B/BMP5 | 5 |
| Female | GO:2000781 | positive regulation of double-strand break repair | 12/505 | 94/18986 | 0.13 | 4.80 | 6.1043444 | 7.26E-06 | 0.03 | 0.03 | ZCWPW1/PELI1/NBN/RAD51AP1/ACTR2/SMARCD3/EPC1/DPF3/TOP2B/PRMT1/UBE2V2/PHF10 | 12 |
| Female | GO:0045739 | positive regulation of DNA repair | 14/505 | 138/18986 | 0.10 | 3.81 | 5.4844527 | 1.92E-05 | 0.04 | 0.04 | ZCWPW1/PELI1/NBN/INO80C/RAD51AP1/ACTR2/SMARCD3/EPC1/DPF3/TOP2B/PRMT1/UBE2V2/PHF10/ABRAXAS1 | 14 |

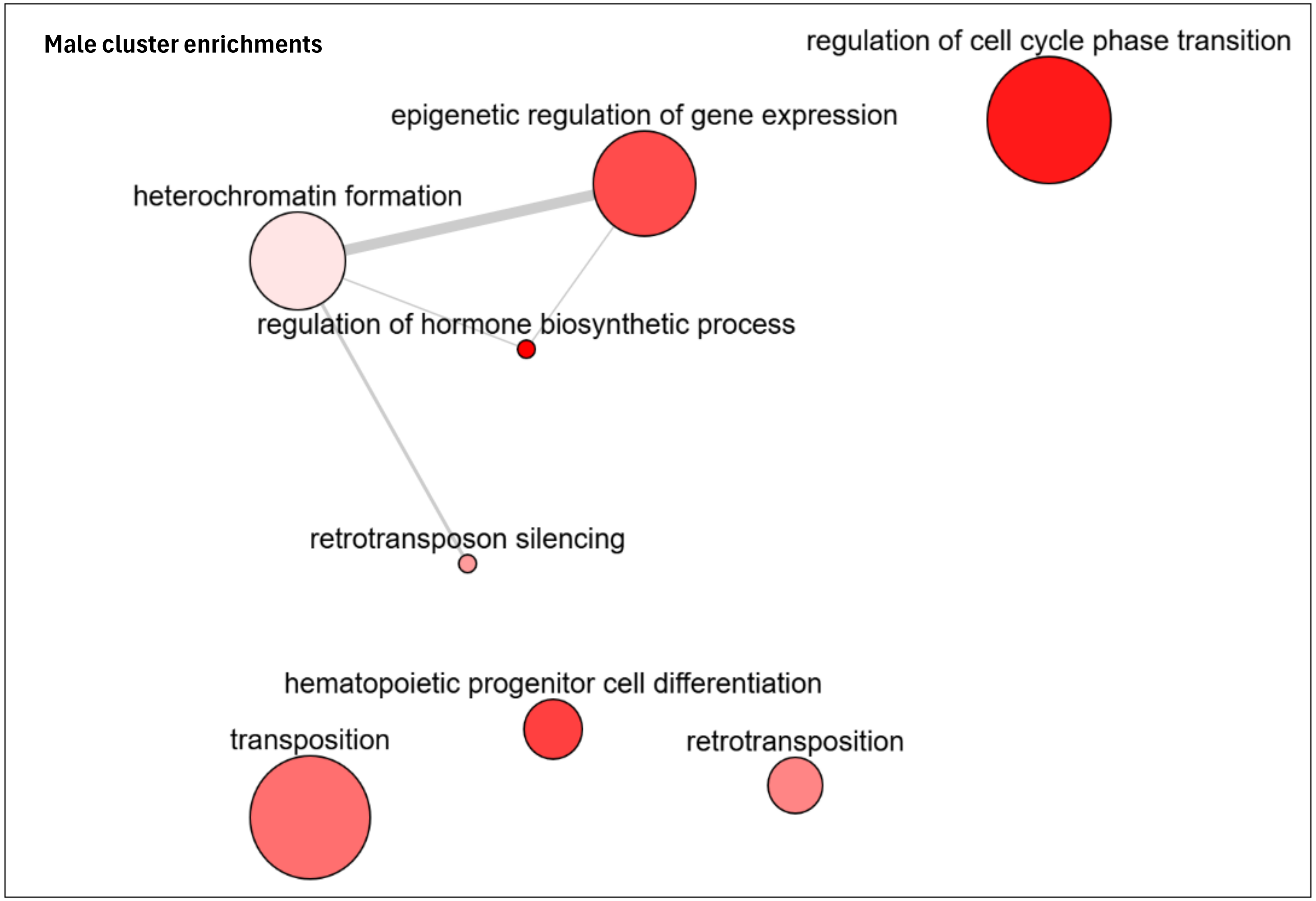

**Table S3**: Enrichment analyses of Gene Ontology terms in male and female higher mutation rate gene groups: The clusterProfiler tool was utilized to conduct enrichment analyses of Gene Ontology (GO). Enriched terms are presented by term ID, description, number and identity of genes in the cluster, and enrichment p-value. Male cluster enrichment terms connections are presented in the above plot, as illustrated by ReviGO tool (http://revigo.irb.hr/)

**Table _S4**: Top Ten Co-Mutated Gene Combinations in Males and Females

| | |
|---|---|
| **Two gene combination** | |
| **Male** | **Female** |
| TNC , KALRN [111] [90] | TNC , KALRN [90] [111] |
| TNC , TSPOAP1 [105] [86] | KALRN , COL12A1 [89] [91] |
| KALRN , PDZD2 [102] [82] | SDHA , KALRN [86] [99] |
| KALRN , SDHA [99] [86] | TSPOAP1 , TNC [86] [105] |
| TSPOAP1 , KALRN [97] [85] | SDHA , TSPOAP1 [86] [85] |
| KIAA1217 , KALRN [96] [76] | COL12A1 , TNC [85] [91] |
| TNC , SDHA [95] [85] | TSPOAP1 , KALRN [85] [97] |
| EFCAB6 , TNC [95] [81] | BIRC6 , KALRN [85] [83] |
| MYOM2 , TNC [94] [79] | TNC , SDHA [85] [95] |
| TNC , PDZD2 [94] [78] | TNC , GOLGA3 [84] [91] |
| **Three gene combination** | |
| **Male** | **Female** |
| KALRN, PDZD2, TNC [75] [65] | COL12A1, KALRN, TNC [75] [69] |
| KALRN, KIAA1217, TNC [74] [65] | COL12A1, KALRN, UTP20 [73] [64] |
| KALRN, MYOM2, TNC [73] [65] | COL12A1, KALRN, TSPOAP1 [71] [56] |
| EFCAB6, KALRN, TNC [73] [66] | COL12A1, FAT1, KALRN [71] [65] |
| KALRN, TEP1, TNC [72] [65] | BIRC6, KALRN, TNC [71] [61] |
| KALRN, TNC, TSPOAP1 [71] [69] | KALRN, TNC, UTP20 [70] [69] |
| KALRN, PDZD2, UTP20 [70] [64] | BIRC6, COL12A1, KALRN [70] [59] |
| KALRN, KIAA1217, UTP20 [70] [67] | KALRN, TNC, TSPOAP1 [69] [71] |
| KALRN, TNC, UTP20 [69] [70] | KALRN, SDHA, TNC [69] [69] |
| KALRN, SDHA, TNC [69] [69] | KALRN, MYO3B, TNC [69] [62] |
| **Four gene combination** | |
| **Male** | **Female** |
| TNC, KALRN, PDZD2, MYOM2 [58] [52] | TNC, KALRN, COL12A1, UTP20 [63] [51] |
| TNC, KALRN, KIAA1217, TEP1 [58] [52] | KALRN, COL12A1, UTP20, FAT1 [63] [51] |
| KALRN, KIAA1217, UTP20, DOP1B [57] [53] | KALRN, COL12A1, BIRC6, UTP20 [63] [48] |
| KALRN, KIAA1217, CUX1, DOP1B [57] [52] | TNC, KALRN, COL12A1, UBR2 [62] [46] |
| TNC, KALRN, MYOM2, EFCAB6 [56] [51] | TNC, KALRN, COL12A1, MYO3B [62] [47] |
| TNC, KALRN, KIAA1217, UTP20 [56] [59] | TNC, KALRN, COL12A1, BIRC6 [62] [46] |
| TNC, KALRN, COL12A1, KIAA1217 [56] [56] | KALRN, COL12A1, UTP20, ABCC4 [62] [45] |
| KALRN, PDZD2, KIAA1217, DOP1B [56] [47] | TNC, KALRN, TSPOAP1, COL12A1 [61] [43] |
| KALRN, KIAA1217, UTP20, NUP205 [56] [56] | TNC, KALRN, COL12A1, FAT1 [61] [53] |
| KALRN, COL12A1, KIAA1217, UTP20 [56] [58] | KALRN, COL12A1, VPS13D, UTP20 [61] [52] |

| Five gene combination | |
|---|---|
| **Male** | **Female** |
| **KALRN**, **PDZD2**, **KIAA1217**, **UTP20**, **DOP1B** [51] **[43]** | **KALRN**, **COL12A1**, **UTP20**, **ABCC4**, **PRDM2** [57] **[39]** |
| **KALRN**, **PDZD2**, **KIAA1217**, **CUX1**, **DOP1B** [50] **[45]** | **KALRN**, **COL12A1**, **UTP20**, **ABCC4**, **CUX1** [57] **[40]** |
| **KALRN**, **KIAA1217**, **UTP20**, **CUX1**, **DOP1B** [50] **[48]** | **KALRN**, **COL12A1**, **BIRC6**, **VPS13D**, **BTAF1** [57] **[38]** |
| **KALRN**, **COL12A1**, **KIAA1217**, **UTP20**, **NUP205** [50] **[50]** | **TNC**, **KALRN**, **COL12A1**, **UTP20**, **FAT1** [56] **[42]** |
| **KALRN**, **PDZD2**, **UTP20**, **MYH15**, **ATRN** [49] **[46]** | **TNC**, **KALRN**, **COL12A1**, **BIRC6**, **UTP20** [56] **[40]** |
| **KALRN**, **PDZD2**, **KIAA1217**, **UTP20**, **NUP205** [49] **[49]** | **KALRN**, **UTP20**, **ABCC4**, **CUX1**, **PRDM2** [56] **[42]** |
| **KALRN**, **KIAA1217**, **UTP20**, **NUP205**, **DOP1B** [49] **[46]** | **KALRN**, **COL12A1**, **UTP20**, **FAT1**, **TEP1** [56] **[41]** |
| **KALRN**, **KIAA1217**, **UTP20**, **DOP1B**, **RPS6KA2** [49] **[45]** | **KALRN**, **COL12A1**, **UTP20**, **FAT1**, **PRDM2** [56] **[41]** |
| **KALRN**, **KIAA1217**, **MYO3B**, **CUX1**, **DOP1B** [49] **[47]** | **KALRN**, **COL12A1**, **UTP20**, **FAT1**, **ABCC4** [56] **[38]** |
| **KALRN**, **KIAA1217**, **CUX1**, **DOP1B**, **UBR2** [49] **[46]** | **KALRN**, **COL12A1**, **UTP20**, **CUX1**, **PRDM2** [56] **[41]** |

**Table S4**: This table presents the top ten co-mutated gene combinations for males and females, including two-gene, three-gene, four-gene, and five-gene combinations. The count of each combination is displayed in square brackets. The **first set of brackets** contains the count of each combination in the respective sex (numbers in black). The **second set of brackets** shows the count of the same combination in the opposite sex, with numbers in **blue** for males and **purple** for females. If a given combination is ranked among the top ten in both sexes, the count in the second brackets appears in black.
